# Investigating the conformational response of the Sortilin receptor upon binding endogenous peptide- and protein ligands by HDX-MS

**DOI:** 10.1101/415620

**Authors:** Esben Trabjerg, Nadia Abu-Asad, Ziqian Wan, Fredrik Kartberg, Søren Christensen, Kasper D. Rand

## Abstract

Sortilin is a multifunctional transmembrane neuronal receptor involved in sorting of neurotrophic factors and apoptosis signalling. So far, structural characterization of Sortilin and its endogenous ligands has been limited to crystallographic studies of Sortilin in complex with the neuropeptide Neurotensin. Here, we use hydrogen/deuterium exchange mass spectrometry to investigate the conformational response of Sortilin to binding biological ligands including the peptides Neurotensin and the Sortilin propeptide and the proteins Progranulin and pro-Nerve growth factor-β. The results show that the ligands employ two binding sites inside the cavity of the β-propeller of Sortilin. However, ligands have distinct differences in their conformational impact on the receptor. Interestingly, the protein ligands induce conformational stabilization in a remote membrane-proximal domain, hinting at an unknown conformational link between the ligand binding region and this membrane-proximal region of Sortilin. Our findings improves our molecular understanding of Sortilin and how it mediates diverse ligand-dependent functions important in neurobiology.

## Introduction

The Sortilin receptor, also known as the Neurotensin receptor-3, is a multifunctional transmembrane receptor that predominantly is expressed in the central and peripheral nerve system, where it is involved in endocytosis and sorting of neurotrophic factors and apoptosis signalling (Nykjaer & Willnow, 2012). Sortilin was discovered in a biochemical screen for orphan endocytic receptors in the human brain and simultaneous described as a receptor for the neuropeptide Neurotensin (NT) (Mazella et al., 1998; C. M. Petersen et al., 1997; Zsürger, Mazella, & Vincent, 1994). The receptor consist of an N-terminal extracellular domain, a single transmembrane helix and a cytoplasmic tail. To date, four other human protein homologues have been identified; SorLA, SorCS1, SorCS2 and SorCS3 (Nielsen et al., 2001; Nykjaer & Willnow, 2012).

Sortilin is expressed in a pro-form and its 44 residue propeptide is thought to serve at least two purposes. Firstly, as an intrinsic chaperone to facilitate the correct folding of the extracellular part. Secondly, to inhibit the interaction of Sortilin with its endogenous ligands until final maturation of the receptor by the protease Furin in the Golgi compartment (Nykjaer & Willnow, 2012; C. Munck Petersen et al., 1999; Quistgaard et al., 2009; Westergaard et al., 2004).

X-ray crystal structures of the extracellular domain of Sortilin in complex with NT and small molecule NT binding inhibitors have been solved (J.L. Andersen et al., 2017; Jacob Lauwring Andersen et al., 2014; Quistgaard et al., 2014, 2009; Schrøder et al., 2014). The extracellular part of Sortilin adopts a donut-shaped fold, where the N-terminal domain forms an oval 10-bladed β-propeller with a slightly conical cavity, flanked by two minor cysteine-rich domains termed the 10CC-a and 10CC-b domains (**Fig. 1a**) (Quistgaard et al., 2009). The primary interaction site of NT was revealed in the first published crystal structure of Sortilin, where the four C-terminal residues of NT form a β-strand extension to strand 1 of blade 6 inside the tunnel of the β-propeller (**Fig. 1a**). Furthermore, hydrophobic and polar interactions with surrounding residues and the formation of a salt bridge between the C-terminal carboxylate of NT and the side chain of Arg292 constitutes the interaction interface of Sortilin and the C-terminal of NT. This interaction site has been referred to as NT interaction site 1 (NTIS1) (Quistgaard et al., 2014). Later, a second crystal structure of Sortilin in complex with NT at elevated concentration (15 times molar excess) was published (Quistgaard et al., 2014). Here, further electron density for a five-residue peptide backbone was observed on the opposite side of the β-propeller cavity forming a short β-strand extension to strand 1 of blade 1 (**Fig. 1a**). Supported by biochemical binding data of NT and fragments thereof, the authors proposed NT binding to Sortilin to be a 1:1 interaction and that the electron density found on the opposing side of NTIS1 inside the β-propeller cavity represented the N-terminal of NT (Quistgaard et al., 2014). This putative interaction site has been referred to as NT interaction site 2 (NTIS2) (Quistgaard et al., 2014).

**Figure 1.**
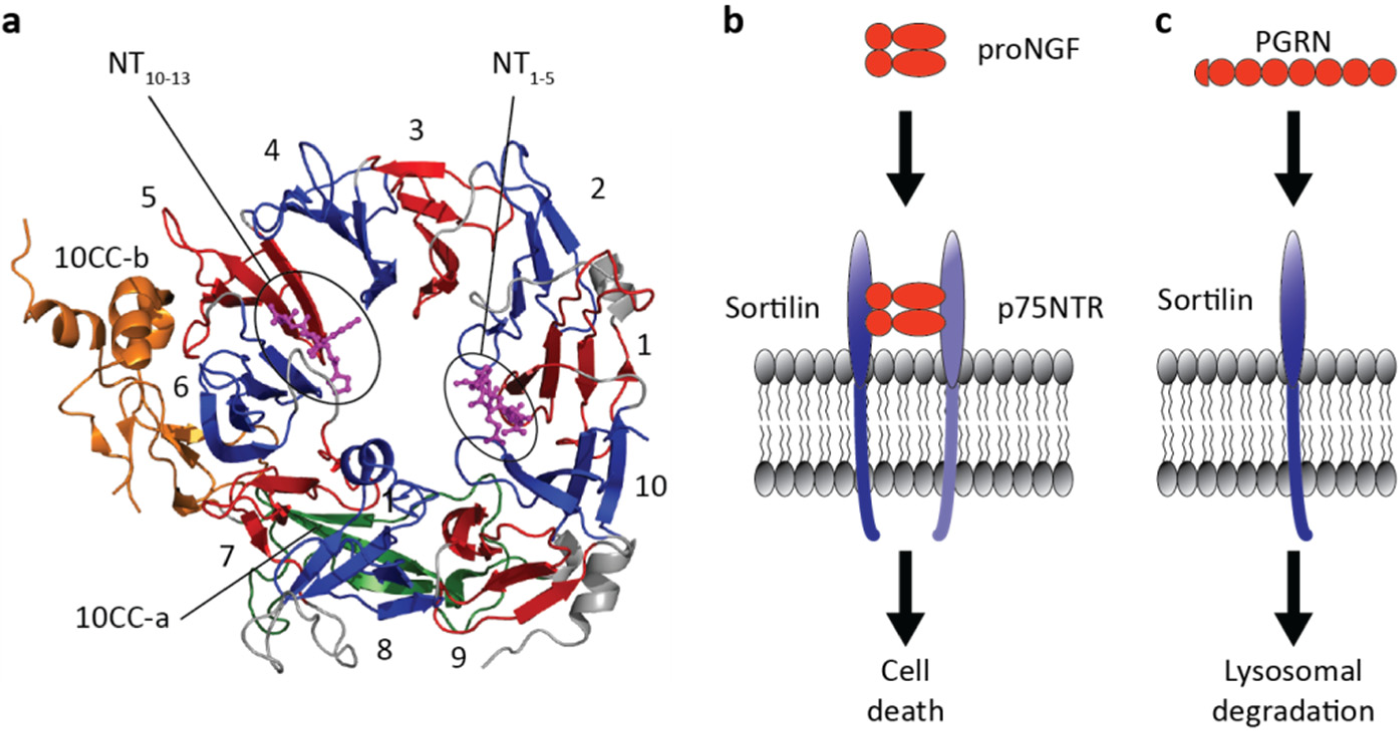
The multifunctional Sortilin receptor. (**a**) Crystal structure of Sortilin in complex with NT (PDB ID: 4PO7). The 10 blades forming the β-propeller are colored blue and red, while the the 10CC domains (a and b) are colored green and orange, respectively. NT is colored magenta, and NTIS1 is shown occupied by NT_10-13_ and the tentative NTIS2 is shown occupied by NT_1-5_. (**b**) proNGF induces apoptosis of neurons by interacting with a receptor complex of Sortilin and p75NTR. (**c**) Sortilin mediates endocytosis of PGRN and directs it for lysosomal degradation. The interaction of Sortilin with both proNGF and PGRN have been proposed as drug targets for treating various conditions of neurodegenerative diseases.

NT and the propeptide of Sortilin are only two out of many biologically relevant Sortilin ligands that include Proneurotrophins (Nykjaer, Lee, Teng, & Jansen, 2004; Teng et al., 2005), Progranulin (PGRN) (Hu et al., 2010), Lipoprotein Lipase (Nielsen, Jacobsen, Olivecrona, Gliemann, & Petersen, 1999), Apo Lipoproteins (Carlo et al., 2013; Nielsen et al., 1999), Amyloid Precursor Protein (Gustafsen et al., 2013), Aβ (Carlo et al., 2013) and the receptor associated protein (C. M. Petersen et al., 1997). Lately, the interaction of Sortilin with both the pro-form of Nerve growth factor-β (proNGF) and PGRN has received increased attention as these interactions have been proposed as putative drug targets for treating neurodegenerative diseases (**Fig. 1b,c**) (Kao, Mckay, Singh, Brunet, & Huang, 2017; Lee et al., 2014; Nykjaer & Willnow, 2012).

proNGF is a dimer at native conditions with a well-defined mature part and a largely disordered pro-part consisting of 103 residues (Feng et al., 2010; Esben Trabjerg, Kartberg, Christensen, & Rand, 2017). proNGF induces apoptosis of neurons by binding to a receptor complex consisting of Sortilin and the common neurotrophine receptor (p75NTR) (**Fig. 1b**) (Nykjaer et al., 2004). Furthermore, increased levels of proNGF have been associated with pathological conditions where apoptosis of neurons are prevalent, e.g. Alzheimer’s disease, Multiple sclerosis, ischemic stroke and seizure (Beattie et al., 2002; Fahnestock, Michalski, Xu, & Coughlin, 2001; Jansen et al., 2007; Lowry et al., 2001; Perez et al., 2015).

PGRN is an extracellular secreted glycoprotein consisting of seven and a half tandem repeats of a conserved granulin fold (Bhandari, Palfree, & Bateman, 1992). The full-length structure of PGRN is unknown, but the structure of several individual granulins has been determined by nuclear magnetic resonance spectroscopy (Hrabal, Chen, James, Bennett, & Ni, 1996; Tolkatchev et al., 2008). An absence of PGRN has been linked to neurodegeneration as mutations causing PGRN deficiency results in frontotemporal dementia (Baker et al., 2006). The bona fide receptor that transduce the neurotrophic PGRN signals is still elusive, but extracellular supplementation of PGRN to PGRN-knockdown neuronal cultures has been shown to rescue neurite outgrowth (Gass et al., 2012; Kao et al., 2017). In contrast, PGRN has been shown to bind Sortilin, which targets PGRN for endocytosis and lysosomal degradation. Hence, a inhibition of this interaction has been proposed as a strategy for treating frontotemporal dementia (**Fig. 1c**) (Kao et al., 2017; Lee et al., 2014).

Insights into the conformation and dynamics of unbound Sortilin and how Sortilin binds to larger endogenous protein ligands is currently lacking. In this study we have investigated the molecular interactions of Sortilin and both small and large binding partners (NT, the Sortilin propeptide, proNGF and PGRN) by hydrogen/deuterium exchange followed by mass spectrometry (HDX-MS). HDX-MS represents a sensitive analytical approach to probe the conformational dynamics of proteins and how these change upon ligand binding. The technique can provide access to large and challenging protein systems at native solution-phase conditions with very little sample consumption (Hvidt & Linderstrom-Lang, 1955; Hvidt & Linderstrøm-Lang, 1954; Hvidt & Nielsen, 1966; Jensen & Rand, 2016; Konermann, Pan, & Liu, 2011; Esben Trabjerg, Nazari, & Rand, 2018; Wales & Engen, 2006). During an HDX-MS experiment the exchange rate of backbone amide hydrogens is measured, which is primarily dependent on the presence and stability of local hydrogen bonding (McAllister & Konermann, 2015; Persson & Halle, 2015; Skinner, Lim, Bédard, Black, & Englander, 2012a, 2012b). Hence, the exchange rate is highly informative on protein higher-order structure and can vary up to eight orders of magnitude, when comparing backbone amides in disordered regions (non-hydrogen bonded) to backbone amides in heavily structured regions (e.g. stable α-helices or β-sheets) (Englander & Kallenbach, 1983). HDX-MS is thus very suitable for detecting subtle changes in protein conformation and dynamics caused by changes to the solution environment or due to specific molecular interactions (Leurs et al., 2014; Morgan & Engen, 2009).

Our HDX-MS results indicate that Sortilin employs two binding surfaces in the cavity of the β- propeller to bind its diverse biological ligands. The conformational imprint of each ligand on the receptor, however, has unique features, and in the case of proNGF and PGRN further involves the remote 10CC domains. The observed distinct changes in dynamics of Sortilin upon ligand binding could be important for function and in mediating diverse biological downstream signalling.

## Results

### Conformational dynamics of the unbound Sortilin Receptor

The extracellular ligand-binding domain of Sortilin accounts for 89% of the total sequence of the receptor and is a highly glycosylated and disulfide-bonded domain, with six N-linked glycosylation sites and a total of eight disulfide bonds (C. M. Petersen et al., 1997; C. Munck Petersen et al., 1999). HDX-MS analysis of Sortilin was performed with an effective sequence coverage of 96% of the extracellular domain of Sortilin domain following significant optimization of conditions for protein digestion (quench buffers) as well as identification of glycopeptides from a glycan analysis of the protein (**Supplementary Table 1** and **Supplementary Fig. 1**).

Solution-phase HDX of unbound Sortilin revealed both well-defined structural segments and highly dynamic regions. In general, the backbone amides in the strands of each β-blade of the β- propeller structure of Sortilin incorporated deuterium slowly, while loops connecting the single β- blades exhibited faster deuterium uptake, in good overall correlation to the ligand-bound crystal structure (**Supplementary Fig. 2** and **Supplementary HDX plots**). One striking discrepancy between the crystal structure of ligand-bound Sortilin and the solution-phase dynamics of unbound Sortilin was observed in the three C-terminal α-helices (residues 664-671, 673-678 and 702-709) of the crystal structure. These segments exhibited fast HDX kinetics with nearly full deuterium incorporation already after 15 seconds (**Supplementary Fig. 2** and **Supplementary HDX plots**). This is highly unusual for secondary structure elements, where a protection from HDX caused by the presence of hydrogen bonds is normally observed. Hence, our results strongly suggest that the three C-terminal α-helices are highly dynamic or unstructured in solution.

**Figure 2.**
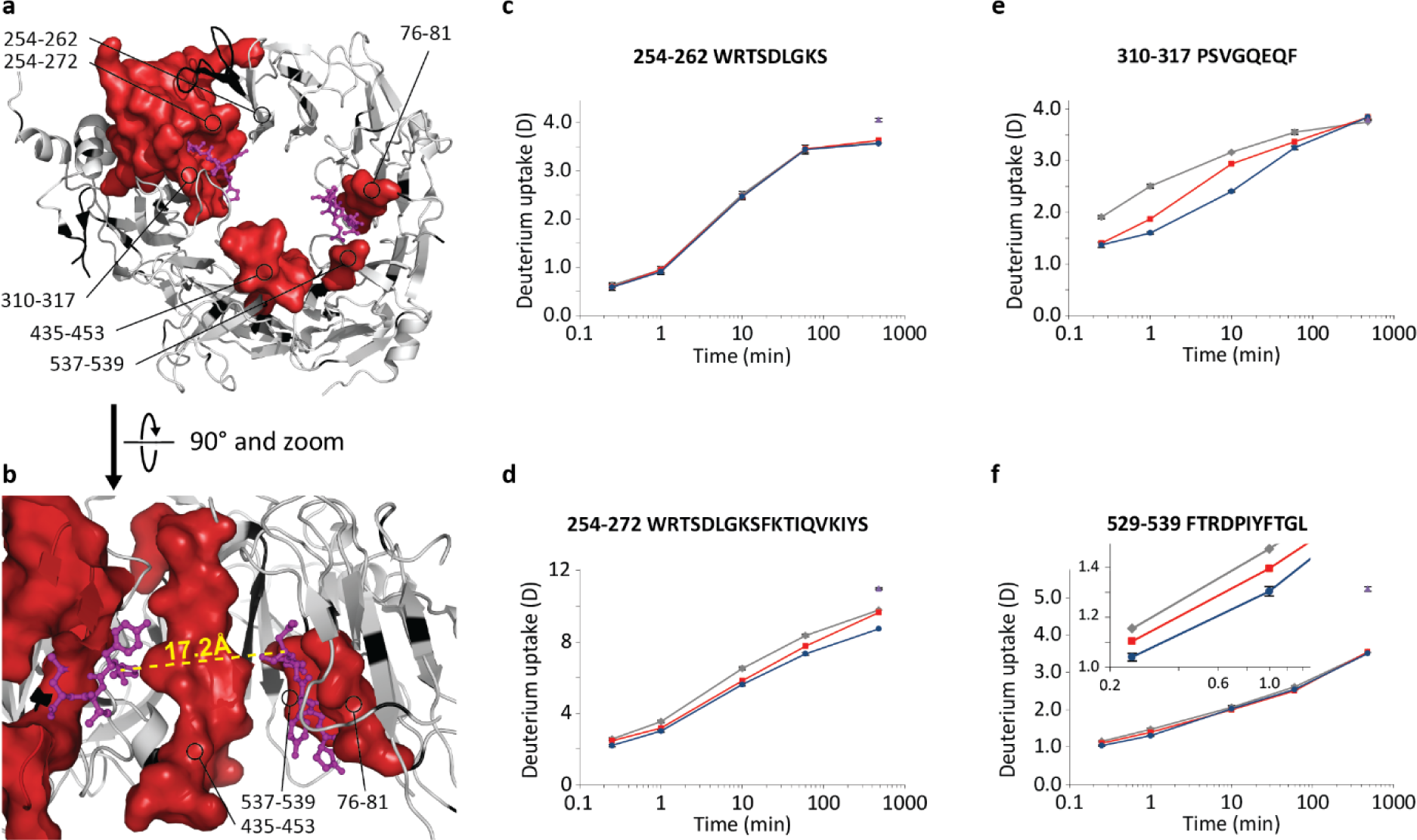
Mapping the conformational stabilization of Sortilin upon NT binding. (**a,b**) Differential HDX results are mapped onto the crystal structure of Sortilin (PDB ID: 4PO7). Regions stabilized in the presence of NT are coloured red. Regions showing no change in HDX are coloured white, while regions without coverage are coloured black. The C- and N-terminal of NT are shown in magenta. The dashed yellow line in (**b**) marks the Euclidean distance on 17.2 Å between the C-α of Asn5 and Pro10 in NT. (**c-f**) HDX plots of selected peptides. Absolute deuterium incorporation is plotted as a function of time for Sortilin (grey lines), Sortilin in the presence of 1.5 times molar excess of NT (red lines) and Sortilin in presence of 15 times molar excess of NT (blue lines). Equilibrium labelled (90%) Sortilin control samples are plotted as filled purple triangles at the 8 h time point. SD is plotted as error bars (are only slightly visible). (For Sortilin and Sortilin in presence of 15 times molar excess of NT: n=3 for all time points and the equilibrium labelled sample, except the 8 h time point where n=1. For Sortilin in presence of 1.5 times molar excess of NT: n=1 for all time points).

### Conformational impact of Neurotensin binding

The neurotransmitter NT was the first identified peptide ligand of Sortilin. To investigate the conformational response of Sortilin to NT binding, we measured the HDX of Sortilin in presence and absence of NT. Upon ligation with NT, a decrease in HDX was observed in peptides covering a large area in and around NTIS1 (**Fig. 2** and **Supplementary HDX plots**).

The conformational impact of NT binding was not confined to residues in NTIS1, but also peptides spanning areas on the other side of the β-barrel cavity in the N-terminal end of β-blade 1 and in β-blade 10 (residues 76-81 and 537-539, respectively) showed a significant reduction in HDX in the presence of NT. This area overlaps with the proposed secondary interaction site of NT, where residues 77-81 engage in hydrogens bonds with the N-terminal residues of NT. Residues 537-539 have not previously been implicated in NT binding, however they are in close proximity to the N-terminal of NT and specifically the backbone amide of Leu539 is in close proximity to the end of the side chain of Glu4 of NT. Furthermore, locally confined reduction in HDX was observed on the top and inside the cavity of the β-propeller in the middle of blade 8 (residues 435-453) located between NTIS1 and the putative NTIS2, indicating a possible third interaction point between NT and Sortilin.

To further dissect the binding mode of NT we mapped the conformational response of Sortilin in the presence of a small molecule Sortilin inhibitor (AF38469) known to emulate the binding mechanism of the C-terminal part of NT (Schrøder et al., 2014). Here, a protection from HDX was observed exclusively for peptides spanning the core of NTIS1 (**Supplementary Fig. 3**) and even more outlying regions of NTIS1 that were perturbed by NT did not show reduced HDX upon AF38469 binding (residues 271-281 and 319-321).

**Figure 3.**
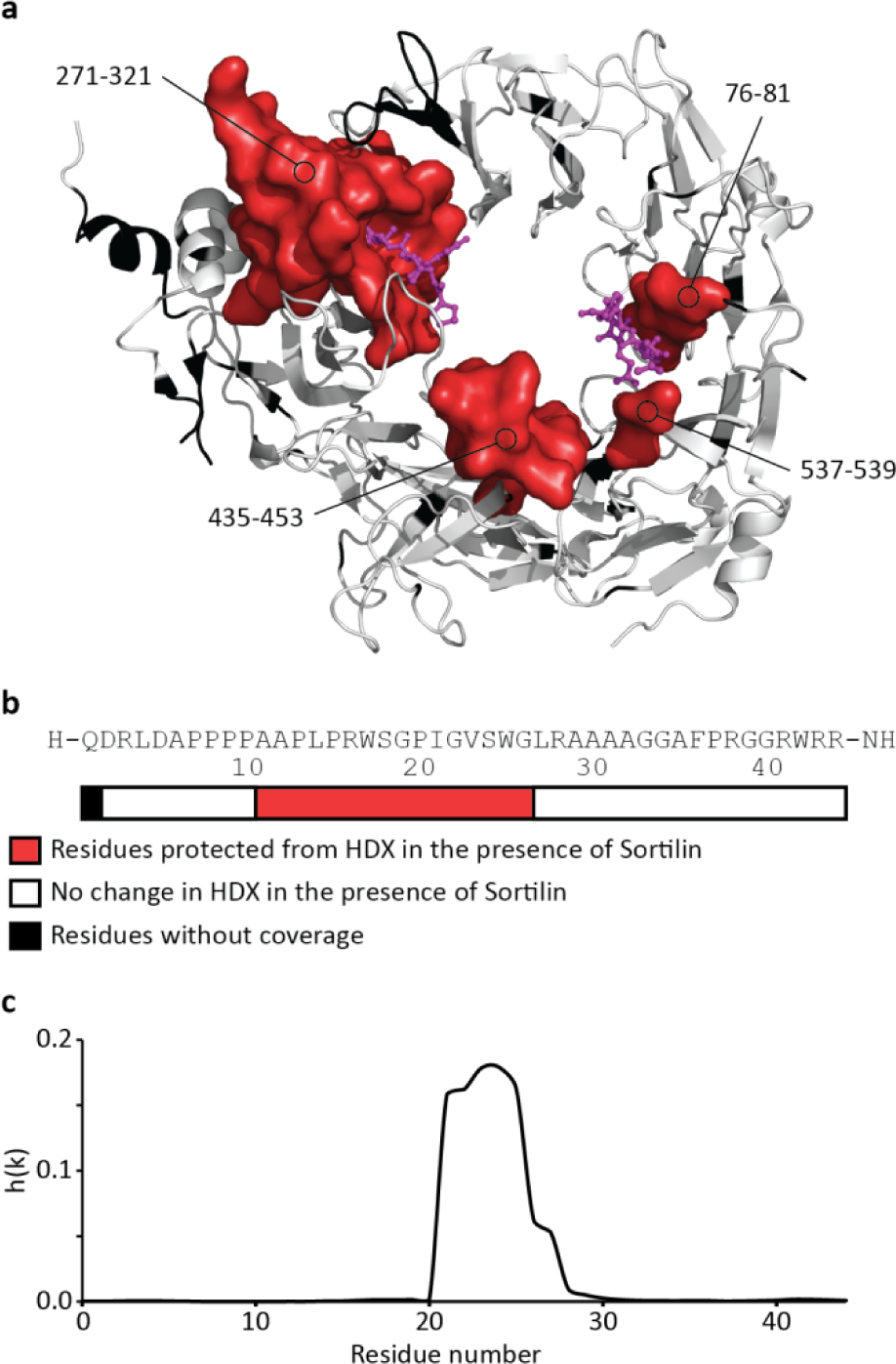
Mapping the conformational binding interface in the Sortilin-proSort complex. (**a**) Differential HDX results are mapped onto the crystal structure of Sortilin (PDB ID: 4PO7). Regions stabilized in the presence of proSort are coloured red. Regions showing no change in HDX are coloured white, while regions without coverage are coloured black. The C- and N-terminal of NT are shown in magenta. (**b**) The proSort sequence is shown and the bar below displays residues with reduced HDX in the presence of Sortilin (red) and unaffected residues (white). Regions without coverage are coloured black. (**c**) Prediction of β-aggregation of proSort by the PASTA server (http://protein.bio.unipd.it/pasta2/)(Walsh et al., 2014). The proSort sequence was submitted to the PASTA server, and the aggregation probability score (h(k)) was plotted as a function of the proSort sequence.

NTIS2 was revealed in a crystal structure of Sortilin in the presence of 15-times molar excess of NT. This NT concentration is probably not physiologically relevant, so we performed a HDX experiment of Sortilin in the presence of either 1.5- or 15-times molar excess of NT directly corresponding to the two published crystal structures of the NT-Sortilin complex (**Fig. 2** and **Supplementary HDX plots**). Even at the lower NT concentration, reduced HDX could be observed in peptides spanning NTIS2 and in the middle of blade 8.

### Structural characterization of Sortilin in complex with its propeptide

The propeptide of Sortilin is a 44-residue N-terminal extension that inhibits interactions between Sortilin and other binding partners. The propeptide is enzymatically cleaved off under maturation of the extracellular domain of Sortilin, hence a synthetic produced free form of the propeptide with an amidated C-terminal (proSort) was employed as a surrogate of the real N-terminal extension (Kitago et al., 2015; C. Munck Petersen et al., 1999).

The conformational impact of proSort binding to Sortilin was investigated by HDX-MS. Overall, Sortilin showed a reduction in HDX in the presence of the proSort in similar regions as upon ligation with NT (**Figs. 2** and **3**). However, while the magnitude of the observed reductions in HDX around NTIS1 were highly similar between NT and proSort, the magnitude of HDX reductions around NTIS2 were substantially more pronounced upon proSort binding relative to NT binding (**Supplementary HDX plots**).

To further elucidate the binding mechanism of the propeptide the reciprocal HDX-MS experiment was performed, where the HDX of proSort was measured in the absence and presence of Sortilin (**Fig.3b** and **Supplementary HDX plots**). In the absence of Sortilin, proSort exhibited HDX kinetics of a highly flexible unstructured polypeptide with no evidence of local higher-order structure as all regions of the peptide were fully deuterated already after 10 seconds of labelling. In the presence of Sortilin, however, a significant protection from HDX could be observed for peptides spanning the N-terminal half of proSort. These changes in HDX could be sub-localized to the segment encompassing residues 11-26 as 1) no significant change in HDX was observed for peptide 26-44 and 2) the magnitude of protection from HDX in peptide 10-44 was equal to the difference observed in intact proSort (**Supplementary HDX plots**). Interestingly, the stabilized region (residues 11-26) encompass the region of proSort that is most prone to form β-amyloid aggregation as predicted by the PASTA server (**Fig. 3c**) (Walsh, Seno, Tosatto, & Trovato, 2014).

### Mapping of the interaction sites of the Sortilin-proNGF complex

We next investigated the conformational response of Sortilin upon proNGF binding by HDX-MS. In the presence of proNGF A significant decrease in HDX could be observed in peptides of Sortilin spanning the C-terminal part of NTIS1 (residues 284-318). Compared to the NT and proSort, no protection from HDX was observed in NTIS2 or any other region inside the central cavity of the β-propeller architecture. In contrast, reduced HDX was observed in a cluster on the outer rim of the β-propeller (residues 387-395, 429-433 and 455-466) (**Fig. 4**). Furthermore, a conformational stabilization was also observed in the 10CC-b domain (residues 683-699). Interestingly, this region is on the opposite face of the β-propeller as both NTIS1 and the stabilized region on the outer rim of the β-propeller.

**Figure 4.**
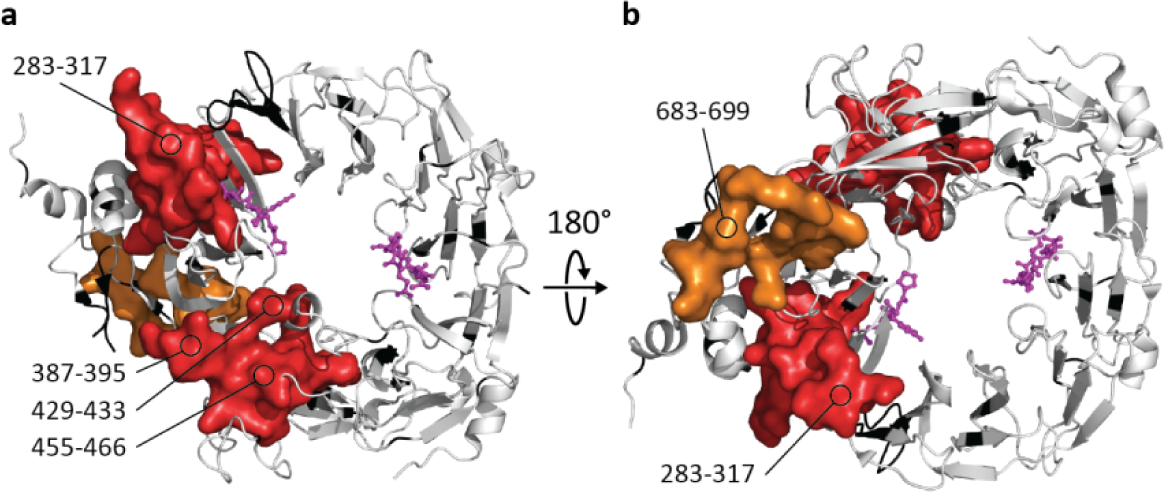
Mapping the conformational stabilization of Sortilin upon proNGF binding. (**a,b**) Differential HDX results are mapped onto the crystal structure of Sortilin (PDB ID: 4PO7). Regions stabilized in the presence of proNGF are coloured red (β-propeller domain) and orange (10CC domains). Regions showing no change in HDX are coloured white, while regions without coverage are coloured black. The C- and N-terminal of NT are shown in magenta.

To further investigate the binding mode of proNGF to Sortilin, we mapped the conformational response of Sortilin upon binding of GSTpro, a protein construct where the pro-part of proNGF is N-terminally fused with Glutathione-S transferase (GSTpro), and which binds with similar affinity to Sortilin as proNGF (Nykjaer et al., 2004). GSTpro binding to Sortilin closely emulated the conformational response of Sortilin upon proNGF binding, as a decrease in HDX was observed in peptides spanning both NTIS1, the outer rim cluster of the β-propeller (residues 386-395, 428-434 and 455-466) and in the 10CC domain (residues 682-699) (**Supplementary Fig. 4** and **Supplementary HDX plots**). As a control experiment, the HDX of Sortilin in the presence of Nerve growth factor-β (the mature part of proNGF) and Glutathione-S transferase was also measured. In both cases, no significant changes in deuterium uptake could be detected (**Supplementary Fig. 4** and **Supplementary HDX plots**).

The conformational impact of Sortilin binding to proNGF and GSTpro was also investigated by measuring the HDX of both in the presence of a molar excess of Sortilin. However, no changes in HDX were observed in the pro-part of either proNGF or in GSTpro (data not shown).

### Mapping of the interaction sites of the Sortilin-PGRN complex

The conformational response of Sortilin to binding of PGRN was investigated by HDX-MS. The impact of PGRN binding shared similarities with NT and proSort binding but also significant differences (**Fig. 5** and **Supplementary HDX plots**). A significant reduction in HDX was detected for peptides in and around NTIS1. However, a larger area around NTIS1 was affected than seen for proSort and NT. The conformational response was not limited to NTIS1 as peptides in NTIS2 also showed a structural stabilization in the presence of PGRN. As seen for proSort the magnitude of the reduction in NTIS2 was larger than seen for NT binding. Furthermore, a large surface on the outer rim of the β-barrel of Sortilin, showed a significant reduction in HDX (residues 362-370, 387-395, 435-453 and 455-466). Additionally, a reduction in HDX was observed in the 10CC-b domain (residues 641-644 and 683-695). Interestingly, structural stabilization was also observed in this region in presence of proNGF.

**Figure 5.**
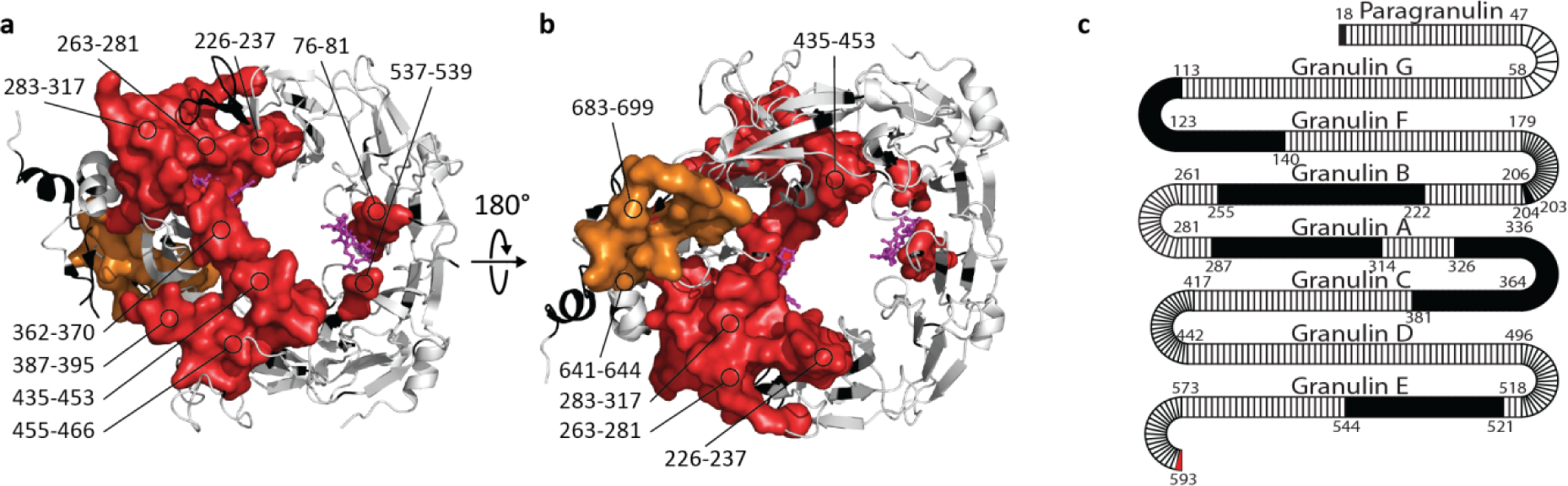
Mapping the conformational stabilization of Sortilin upon PGRN binding. (**a,b**) Differential HDX results are mapped onto the crystal structure of Sortilin (PDB ID: 4PO7). Regions stabilized in the presence of PGRN are coloured red (β-propeller domain) and orange (10CC domains). Regions showing no change in HDX are coloured white, while regions without coverage are coloured black. The C- and N-terminal of NT are shown as sticks in magenta. (**c**) Differential HDX resuls are mapped onto a model representation of PGRN. Regions stabilized in the presence of Sortilin are coloured red, regions showing no change in HDX are coloured white, while regions without coverage are coloured black. Granulin domains (Paragranulin, G, F, B, A, C, D and E) are depicted as horizontal segments, while the connecting linker regions are represented as loops.

We also performed the reciprocal HDX-MS experiment, where the binding impact of Sortilin on PGRN was investigated. To obtain an adequate sequence coverage of PGRN, due to the presence of multiple highly stable disulfide bonds, on-line electrochemical reduction was applied during the HDX-MS workflow (**Supplementary Fig. 5**) (Mysling, Salbo, Ploug, & Jørgensen, 2014; Esben Trabjerg et al., 2015). To our knowledge this is the first time this approach has been employed to facilitate HDX-MS analysis of a large protein complex. Strikingly, the conformational impact of Sortilin on PGRN was restricted to the very last C-terminal residue, as only peptides spanning the last residue of PGRN showed a significant protection from exchange in presence of Sortilin (**Fig. 5c** and **Supplementary HDX plots**).

## Discussion

### Peptide ligands employ two distinct binding sites in the cavity of the β-barrel of Sortilin

NT binding to Sortilin resulted in significant reductions in HDX for residues in both NTIS1 and NTIS2 and a patch of residues on the top and inside the cavity of the β-propeller between NTIS1 and NTIS2 (residues 435-453) (**Fig. 2**). The observed reduction in HDX in NTIS1 correlate very well with published crystal structures of the NT-Sortilin complex, as all residues forming either polar or Van der Waals contacts with NT showed reduced HDX and thus conformational stabilization in the presence of NT (Quistgaard et al., 2014, 2009).

The magnitude of the reduced HDX observed in NTIS2 are markedly less than the reductions in HDX observed in NTIS1, which could suggest that the observed reduction of HDX in NTIS2 represent an indirect effect of NT binding. From HDX data alone, it is not straightforward to discern whether a protection in HDX at a specific site is caused by a direct interaction between two binding partners or are an indirect effect of binding (allosteric) (Wales & Engen, 2006). However, the mapping of the conformational response of Sortilin to AF38469 showed that AF38469 binding to Sortilin only reduced HDX in residues of NTIS1 (**Supplementary Fig. 3**). Initially, NTIS2 was revealed in a crystal structure of Sortilin in the presence of 15-times molar excess of NT. The presented HDX results showed a reduction in HDX in NTIS2 even at 1.5-times molar excess of NT discarding the concern of crystallization artefacts (**Fig. 2** and **Supplementary HDX plots**) (Quistgaard et al., 2014). Our results thus strongly suggest that the reduced HDX in NTIS2 upon NT binding is due to a direct interaction with the N-terminal of NT as proposed by Quistgaard et al.

Recently, a crystal structure of a Sortilin homologue (SorLA) in complex with fragments of peptide ligands has been published (Kitago et al., 2015). These ligands bind in a binding site similar to NTIS2, where the interaction between the peptide ligands and SorLA mainly is mediated by the peptide backbone (**Supplementary Fig. 6**). The reduced HDX observed in the cavity of the β-propeller in the middle of blade 8 (residues 435-453) has not been described before. Interestingly, the segment of SorLA corresponding to residues 433-444 in Sortilin forms an unstructured loop in the unbound crystal structure of SorLA, but is essential for peptide binding to SorLA and adopts structure in the presence peptide ligands (**Supplementary Fig. 6**). In the SorLA structure, this loop has been suggested to “push” the peptide ligands against the cavity wall of the β-propeller (Kitago et al., 2015). Based on the obtained HDX data, a similar mechanism seems to occur in Sortilin, as the majority of protected sites can be sub-localized to the first part of residues 435-453 (**Supplementary HDX plots**). The observed HDX reduction in segment 435-453 could also be due to a direct interaction between residues in this region and the NT residues not defined in the crystal structures of the Sortilin-NT complex (Quistgaard et al., 2014, 2009). However, this is highly unlikely as the distance between the Cα Asn5 and the electron density tentatively assigned to Cα of Pro10 of NT is only 17.2 Å (Quistgaard et al., 2014). Thus, the middle part of NT needs to fully extend across the cavity to connect Asn5 and Pro10, leaving insufficient conformational flexibility to bend towards and interact with segment 435-453. Hence, we propose that residues in the beginning of residues 435-453 of Sortilin folds down to interact with the N-terminal of NT, thus essentially directly participating in the NTIS2 binding site.

The conformational stabilization induced by NT and proSort on Sortilin was highly similar (**Figs. 2** and **3**). In good agreement, it has been shown that proSort binding can be outcompeted by NT, and that mutation in NTIS1 (Ser283Glu) abolishes proSort binding to Sortilin (C. Munck Petersen et al., 1999; Quistgaard et al., 2009). However, the magnitude of reduction in HDX in NTIS2 was markedly larger for proSort binding compared to NT binding (**Supplementary HDX plots**), suggesting that NTIS2 is not a low affinity modulatory interaction site for proSort, as it is for NT. Further, this is supported by the crystal structure of SorLA, where fragments of the SorLA propeptide were shown to engage in a binding site similar to NTIS2 in Sortilin (**Supplementary Fig. 6**). As the interaction between SorLA and peptide ligands was only mediated by backbone interactions Kitago et al. suggested that the binding site in SorLA is highly promiscuous and able to bind peptide ligands prone to β-amyloid aggregation. We find this suggestion to appear valid for the Sortilin-proSort interaction as well, as there is a good correlation between residues in proSort undergoing reductions in HDX upon Sortilin binding and the theoretical propensity of residues in proSort to form β-amyloid aggregates (**Fig. 3b,c**). Interestingly, Sortilin has also been reported to bind to Amyloid-β_40_, a prototype of β-amyloid forming peptides. This interaction is of moderate affinity (K_d_=0.8µM) (Carlo et al., 2013), highlighting that the interaction of proSort with NTIS1 is probably necessary for the high binding affinity of proSort to Sortilin. Furthermore, the observed reduction in HDX of proSort correlates with data in the literature, where the binding core of a similar construct was suggested to be residues 23-28 (Westergaard et al., 2004).

Overall, our results suggest that the C-terminal part of the region in proSort observing a structural stabilization in presence of Sortilin (residues 11-26) form a β-strand extension to blade 1 in the Sortilin β-propeller (NTIS2). A β-strand that is pushed against the cavity wall of the β-propeller by the N-terminal part of residues 435-453 of Sortilin. Furthermore, some of the more N-terminal residues inthe segment of proSort experiencing a reduction in HDX, when bound to Sortilin, directly engage with residues in NTIS1.

Thus, the presented HDX results of the proSort-Sortilin complex provide a molecular mechanism for how the propeptide of Sortilin shields Sortilin from binding with other known Sortilin ligands, as it specifically binds strongly to residues in both NTIS1 and NTIS2 (in contrast to NT) (C. Munck Petersen et al., 1999). In a biological context this is highly important as only a minor fraction of active Sortilin is situated on the cell surface (~10%), whereas the remainder of the receptor population is found in intracellular compartments, especially in the Golgi compartment and vesicles, where Sortilin is implicated in complex trafficking and sorting pathways (Nielsen et al., 1999; Nykjaer & Willnow, 2012). Furthermore, it has recently been proposed that binding of the propeptide to Sortilin inhibits homodimer formation of Sortilin at acidic pH in late endosomes (Januliene et al., 2017; Leloup et al., 2017). Hence, the ability of the propeptide to shield immature Sortilin from binding with its biological ligands and inhibit homodimer formation at low pH could be important to secure that these trafficking and sorting pathways are not disturbed.

### Conformational response of Sortilin upon binding protein ligands

Sortilin is capable of binding several proteins (Hu et al., 2010; Nielsen et al., 1999; Nykjaer et al., 2004;C. M. Petersen et al., 1997; Teng et al., 2005). In the present study, we have investigated the complex formation of Sortilin with proNGF and PGRN. The mapping of the binding impact of proNGF on Sortilin showed that the presence of proNGF caused conformational stabilization of Sortilin residues in the N-terminal part of NTIS1, while no conformational effects were observed in NTIS2 (**Fig. 4**). Earlier studies have shown that NT is able to outcompete proNGF binding to Sortilin (Nykjaer et al., 2004). As NTIS1 was stabilized by both proNGF and NT, and NTIS2 was unperturbed by proNGF, our results strongly indicate that NTIS1 is the primary interaction site for proNGF on Sortilin. This suggestion is in contrast to a prior study that identified a linear epitope of Sortilin residues 163-174 to be essential for proNGF binding to Sortilin (Serup Andersen et al., 2010). In the present study, no signs of conformational changes were observed in these residues for either proNGF, GSTpro or NT binding to Sortilin (**Figs. 2** and **4**, **Supplementary Fig. 4** and **Supplementary HDX plots**). It is important to note that the interaction of proNGF with Sortilin appears to be of a different molecular nature than that of proSort and NT, as the mutation Ser283Glu has been shown to abolish proSort and NT binding to Sortilin but not proNGF binding (Quistgaard et al., 2009). Hence, the ability of NT to inhibit proNGF binding to Sortilin could be steric in nature rather than a competition of binding to an identical binding pocket.

Our results demonstrate that proNGF binding impacts residues outside of NTIS1 evidenced by selective reductions in HDX in regions on the outer rim of the β-propeller (residues 428-433, 387-395 and 455-466) of Sortilin upon proNGF binding. The specific importance of region 455-466 have recently been supported by mutational studies, as proNGF has decreased affinity towards a Ala464Glu mutation of Sortilin (Leloup et al., 2017). ProNGF impacts primarily the top of the β-propeller as HDX reductions in NTIS1 of the largest magnitude can be sub-localized to residues 307-317 and the surface patch on the outer rim of the β-propeller. Interestingly, this entails that the reduction in HDX observed in the 10CC-b domain is of an indirect (allosteric) nature (**Fig. 4**). This will be discussed further in the context of PGRN binding.

The binding mode of proNGF to Sortilin was further revealed by the HDX analysis of Sortilin in the presence of NGF and GSTpro. NGF alone had no impact on Sortilin, while GSTpro binding closely emulated the impact of proNGF binding, even the allosteric effects observed in the 10CC-b domain (**Fig. 4** and **Supplementary Fig. 4**). Thus, we conclude that the conformational stabilization of Sortilin by proNGF is mediated exclusively by the propeptide of proNGF. Analysis of the HDX of either proNGF or GSTpro upon Sortilin binding showed that Sortilin did not induce significant folding or induction of stable higher-order structure in the interacting propeptide of proNGF. This is probably due to the highly flexible nature of the pro-part of proNGF (Paoletti et al., 2009, 2011; Esben Trabjerg et al., 2017). Earlier studies have also suggested that the proNGF-Sortilin interaction is mediated via the pro-part of proNGF, as GSTpro has similar affinity towards Sortilin as proNGF and is able to outcompete proNGF binding (Nykjaer et al., 2004). Hence, GSTpro has been used as a surrogate ligand for proNGF binding in several mechanistic studies investigating Sortilin biology despite the incomplete molecular understanding of either proNGF or GSTpro binding to Sortilin (Carlo et al., 2013; Domeniconi & Chao, 2008; Mortensen et al., 2014; Nykjaer et al., 2004; Serup Andersen et al., 2010). To our knowledge, this study shows for the first time that GSTpro impacts the conformation of Sortilin in a similar manner as proNGF and thus is an accurate surrogate. However, it is important to note that GSTpro is not able to induce the apoptotic effect of proNGF against neurons as simultaneous binding of proNGF to Sortilin and p75NTR (mediated by the mature part) is essential (He & Garcia, 2004; Nykjaer et al., 2004; Wehrman et al., 2007).

Prior structural studies of the PGRN-Sortilin complex have shown that NT is able to outcompete PGRN binding and that residue Ser283 in NTIS1 is essential for PGRN binding to Sortilin (Hu et al., 2010; Zheng, Brady, Meng, Mao, & Hu, 2011). This is in good agreement with the conformational response of Sortilin to PGRN binding as residues of NTIS1 showed reduced HDX in the presence of PGRN (**Fig. 5**). Furthermore, protection from HDX was also observed in NTIS2. The reduction of HDX in NTIS2 by PGRN binding was markedly more pronounced than for NT binding demonstrating a significant involvement of NTIS2 in the PGRN binding event (**Supplementary HDX plots**). In support, anti-Sortilin antibodies have recently been identified which are able to inhibit the PGRN-Sortilin interaction by binding to epitopes in and not far from NTIS2 (Biilmann-Rønn et al., 2017; Rosenthal, Schwabe, & Kurnellas, 2016).

We observed that the conformational response of PGRN to Sortilin binding was restricted to the C-terminal of PGRN. In direct support of this finding, PGRN binding to Sortilin has been shown to be abolished by deletion of the three last residues in the C-terminal (Hu et al., 2010; Zheng et al., 2011). However, the large conformational impact of Sortilin to PGRN binding, involving both a large surface of NTIS1 as well as NTIS2, could suggest additional direct interaction points in PGRN than just the very tip of the C-terminal. Especially, when the limited conformational response of Sortilin to AF38469 binding, which is restricted to NTIS1, is considered (**Supplementary Fig. 3**). The missing interaction sites could be present in PGRN in the areas, where HDX data were not obtained for the PGRN-Sortilin complex (**Fig. 5** and **Supplementary Fig. 5**).

In general, when attempting to derive a structural model of a protein complex based on HDX data alone two challenges arises: 1) Conformational effects in the direct binding interface are not readily distinguishable from indirect conformational effects observed spatially remote from the binding interface (allosteric effects) (Wales & Engen, 2006), and 2) no spatial or orientational restraints are obtained. However, taken together with evidence from prior biochemical studies, inferences concerning the structure of the Sortilin-PGRN complex can be made. Our HDX results suggest that the C-terminal of PGRN is interacting with a similar binding pocket in NTIS1 as employed by the C-terminal of NT, as also suggested in the literature (Zheng et al., 2011). Whether the C-terminal is inserted into the binding pocket from the top or the bottom of the β-propeller is not obvious from our HDX data. However, HDX reduction in NTIS1 is most pronounced for residues on the top of the β-propeller (residues 271-272 and 307-317) and the rest of the areas affected by the presence of PGRN are likewise found on the top of the β-propeller (**Fig. 5** and **Supplementary HDX-plots**). Furthermore, human anti-Sortilin antibodies able to inhibit PGRN binding bind to a region encompassing NTIS2 and its vicinity on the top of the β-propeller (Biilmann-Rønn et al., 2017; Rosenthal et al., 2016). Finally, the access to the cavity of the β-propeller from below is partly blocked by the N-linked glycosylation found at Asn373 (Quistgaard et al., 2009). Together, these observations thus suggest an interaction mode for PGRN and Sortilin, where the C-terminal of PGRN is inserted into the C-terminal NT binding pocket in NTIS1 from the top side of the β-propeller.

The model discussed above further implies that the reduced HDX observed in the 10CC-b domain upon PGRN binding is of an indirect/allosteric character, as was also indicated by HDX data on proNGF binding. It has in one earlier study been tentatively suggested that ligand binding in NTIS1 could be transmitted to the 10CC domains via a salt bridge interaction between Arg293 and Glu667 (Quistgaard et al., 2009). For both proNGF and PGRN binding, the reduced HDX in the 10CC-b domain strongly indicate allosteric signalling in Sortilin from NTIS1, or the nearby protected region of the outer rim of the propeller, to the 10CC-b domain (**Figs. 4** and **5**). Interestingly, upon PGRN binding, reduced HDX was observed in a segment of Sortilin (residues 641-644) that could bridge the reduced HDX observed in NTIS1 with the conformational stabilization observed in the 10CC-b domain (**Fig. 5**). Furthermore, a slight, but less pronounced reduction in HDX was also seen for the same residues upon proNGF and GSTpro binding. However, perhaps due the lower binding affinity of proNGF and GSTpro (Feng et al., 2010; Nykjaer et al., 2004), this HDX reduction did not exceed the general threshold value used in this study to conclude significance (**Supplementary HDX plots**). By inspecting the crystal structure, the residues Gln308 and Phe644, both placed in regions of Sortilin exhibiting reduced HDX upon PGRN binding, are in close proximity for propagating an allosteric signal from NTIS1 to the 10CC-b domain. However, any specific involvement of these two residues in transmitting a conformational change from the primary ligand binding region (NTIS1) on the top side of the propeller to the 10CC-b domain needs to be confirmed by future studies.

We note that the changes in HDX observed in the 10CC domain upon binding the larger protein ligands proNGF and PGRN could, in theory, also be due to an additional direct interaction site between protein ligands and the 10CC domain. However, we deem such a “tertiary” interaction site unlikely for the following reasons: 1) we did not observe separate regions in proNGF and PGRN impacted by Sortilin binding and 2) we observed that the pro-peptide of NGF was sufficient to induce the full binding impact on Sortilin, including effects in the 10CC domain. Furthermore, the NGF pro-peptide would need to span and interact with both NTIS1 in the beta-barrel as well as the 10CC domain which would involve at a minimum 23 residues (corresponding to approximately 85 Å solvent accessible surface distance (Matthew Allen Bullock, Schwab, Thalassinos, & Topf, 2016)) out of the total length of the pro-part of 103 residues. In doing so, a significant proportion of proNGF would thus need to adopt regions of ordered and less dynamic backbone structure. Our HDX data does not support such a notion, as the HDX of the backbone of the pro-part of NGF is unaffected by Sortilin binding and remains highly flexible even in the bound state. Irrespective of the direct or indirect nature of the observed stabilization in the membrane-proximal 10CC domain, it remains a unique conformational consequence of proNGF and PGRN binding, and as such could be important in ligand-specific downstream signaling.

The 10CC domains have shown to be highly flexible, as their structure dramatic changes upon change in pH and the oligomeric state of Sortilin (Januliene et al., 2017; Leloup et al., 2017). Furthermore, in full-length Sortilin the 10CC domains are attached to the transmembrane helix, which connects the extracellular part of Sortilin with its cytoplasmic tail. Our observation of structural stabilization in the 10CC-b domain upon ligand binding of proNGF and PGRN could thus be of biological relevance, as a part of a conformational trigger mechanism. A trigger mechanism that either communicate a binding event on the extracellular side of the receptor across the membrane directing the Sortilin-ligand complex for endocytosis and lysosomal degradation (PGRN) or for recruitment of associated pro-apoptosis adapter proteins, such as P75NTR (proNGF). Additional biological studies are needed to confirm this hypothesis.

In summary, in this work we used HDX-MS to investigate the conformational dynamics of Sortilin in complex with several biological ligands. Our results revealed that all investigated ligands employ two separate binding surfaces opposite each other in the cavity of the β-propeller. A significant difference was observed in the conformational response of Sortilin to binding of peptide ligands (Sortilin propeptide and NT) compared to protein ligands (proNGF and PGRN). Most notably, the protein ligands caused conformational stabilization of residues in the 10CC-b domain, hinting at a previously unknown conformational link between the ligand binding region and this membrane-proximal region of the receptor. We suggest that this observed indirect stabilization could be implicated in the transmission of a ligand-binding signal across the membrane to initiate intracellular signalling. Overall, the obtained new insights into the solution-phase conformational dynamics of unbound and ligand-bound Sortilin, unravel how the conformation of the Sortilin receptor is impacted by its main endogenous peptide and protein ligands. This information could pave the way for a better understanding of how Sortilin exert endocytosis, sorting and apoptotic signalling in a ligand-dependent manner.

## MATERIALS

All reagents were purchased from Sigma-Aldrich, St. Louis MO, USA in analytical grade except the following: immobilized pepsin beads (Thermo Scientific, Waltham, MA, USA), recombinant Nerve growth factor-β (residues 1-117) (Sino Biological Inc., Beijing, China), glutathione-S-transferase (GST) (GenScript, Piscataway, NJ, USA), proSort (GenScript), GlycoWorks RapiFluor-MS N-Glycan Kit (Waters Inc., Milford, MA, USA), tris(2-carboxyethyl)phosphine (TCEP) (Thermo Scientific) and AF38469 (H. Lundbeck A/S, Valby, Denmark).

### METHODS

### Expression and purification of proteins

#### Sortilin

The extra cellular soluble domain of human Sortilin (residue −33 to 723) was engineered with a C-terminal fXa and His tag extension (GSAMIEGRGVGHHHHHH). The construct was cloned into an episomal expression vector pCEP-PU and used for transient expression in HEK293 cell. Cell culture was harvested by centrifugation followed by sterile filtration and captured on a HisTrap Column (GE Healthcare, Little Chalfont, UK) equilibrated with 20mM Sodium phosphate pH 7.4, 1M NaCl. Sortilin was eluted in a linear gradient to 0.5M imidazole over 15 column volumes in the same buffer. Fractions were analysed by SDS-PAGE and pooled according to purity and concentration of Sortilin determined by bicinchoninic acid assay (BCA) (Pierce Biotechnology, Waltham MA, USA). The pooled fractions was further purified by size-exclusion chromatography on a Superdex200 column (GE Healthcare) in a phosphate-buffered saline (PBS) solution of 2.67mM KCl, 1.47mM KH_2_PO_4_, 137.93mM NaCl, 8.06mM Na_2_HPO_4_ · 7H_2_O, pH 7.4 (Gibco, Thermo Fischer Scientific). Fractions were pooled according to purity (SDS-PAGE) and the final protein concentration was determined by BCA. Samples used for mapping the conformational response of PGRN to Sortilin binding were buffer exchanged by spin filtering (Vivapsin 500, Sartorius, Göttingen, Germany) into 20mM tris(hydroxymethyl)aminomethane (Tris), pH 7.4 and the protein concentration was determined by UV-wavelength spectrometry (NanoDrop 2000, Thermo Scientific).

#### Progranulin

A synthetic gene of human Progranulin gene was cloned into pcDNA 3.1 for expression in the freestyle system (Thermo Fischer Scientific). The culture media was clarified by centrifugation and sterile filtration. Subsequent purification from culture media was done by capture on Capto-Q (GE Healthcare) in 20mM Tris pH 8.0, 1M NaCl followed by elution in a gradient to 0.5M imidiazole in same buffer over 10 column volumes. Fractions eluting from the column was precipitated with 2M (NH_4_)_2_SO_4_ and centrifuged. The precipitate was dissolved in PBS and separated by size-exclusion chromatography on a Superdex200 column (GE Healthcare). Fractions with Progranulin were identified by SDS-PAGE and pooled accordingly. Samples used for mapping the conformational response of PGRN to Sortilin binding were buffer exchanged by spin filtering (Vivapsin 500, Sartorius) into 20mM Tris, pH 7.4 and the protein concentration was determined by UV spectrophotometry (NanoDrop 2000, Thermo Scientific).

#### The pro-form of Nerve growth factor-β

Human proNGF is difficult to express and purify in high homogenous yield, because of the presence of three Furin cleavage sites in the pro-part. To meet this challenge a triple mutant of these Furin cleavage sites has been developed (Feng et al., 2010; Pagadala, Dvorak, & Neet, 2006). In the current study human glycosylated proNGF was expressed in a mammalian host cell as described in the literature (Esben Trabjerg et al., 2017).

#### GSTpro

GSTpro was engineered as a fusion of glutathion S-transferase (GST) merged at the C-terminal of GST to the pro part of proNGF. The construct was cloned into pGEX expression plasmid and used for expression in *E. coli* using the Overnight Express™ Autoinduction System 1 (Novagen Inc., Merck Corporation, Darmstadt, Germany). The cells were harvested, lysed and GSTpro was purified from the supernatant using standard Glutathion-sepharose affinity chromatography.

### Quality control of proteins

The intact mass of all protein constructs and peptides was determined by either electrospray ionization (ESI) mass spectrometry (MS) or matrix-assisted laser desorption/ionization MS (data not shown). Furthermore, the sequence of all proteins was confirmed by generating a tryptic peptide map of all investigated proteins (data not shown). In short, the proteins were denatured (6M guanidinium hydrochloride, 30min at 60°), reduced (10mM DTT, 2h at 60°C) alkylated in the dark (20mM iodoacetamide, 30 min at 25°C) and digested with Trypsin (w/w 1:20 of enzyme:protein, 16h at 37°C). The tryptic peptide mixture was acidified with formic acid and loaded onto a reverse-phase UPLC-system (nanoAcquity, Waters Inc.), trapped on a C18-trap column (ACQUITY UPLC BEH C18 1.7µM VanGuard column, Waters Inc.) and desalted for 6min at 200µL/min. The trap column was subsequently put in-line with a C18-analytical column (ACQUITY UPLC BEH C18 1.7µm, 1 × 100mm column, Waters Inc.), where the tryptic peptides were separated by reverse-phase chromatography and ionized by positive ESI into a Q-TOF mass spectrometer (Synapt G2 or Synapt G2Si, Waters Inc., Wilmslow, UK). The tryptic peptides were analysed by tandem mass spectrometry (MS/MS) with collisional induced dissociation (CID). The mass spectrometer was operated in data-dependent acquisition mode (DDA). The acquired mass spectra were lock mass corrected against Glu-1-fibrinopeptide B (GFP) and analyzed in the PLGS 3.0 software, which matches precursor- and fragment ions to a local protein database. All positive hits were manually validated. Oxonium fragment ions of N-acetylglucosamine, sialic acid subtracted water and N-acetylglycosamine connected with galactose (204.0875m/z, 274.1732m/z and 366.14m/z) were used as reporter ions to manually identify N-linked glycopeptides (Cao et al., 2014).

### Glycan analysis of Sortilin

The glycan analysis was performed as described for the GlycoWorks RapiFluor-MS N-Glycan Kit (Waters Inc.). In short Sortilin was denatured, reduced and treated with PNGase F. The released glycans were labelled with an easily ionized fluorophore and extracted by solid-phase-extraction. Following extraction the labelled glycans were separated by hydrophilic interaction liquid chromatography (HILIC) (nanoAcquity, Waters Inc.) and detected by fluorescence (Acquity UPLC FLR Detector, Waters Inc.) and ESI-MS (Xevo G2-XS QTof, Waters Inc.).

### HDX-MS experiments

#### Formation of protein complexes

The protein complexes were formed by incubating the binding partners for at least 15min at 25°C before initiation of the HDX experiment to secure the labelling process to take place at steady-state conditions. The amount of ligand was adjusted to secure that at least 90% of the investigated protein/peptide was bound to its ligand, except for Sortilin bound to NT and GST (**Supplementary table 2**). NT was added in 1.5 and 15 times molecular excess to be able to compare the results with the published crystal structures (Quistgaard et al., 2014, 2009) and GST was added in the same molecular amount as GSTpro.

#### Exchange reaction

The exchange reaction was started by diluting the target protein or peptide (10-20pmol in 150mM NaCl, 20mM Tris pH 7.4) in absence or presence of ligand 1:9 with 99% D_2_O (150mM NaCl, 20mM Tris pD_read_ 7.6) at 25°C. After various time intervals (**Supplementary table 3**) the exchange reaction was quenched by addition of ice cold quench buffer (0.8M TCEP, 2M glycine buffer pH 2.3) in an 1:1 relationship, thereby decreasing the pH to 2.3. The quenched samples were immediately frozen to −80°C until analysis. Equilibrium deuterated control samples were prepared by incubating samples for 24h at denaturing conditions (6M deuterated GdnHCl) 25°C. At least two time points for all states were recorded in triplicate except for AF38469 binding to Sortilin and Sortilin binding to proSort, proNGF and GSTpro (**Supplementary table 3**). For investigation of the impact of Sortilin binding to proSort, proNGF and GSTpro the samples was quenched with ice-cold 300mM phosphate buffer pH 2.3. For investigation of the impact of Sortilin binding to PGRN the samples were prepared in buffers depleted for NaCl and quenched with ice-cold 2% formic acid (FA). Furthermore, the equilibrium labelled PGRN sample was prepared by labelling a PGRN sample for 6 days.

#### UPLC-HDX-MS setup

The quenched samples were thawed and injected onto a cooled (0°C) reverse-phase UPLC-HDX-system (Waters Inc.) with an online home-packed pepsin column (internal volume of 60µL). The resulting peptic peptides were trapped unto a C18-trap column (ACQUITY UPLC BEH C18 1.7µM VanGuard column, Waters Inc.) and desalted for 3min at 200µL/min with 0.23% FA in water. The trap column was put in-line with a C18 analytical column (ACQUITY UPLC BEH C18 1.7µm, 1 × 100mm column, Waters Inc.) and the peptic peptides were separated by conventional reverse-phase chromatography (solvent A: 0.23% FA in water, solvent B: 0.23% FA in acetonitrile) and ionized by positive ESI into a high resolution Q-TOF mass spectrometer (Synapt G2Si, Waters Inc.). Here, the peptides were separated according to their ion mobility and mass-to-charge. Identification of peptides was performed on fully reduced and non-deuterated samples by MS/MS using a combination of data independent acquisition (MSe) and DDA. For investigation of Sortilin binding to PGRN the described UPLC-HDX-MS setup was modified to implement online electrochemical reduction as described in the literature (Mysling et al., 2014; Esben Trabjerg et al., 2015).

### Data analysis

Peptide identification: The acquired mass spectra were lock mass corrected against GFP and analysed in PLGS 3.0, which matches precursor- and fragment ions to a local protein database. All fragment spectra of identified peptides were manually inspected. N-linked peptic glycopeptides were identified manually from data recorded by DDA acquisition. Determination of deuterium incorporation: The acquired mass spectra were lock mass corrected against GFP and the deuterium incorporation for all peptides was determined by the software DynamX 3.0 (Waters Inc.). Statistical analysis: All statistical analyses were performed in the Excel software (Microsoft, Redmond, WA, USA). All comparisons were performed with either a homoscedastic or a heteroscedastic Student’s T-test with an α-value set to 0.01. The choice to use either a homoscedastic or a heteroscedastic Student’s T-test was determined by using an F-test with the α-level set to 0.05. The F-test compares the variance of deuterium uptake of a single peptide at a single time point from two different states. In the comparative HDX-MS analysis of protein or peptides in absence or presence of a ligand, a protein segment was considered to experience a significant difference in HDX if all of the following three requirements was fulfilled.1) A triplicate data point shows a significant difference in deuterium incorporation (p<0.01), 2) the absolute difference in deuterium incorporation was four times larger than the propagated standard error for the difference 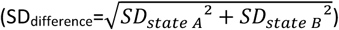 and 3) overlapping peptides showed similar HDX kinetics to dismiss false positive results.

## ACKNOWLEDGMENTS

We would like thank H. Lundbeck for providing protein material and AF38469. We gratefully acknowledge financial support from The Innovation Fund Denmark (Grant No. 1355-00165), The Marie Curie Actions Programme of the E.U (Grant No. PCIG09-GA-2011-294214), and the Danish Council for Independent Research|Natural Sciences (Steno Grant No. 11-104058).

## AUTHOR CONTRIBUTIONS

S.C. and K.D.R. conceived and coordinated the study. S.C. and F.K. designed and constructed the vector for expression of Sortilin, PGRN and proNGF and performed the expression and purification. N.A.A. investigated the conformational impact of Sortilin binding to PGRN Z.W. investigated the conformational impact of AF3864 to Sortilin. E.T. conducted all other mass spectrometry experiments. S.C., K.D.R. and E.T. analysed the data and wrote the paper. All authors approved the final version of the manuscript.

---

**Table S1.**
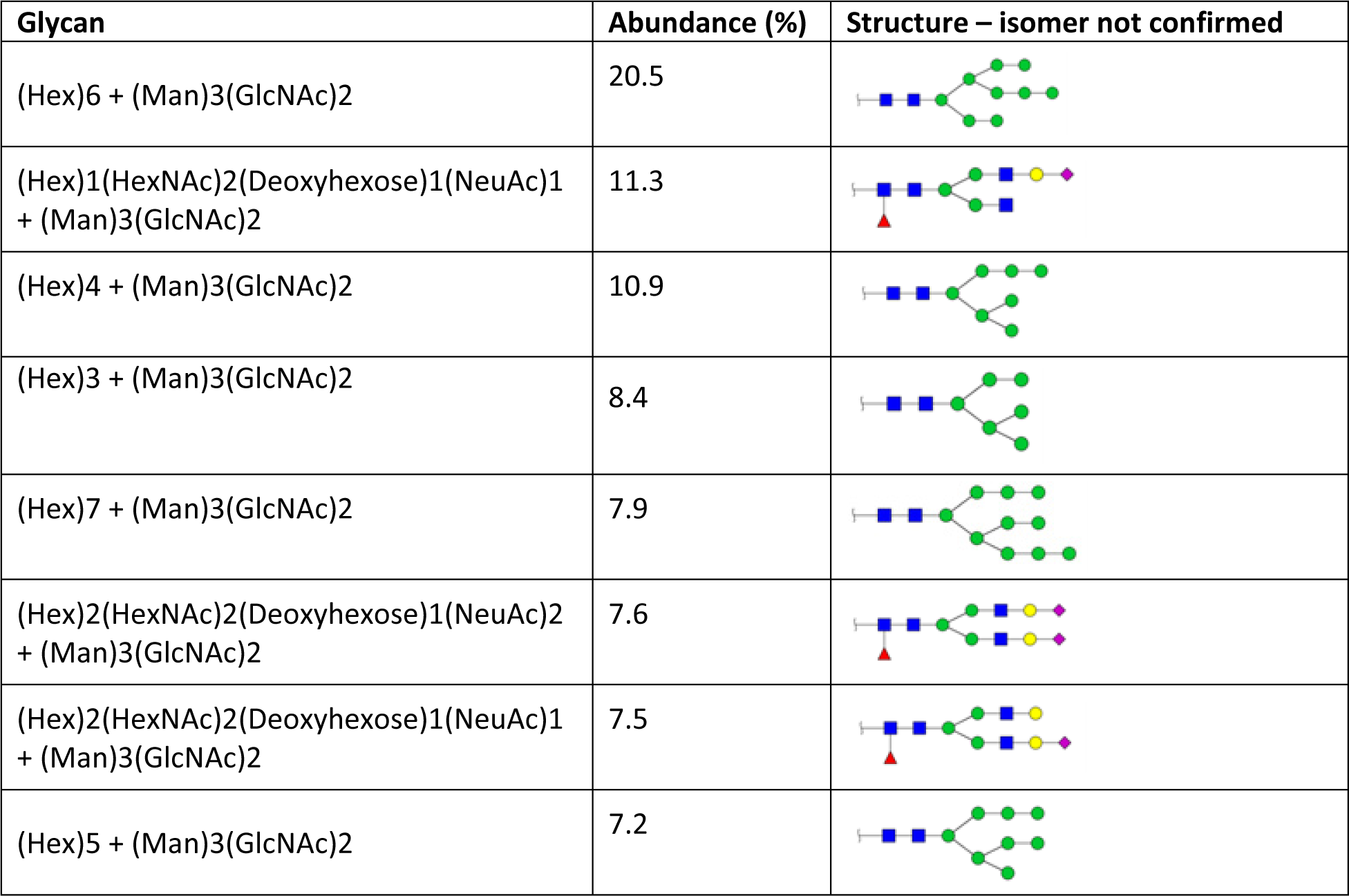
Analysis of PNGase F released glycans from Sortilin. Sortilin was denatured, reduced and treated with PNGase F as described by the vendor of the GlycoWorks RapiFluor-MS N-Glycan Kit. The released glycans were labelled with an easily ionized fluorophore and extracted by solid-phase-extration. The labeled glycans were separated by hydrophilic interaction liquid chromatography (HILIC) and detected by fluorescence and mass spectrometry. The table shows all identified N-linked glycans with abundance higher than 5%. The exact glycan isomer has not been confirmed by mass spectrometry. Blue square: N-acetylglucosamin, yellow circle: galactose, green circle: mannose, purple diamond: sialic acid, red triangle: fucose.

| Glycan | Abundance (%) | Structure – isomer not confirmed |
| --- | --- | --- |
| (Hex)6 + (Man)3(GlcNAc)2 | 20.5 |  |
| (Hex)1(HexNAc)2(Deoxyhexose)1(NeuAc)1 + (Man)3(GlcNAc)2 | 11.3 |  |
| (Hex)4 + (Man)3(GlcNAc)2 | 10.9 |  |
| (Hex)3 + (Man)3(GlcNAc)2 | 8.4 |  |
| (Hex)7 + (Man)3(GlcNAc)2 | 7.9 |  |
| (Hex)2(HexNAc)2(Deoxyhexose)1(NeuAc)2 + (Man)3(GlcNAc)2 | 7.6 |  |
| (Hex)2(HexNAc)2(Deoxyhexose)1(NeuAc)1 + (Man)3(GlcNAc)2 | 7.5 |  |
| (Hex)5 + (Man)3(GlcNAc)2 | 7.2 |  |

**Table S2.** Overview of affinity constants used in the present study. All investigated binding interfaces were investigated in presence of excess ligand to secure a saturation of at least 90% of the binding interface of the target protein/peptide.

| Sortilin ligand | Assay | K<sub>d</sub> (nM) | Reference |
| --- | --- | --- | --- |
| NT | Isothermal titrations calorimetry (ITC) | 85 | (Quistgaard et al., 2014) |
| AF38469 | Scintillation Proximity Assay (SPA) | Equipotent to NT | (Schrøder et al., 2014) |
| proSort | Surface Plasmon Resonance (SPR) | 20-30 | (C. Munck Petersen et al., 1999) |
| PGRN | Surface Plasmon Resonance (SPR) | 8 | (Hu et al., 2010) |
| proNGF | Surface Plasmon Resonance (SPR) | 5 | (Nykjaer et al., 2004) |
| GSTpro | Surface Plasmon Resonance (SPR) | 8 | (Nykjaer et al., 2004) |
| GST | Surface Plasmon Resonance (SPR) | No binding detected | (Nykjaer et al., 2004) |
| NGF | Surface Plasmon Resonance (SPR) | 87 | (Nykjaer et al., 2004) |

**Table S3.**
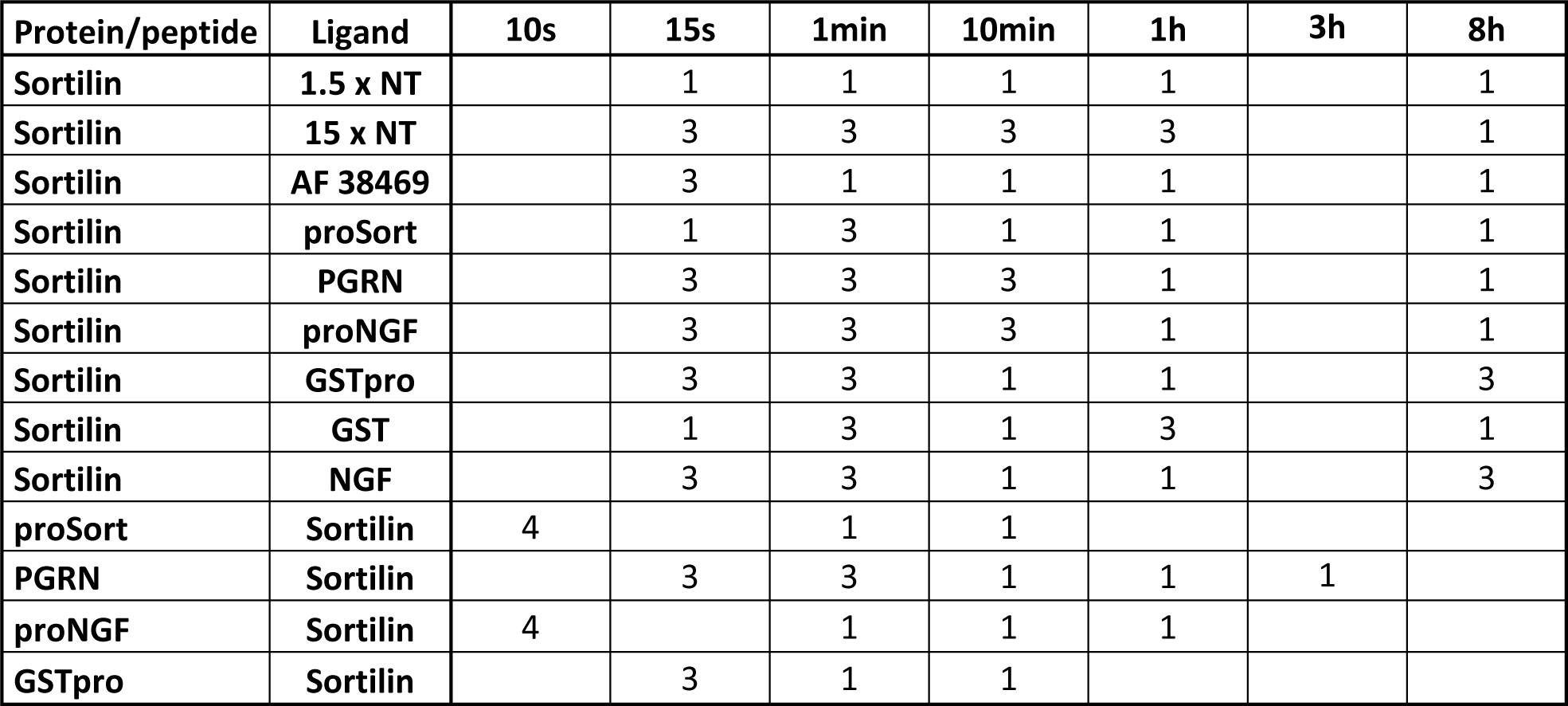
Overview of recorded HDX data. The numbers in the table refers to the number of replicate data for every single time point.

**Figure S1.**
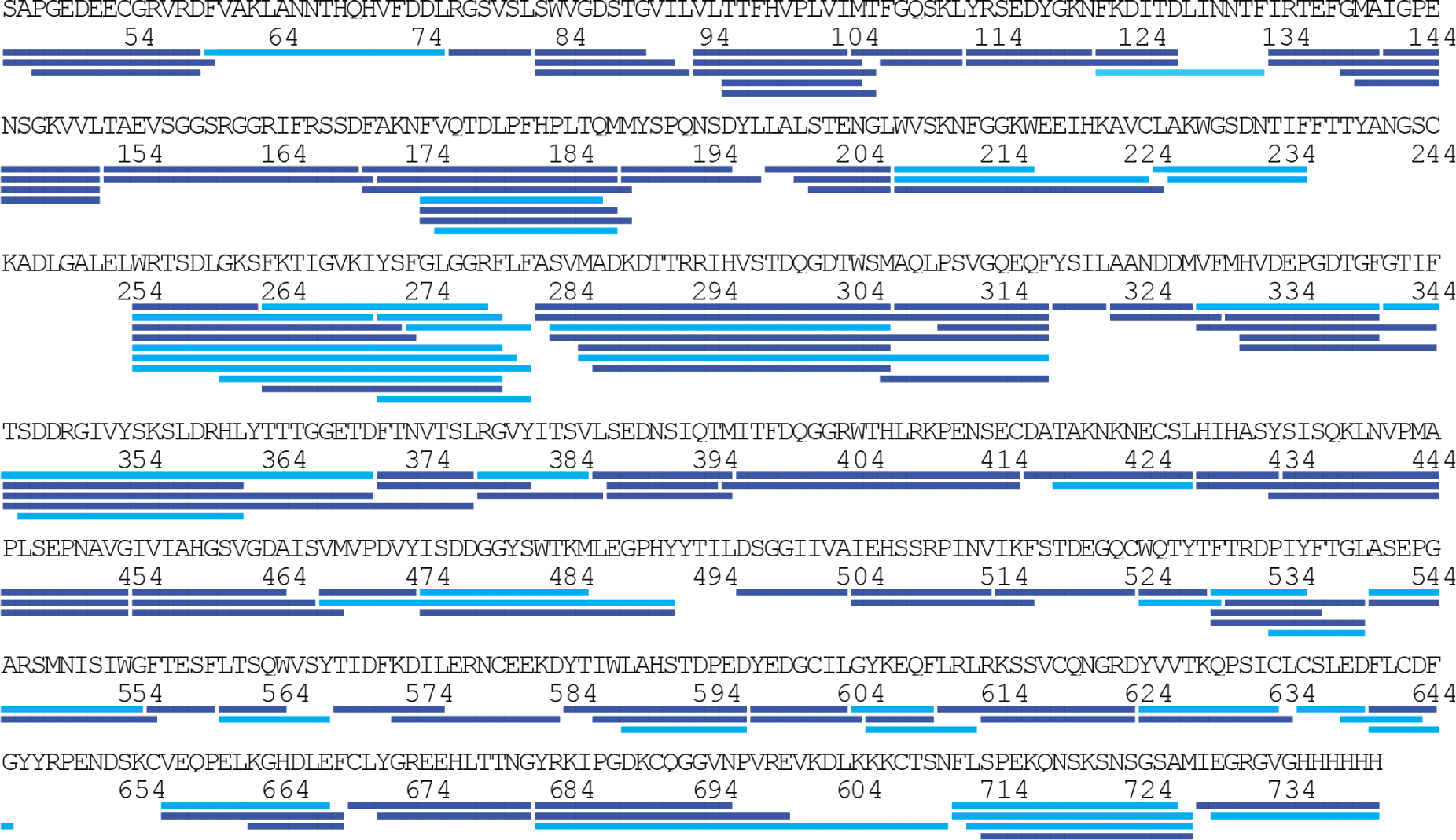
HDX-MS coverage map of Sortilin. The full sequence of the soluble extracellular domain for Sortilin is shown, including the C-terminal attached His6-tag. Peptic peptides from which HDX data could be obtained for all states are shown as blue bars. Light blue bar represents peptides, where HDX data was not obtained for all states.

**Figure S2.**
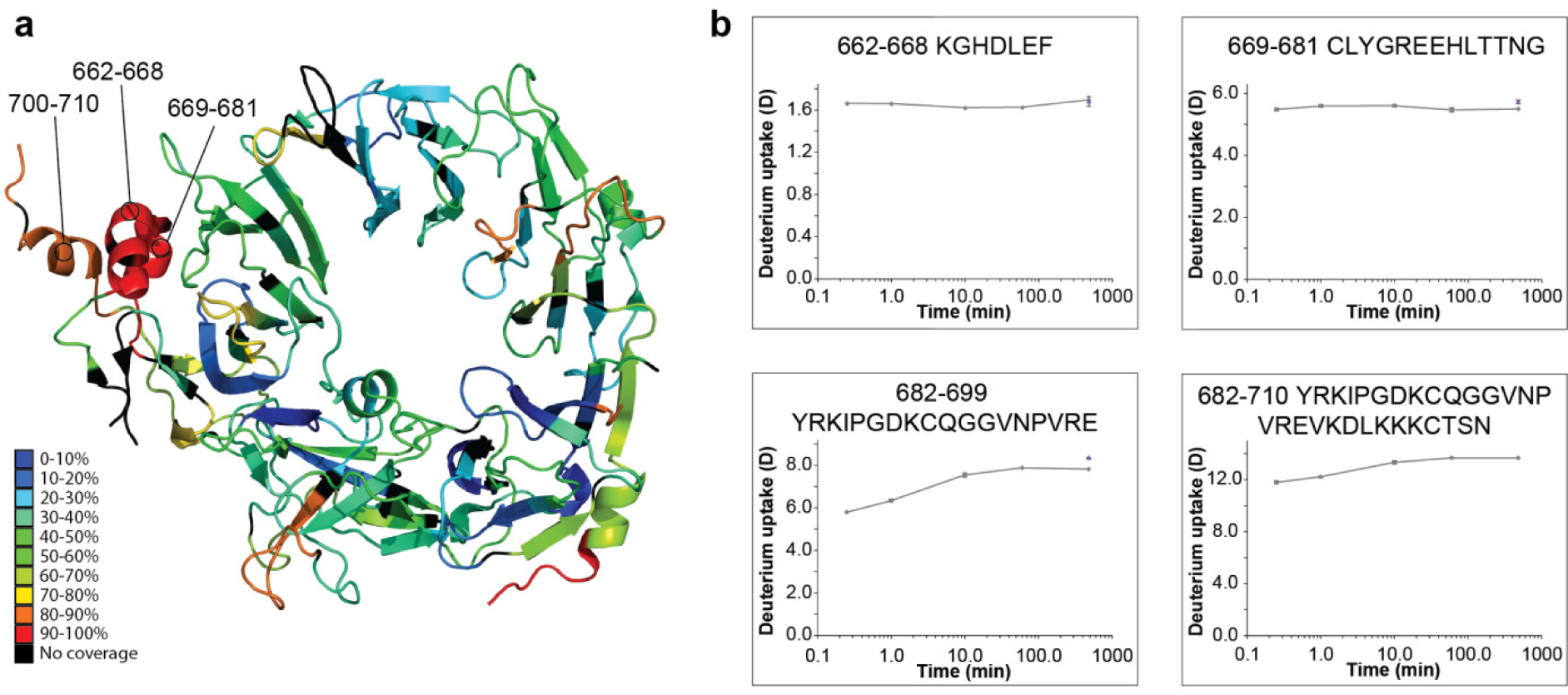
Observation of discrepancies between the X-ray crystal structure and the solution-phase conformational dynamics of Sortilin. The three N-terminal α-helices of the extracellular part of Sortilin show no or very little protection from exchange. (**a**) The crystal structure of Sortilin (PDB ID: 4PO7) are coloured according to the normalized deuterium incorporation after 15s. (**b**) Absolute deuterium incorporation is plotted as a function of time for Sortilin peptides spanning the three N-terminal helices. Equilibrium labelled (90%) Sortilin control samples are plotted as filled purple triangles at the 8h time point. SD is plotted as error bars (are only slightly visible). (n=3 for all time points and equilibrium labelled control sample, except the 8h time point where n=1)

**Figure S3.**
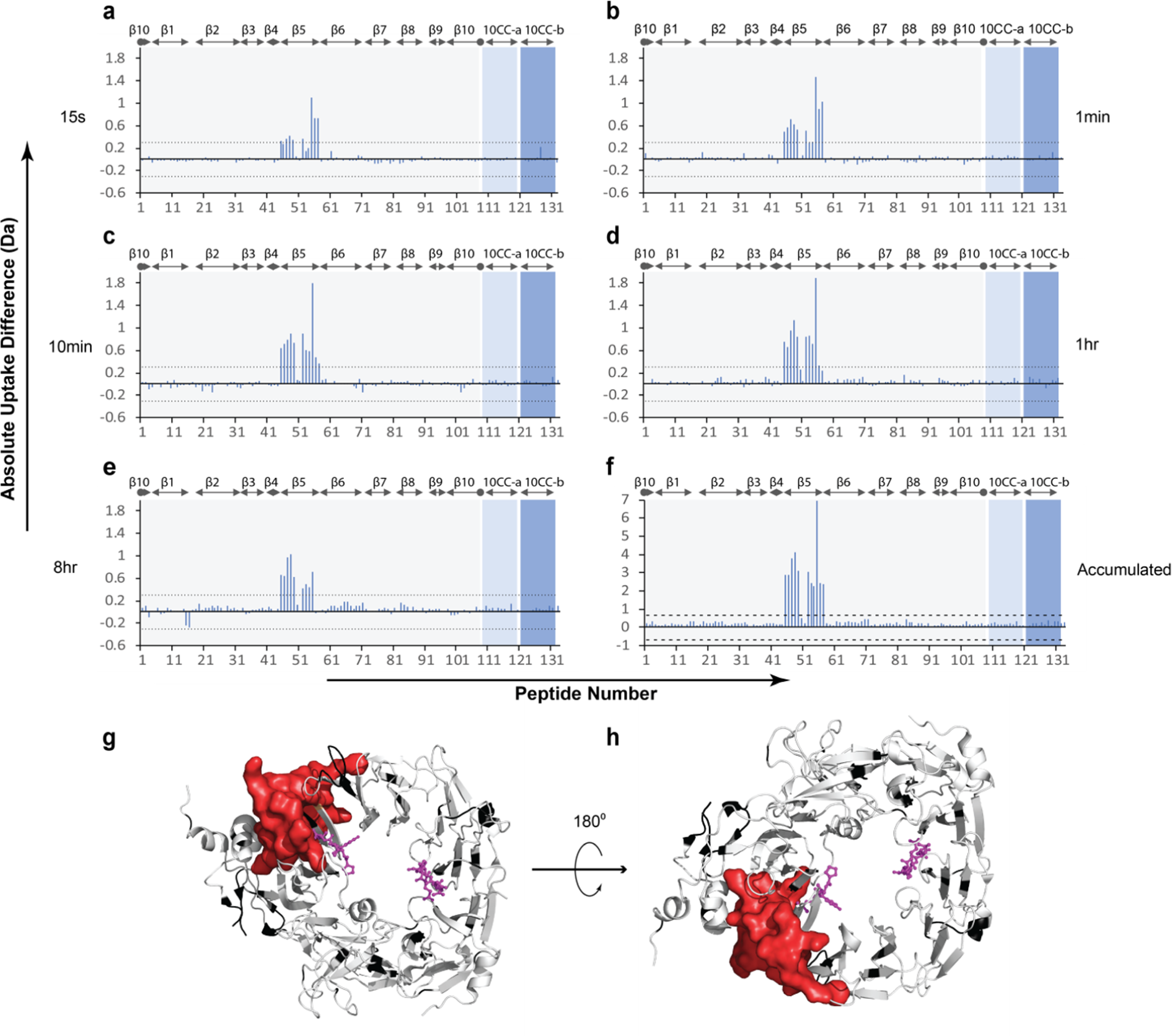
Conformational impact of AF38469 binding to Sortilin. **(a-e)** Absolute differences in deuterium uptake between Sortilin in the presence and absence of AF38469 at 5 exposure time points determined by HDX-MS. The horizontal axis denotes the peptide numbers from 1 to 133 starting from the N-terminal (**Supplemental HDX plots**). The vertical axis shows the absolute differences in deuterium uptake between the two states. Positive bars indicate a decrease in deuterium uptake upon binding of AF38469 to Sortilin, while negative bars indicate an increase in deuterium uptake upon binding of AF38469 to Sortilin. The dotted lines at ±0.3 Da indicate the 98% confidence limits for significant differences, as defined in the literature (Arora et al., 2015; Houde, Berkowitz, & Engen, 2011). The gray shades in the background represent blade 1 to 10 of the β-propeller, and the light blue and dark blue shades represent the 10CC-a and 10CC-b domains, the boundaries of which are labelled at the top of each chart. **(f)** Accumulated differences in deuterium uptake across all the exposure time points. The dashed lines at ±0.66 Da indicate the 98% confidence limits for significant differences, as defined in the literature (Arora et al., 2015; Houde et al., 2011). **(g)** Differential HDX results are mapped onto the crystal structure of Sortilin (PDB ID: 4PO7). Regions stabilized in the presence of NT are coloured red. Regions with no change in HDX are coloured white, while regions without coverage are coloured black. The C- and N-terminal of NT are shown in magenta. (n=1 for all time points, except the 15s time point, where n=3)

**Figure S4.**
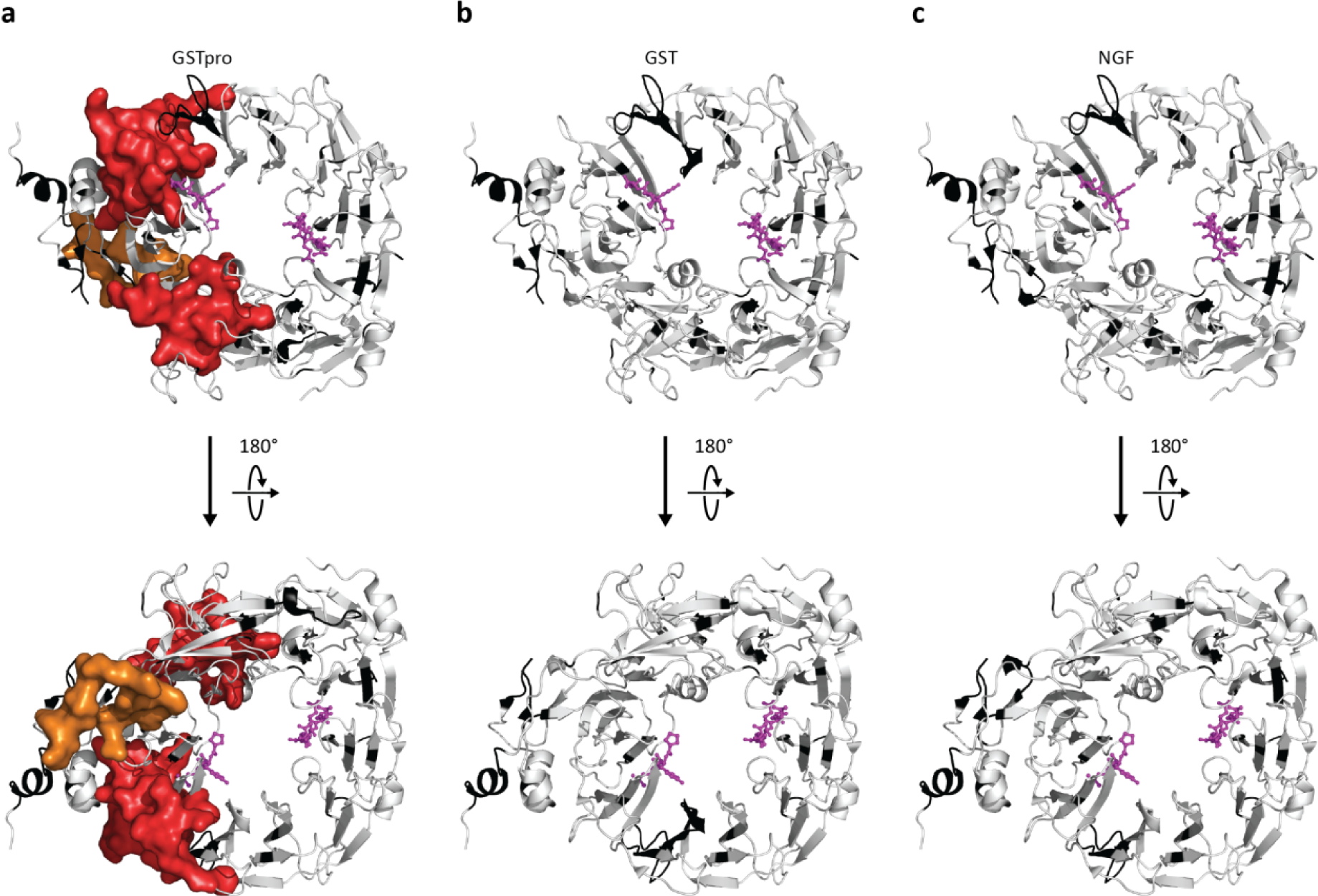
Conformational response of Sortilin in presence of GSTpro, GST or NGF. Differential HDX results of Sortilin in presence and absence of GSTpro (**a**), GST (**b**) and NGF (**c**) are mapped onto the crystal structure of Sortilin (PDB ID: 4PO7). Regions stabilized in the presence of proNGF are coloured red (β-propeller domain) and orange (10CC domains). Regions showing no change in HDX are coloured white, while regions without coverage are coloured black. The C- and N-terminal of NT are shown in magenta.

**Figure S5.**
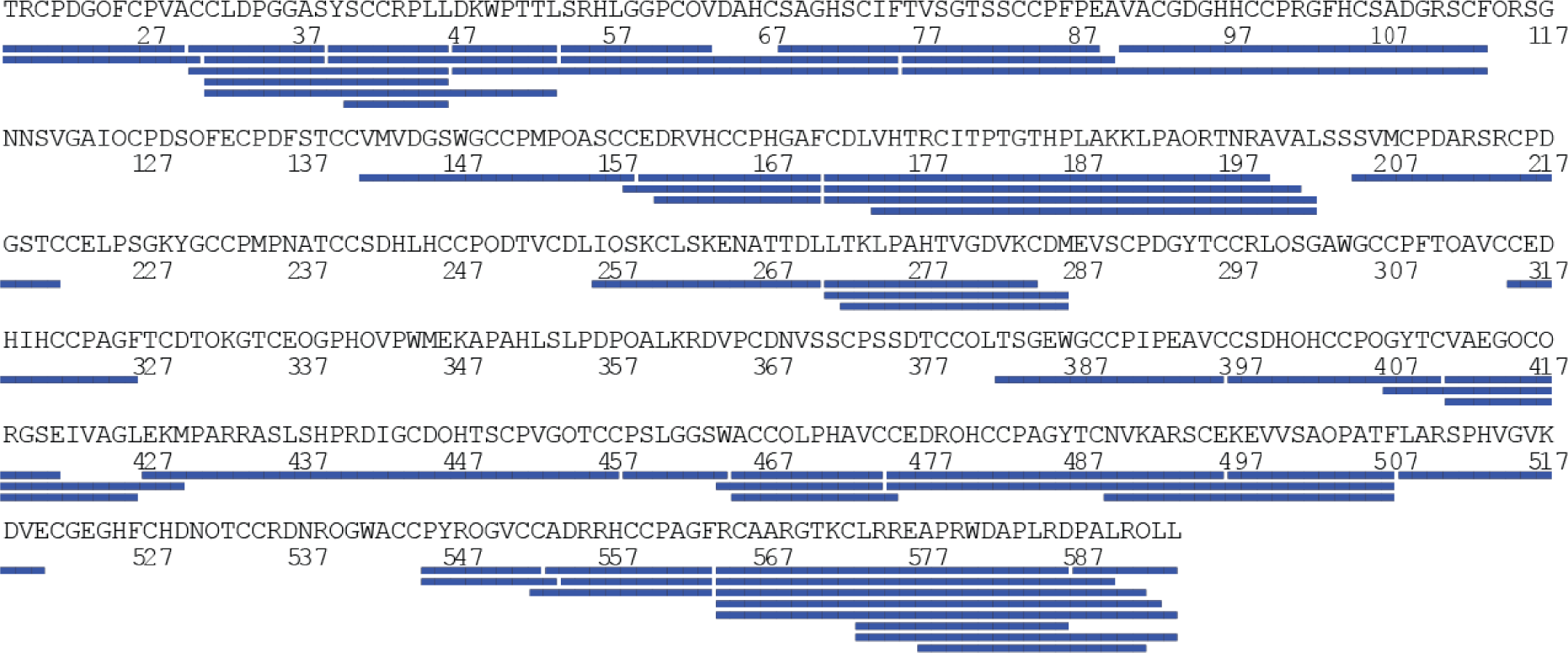
HDX-MS coverage map of PGRN. The full sequence of the PGRN construct used in the current study is shown. Peptic peptides from which HDX data could be obtained are shown as blue bars. PGRN contains 88 cysteines forming a total 44 disulfide bonds, which were very difficult to reduce by conventional chemical reduction by TCEP, resulting in a sequence coverage below 50% (data not shown). By implementing online electrochemical reduction into the HDX-MS workflow as described in the literature (Mysling et al., 2014; E. Trabjerg et al., 2015), the sequence coverage was markedly increased to 71%. The lacking sequence coverage can mostly be ascribed to the presence of five N-linked glycosylation sites at (Asn118, Asn236, Asn265, Asn368, Asn530), where coverage only was obtained for a single site (Asn265).

**Figure S6.**
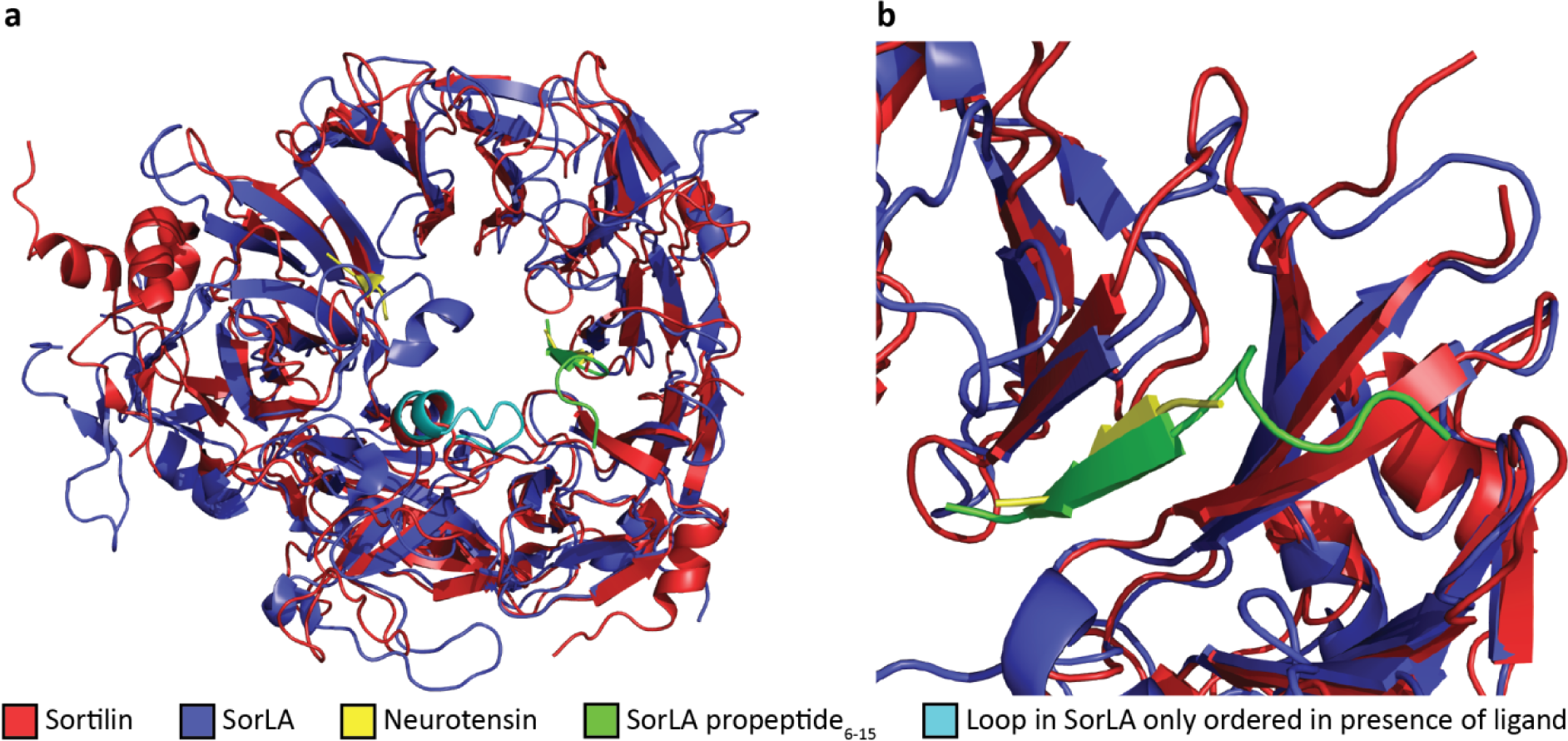
Comparison of ligand-bound Sortilin (PDB ID: 4PO7) and SorLA (PDB ID: 3WSY). **(a)** The peptide backbone of Sortilin (red) bound to NT (yellow) and SorLA (blue) bound to a fragment of its own propeptide (green) was aligned using the PYMOL software (DeLano Scientific LLC) (RMSD=3.2Å). The β-propellers for the two proteins align very well, while larger differences are observed for the 10CC domains. The cyan coloured loop in the SorLA structure is only ordered in the presence of ligands (fragments of SorLA propeptide or Aβ) **(b)** Zoom of the SorLA propeptide binding site and the secondary NT interaction site (NTIS2). Both ligands are employing a similar binding interface by forming a β-strand extension to blade 1 of both receptors.

## Supplementary HDX plots

HDX plots of Sortilin in presence of NT. Absolute deuterium incorporation is plotted as a function of time for Sortilin (gray lines), Sortilin in presence of 1.5 times molar excess of NT (red lines) and Sortilin in presence of 15 times molar excess of NT (blue lines). Equilibrium labelled (90%) Sortilin control samples are plotted as filled purple triangles at the 8h time point. SD is plotted as error bars (are only slightly visible). (n=3 for all time points and the equilibrium labelled sample, except the 8h time point where n=l for Sortilin and Sortilin in presence of 15 times molar excess of NT. n=l for Sortilin in presence of 1.5 times molar excess of NT)

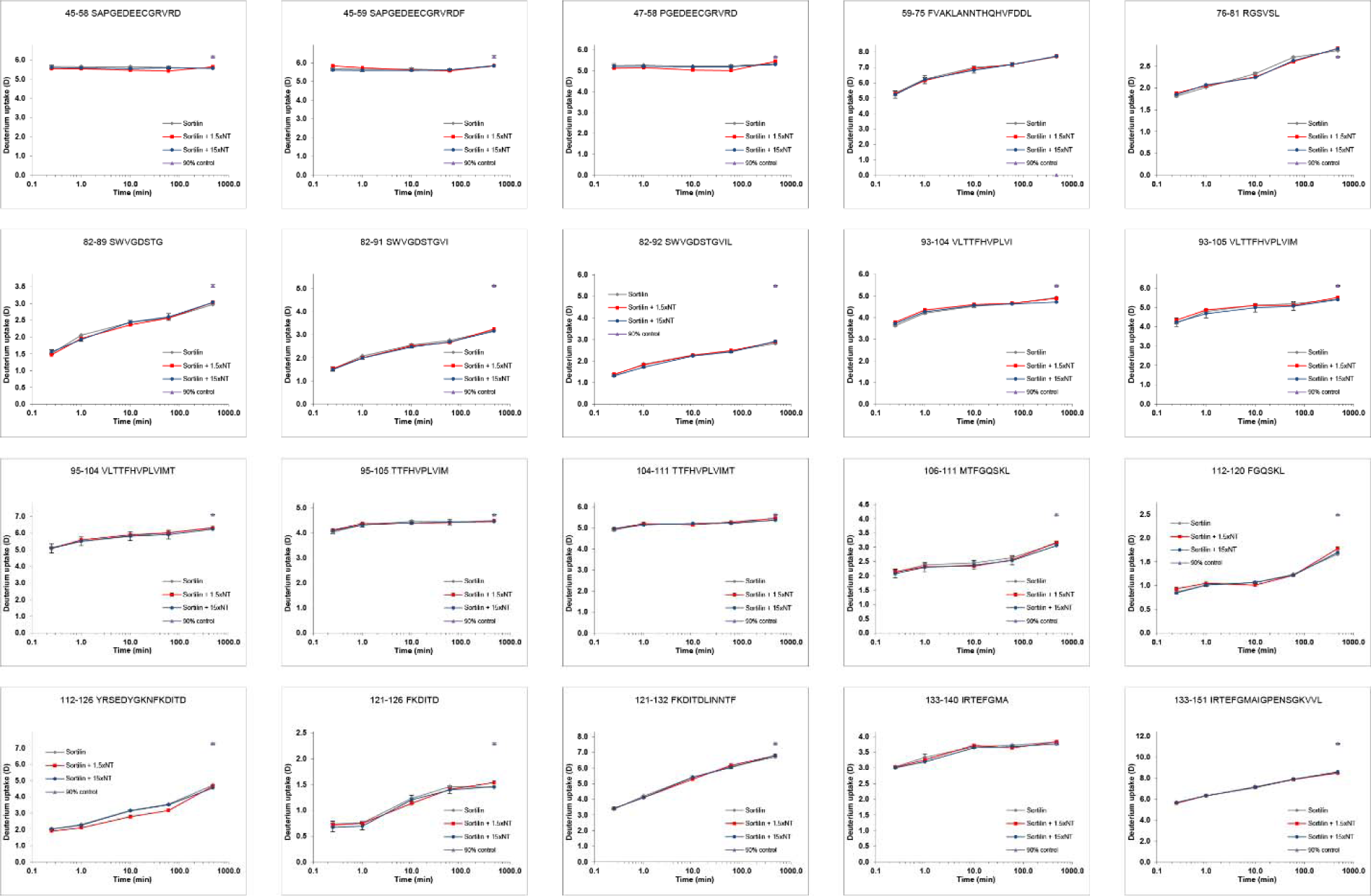

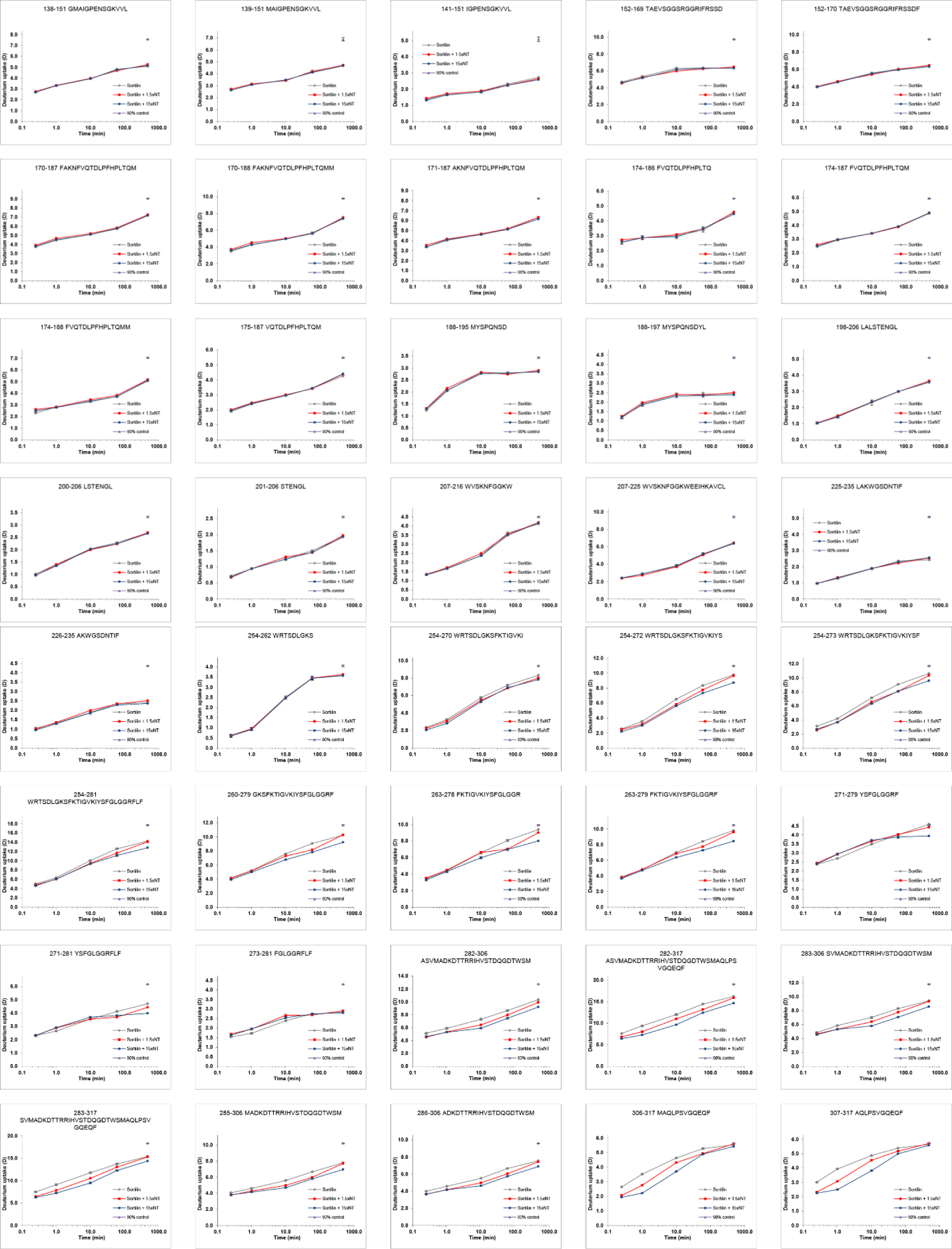

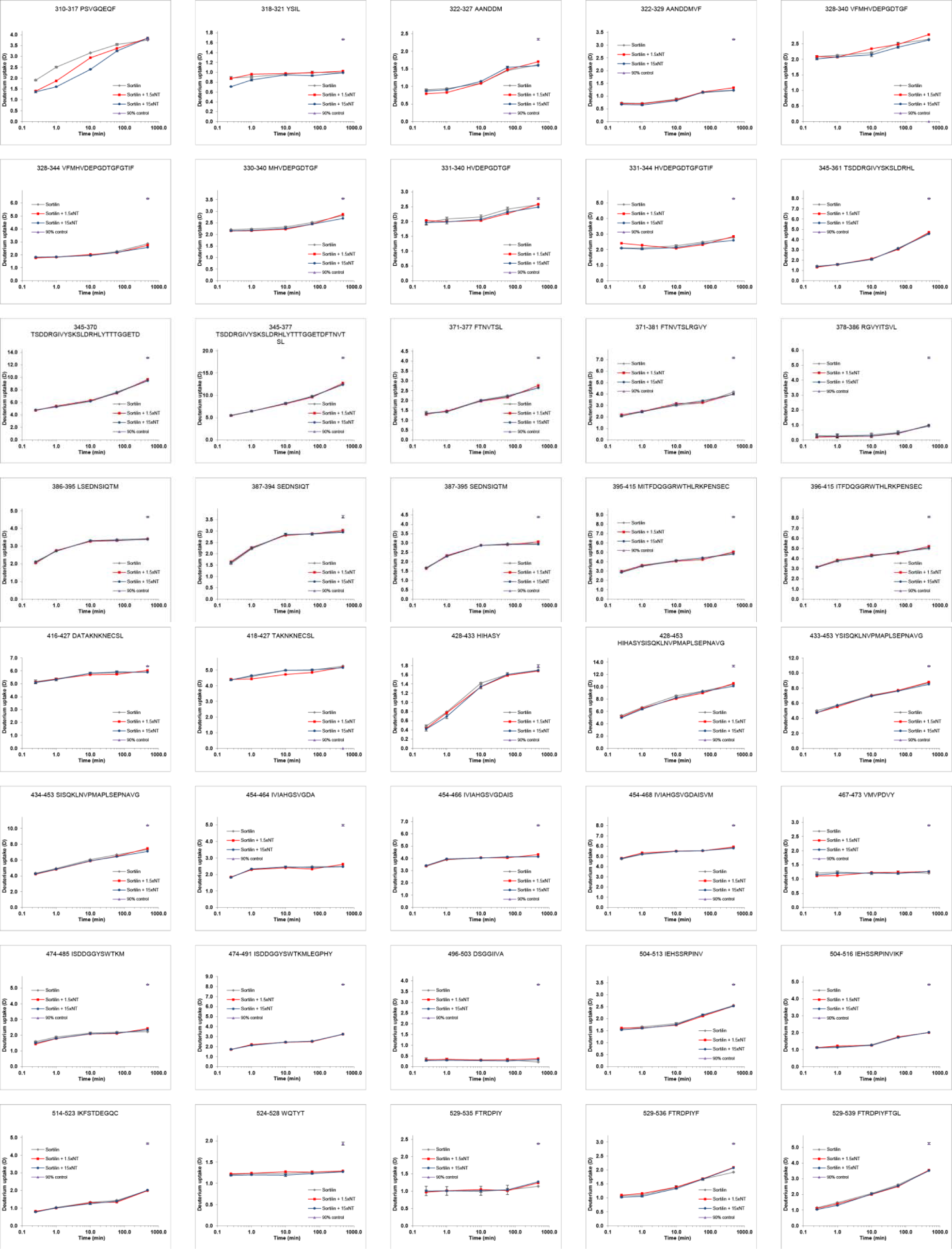

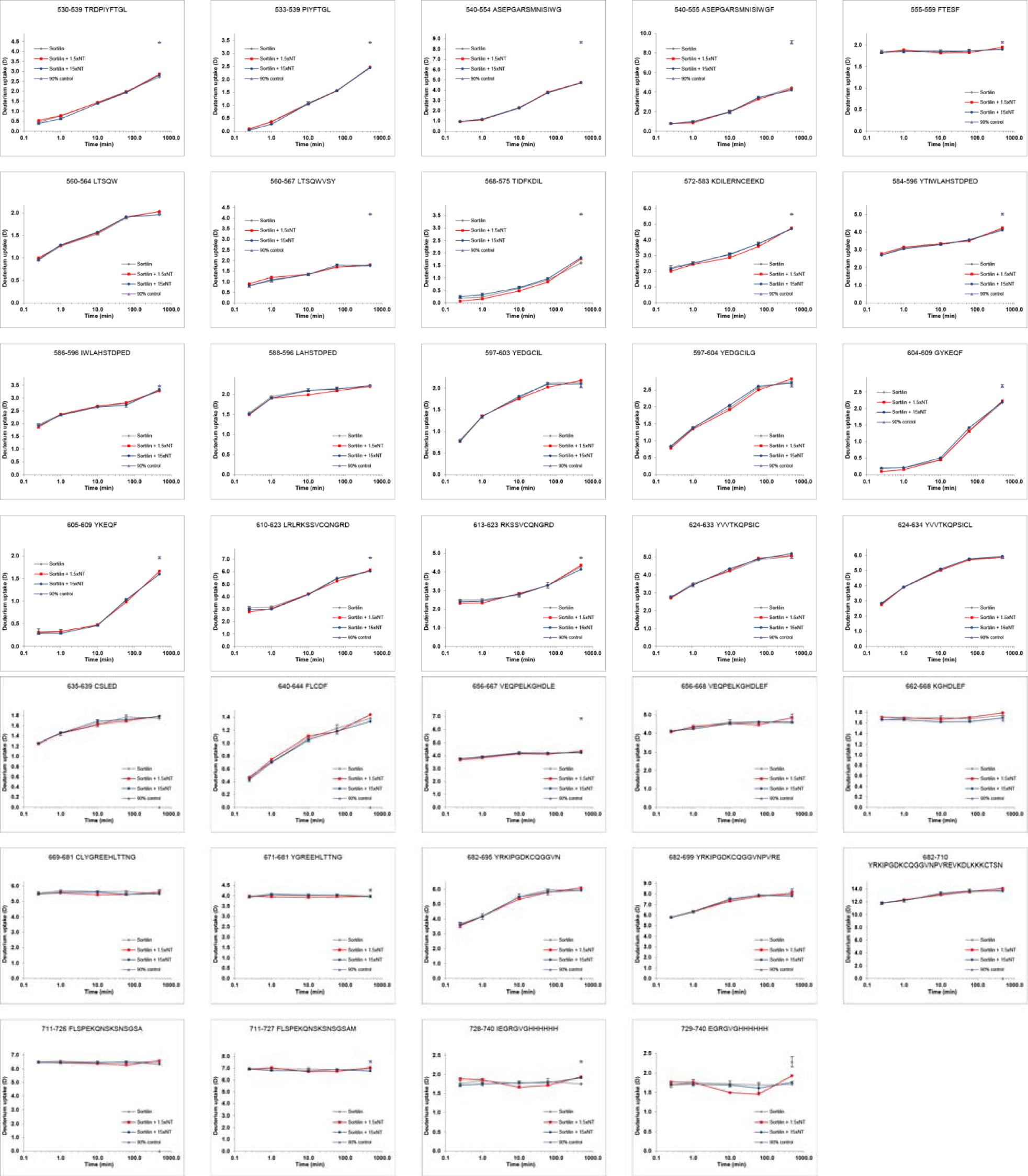

HDX plots of Sortilin in presence of AF38469. Absolute deuterium incorporation is plotted as a function of time for Sortilin (gray lines) and Sortilin in presence of AF38469 (red lines). Equilibrium labeled (90%) Sortilin control samples are plotted as filled purple triangles at the 8h time point. SD is plotted as error bars (are only slightly visible). (n=l for all time points, except the 15s time point, where n = 3)

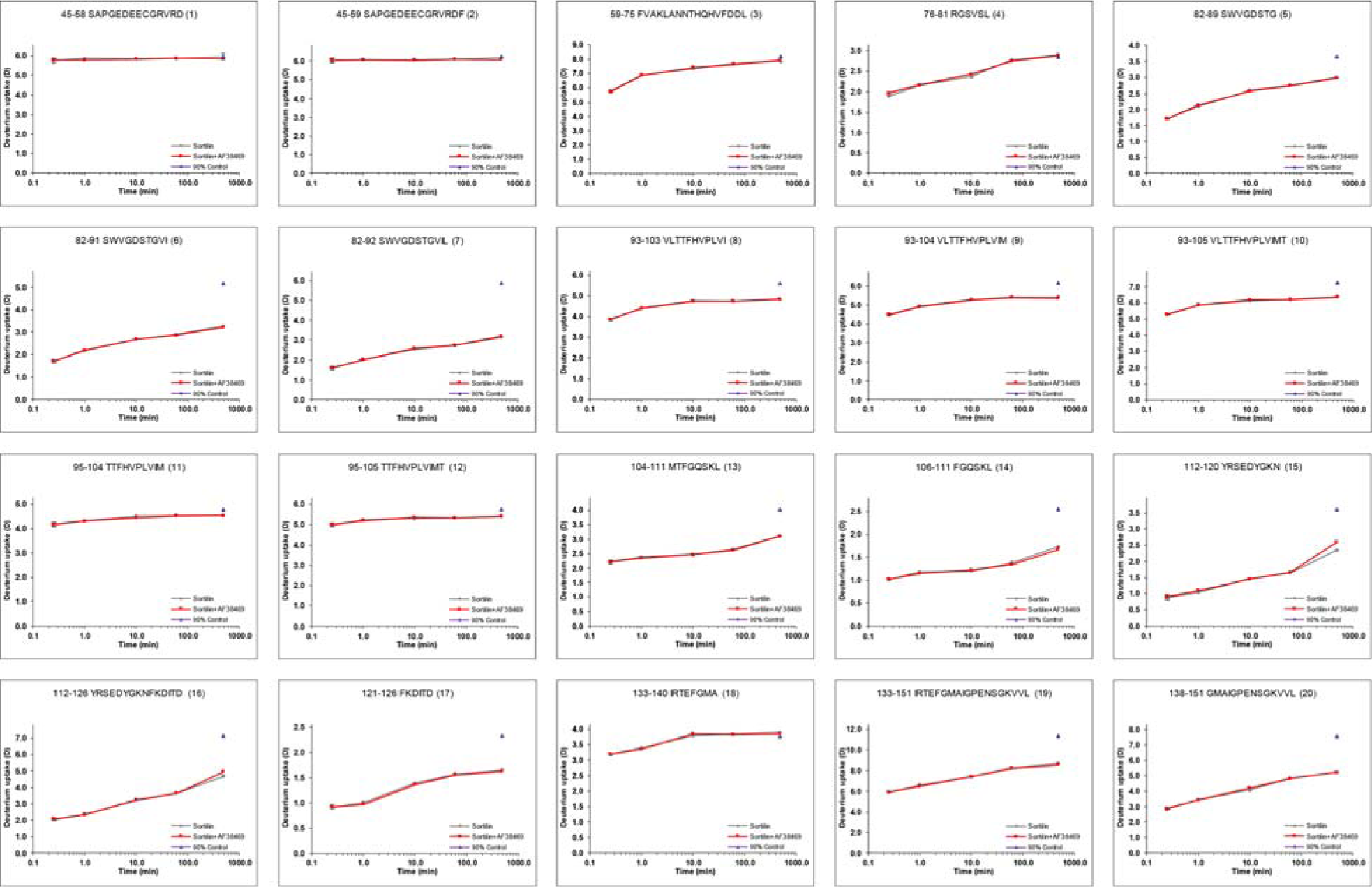

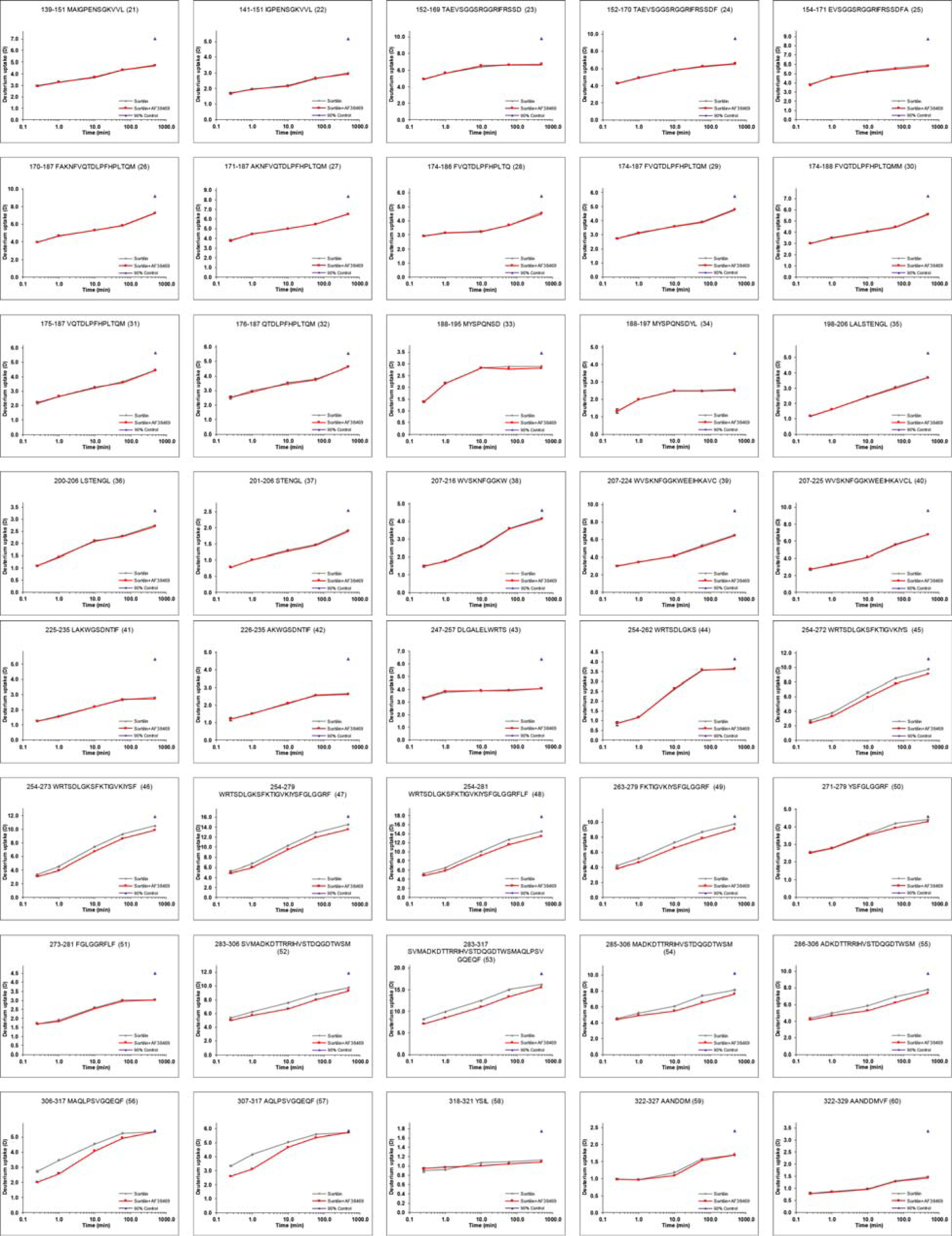

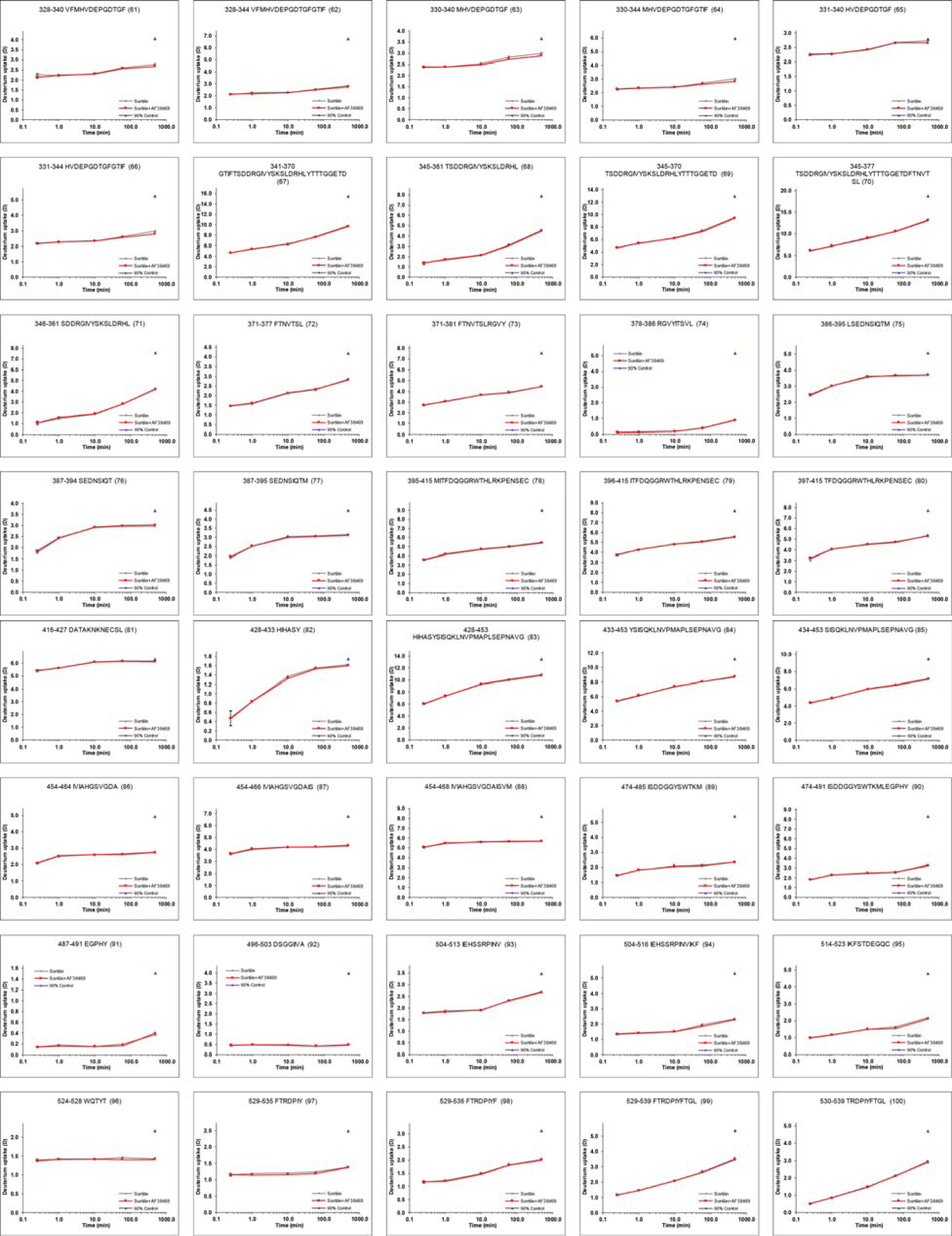

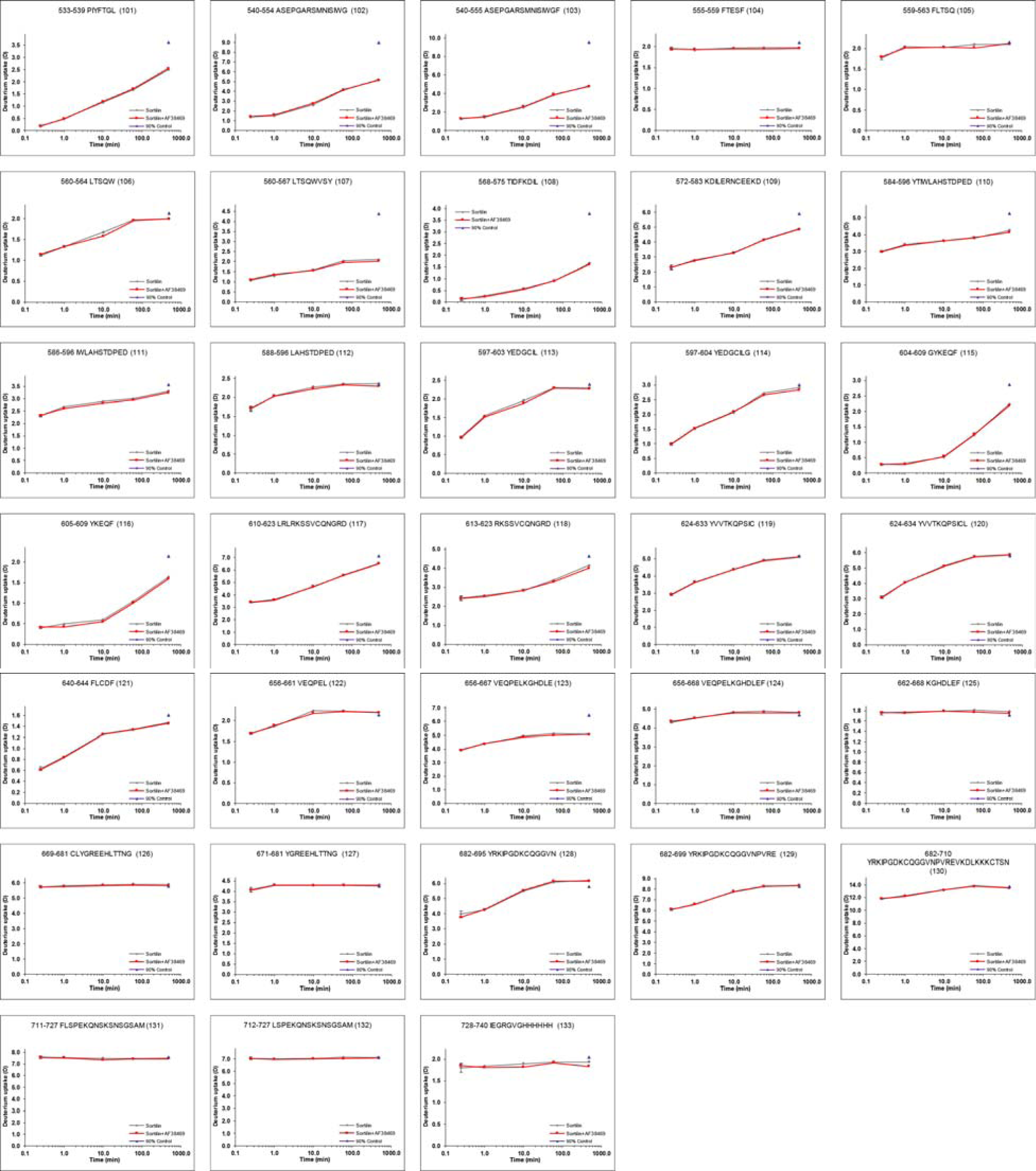

HDX plots of Sortilin in presence of proSort. Absolute deuterium incorporation is plotted as a function of time for Sortilin (gray lines) and Sortilin in presence of proSort (red lines). Equilibrium labelled (90%) Sortilin control samples are plotted as filled purple triangles at the 8h time point. SD is plotted as error bars (are only slightly visible). (n=3 for the lmin and lh time points and the equilibrium labelled sample. n=l for the 15s, lOmin and 8h time points).

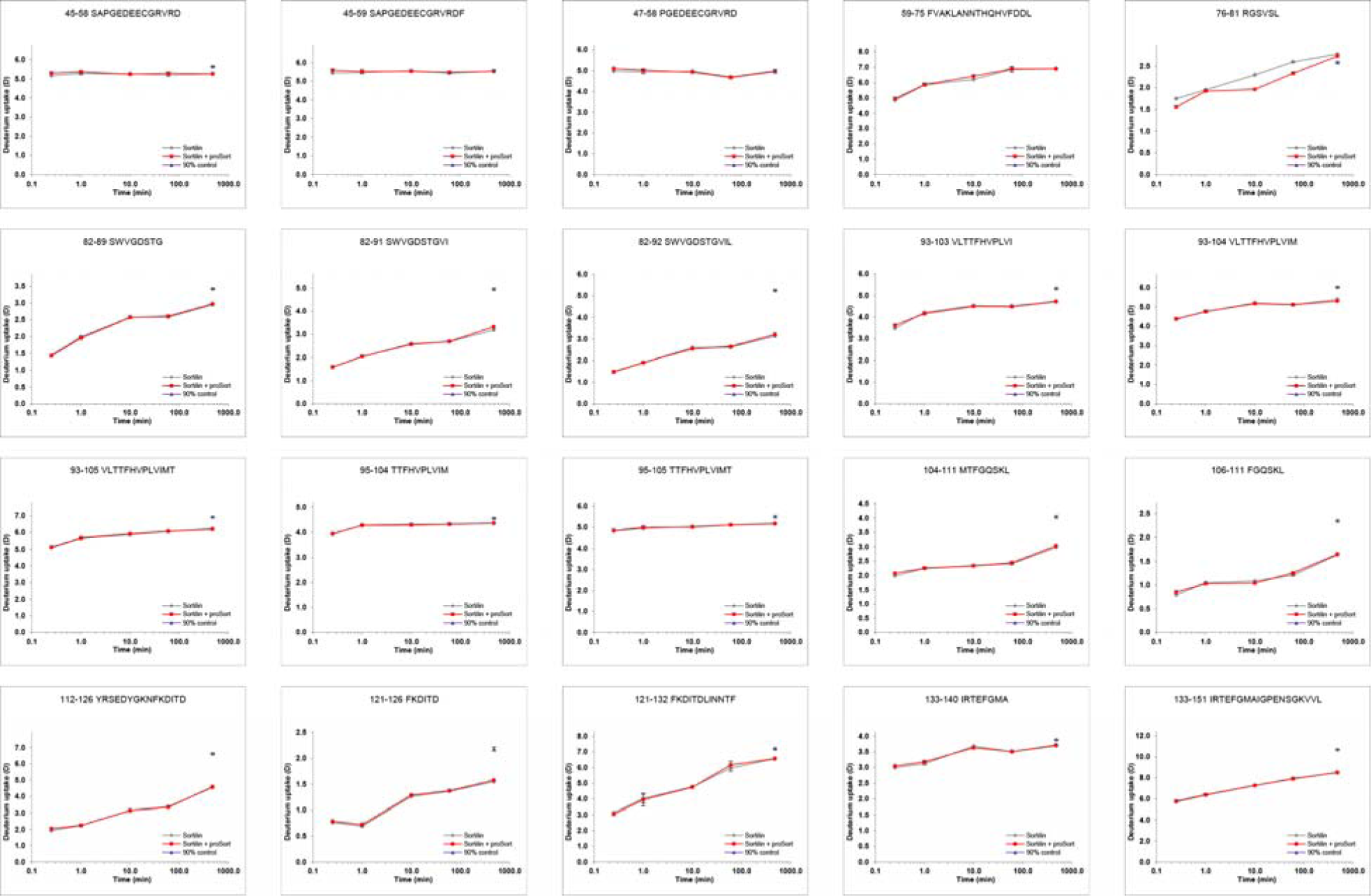

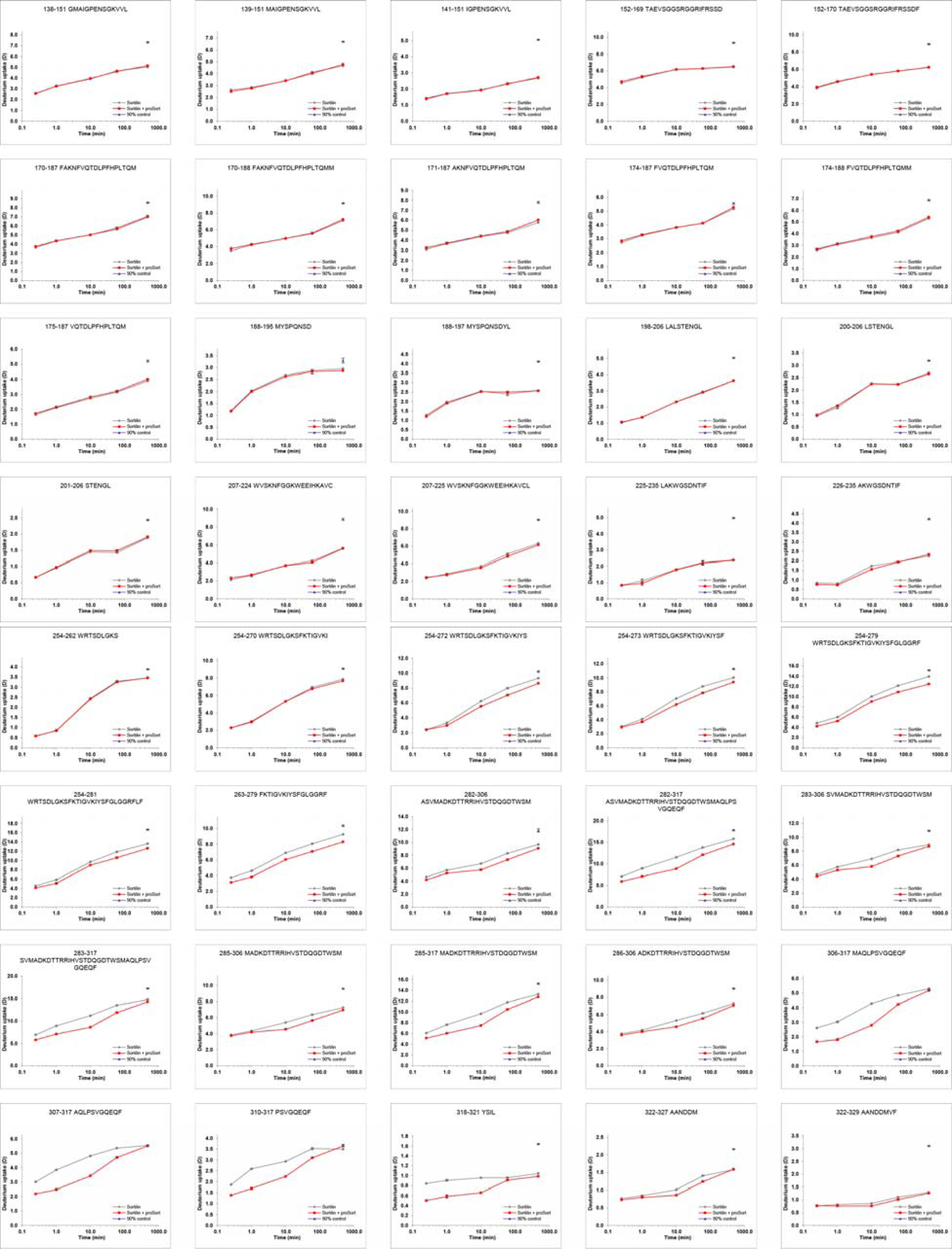

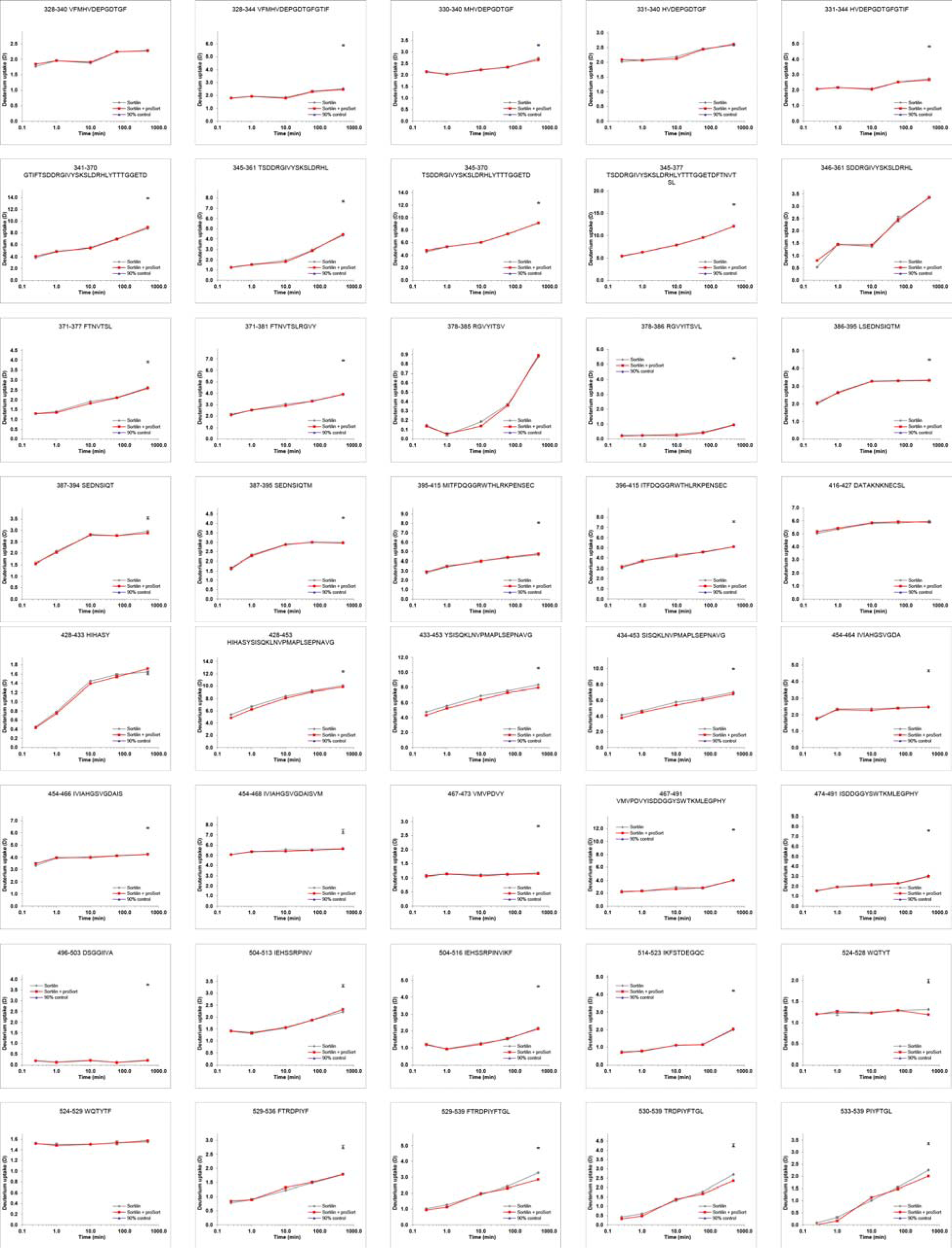

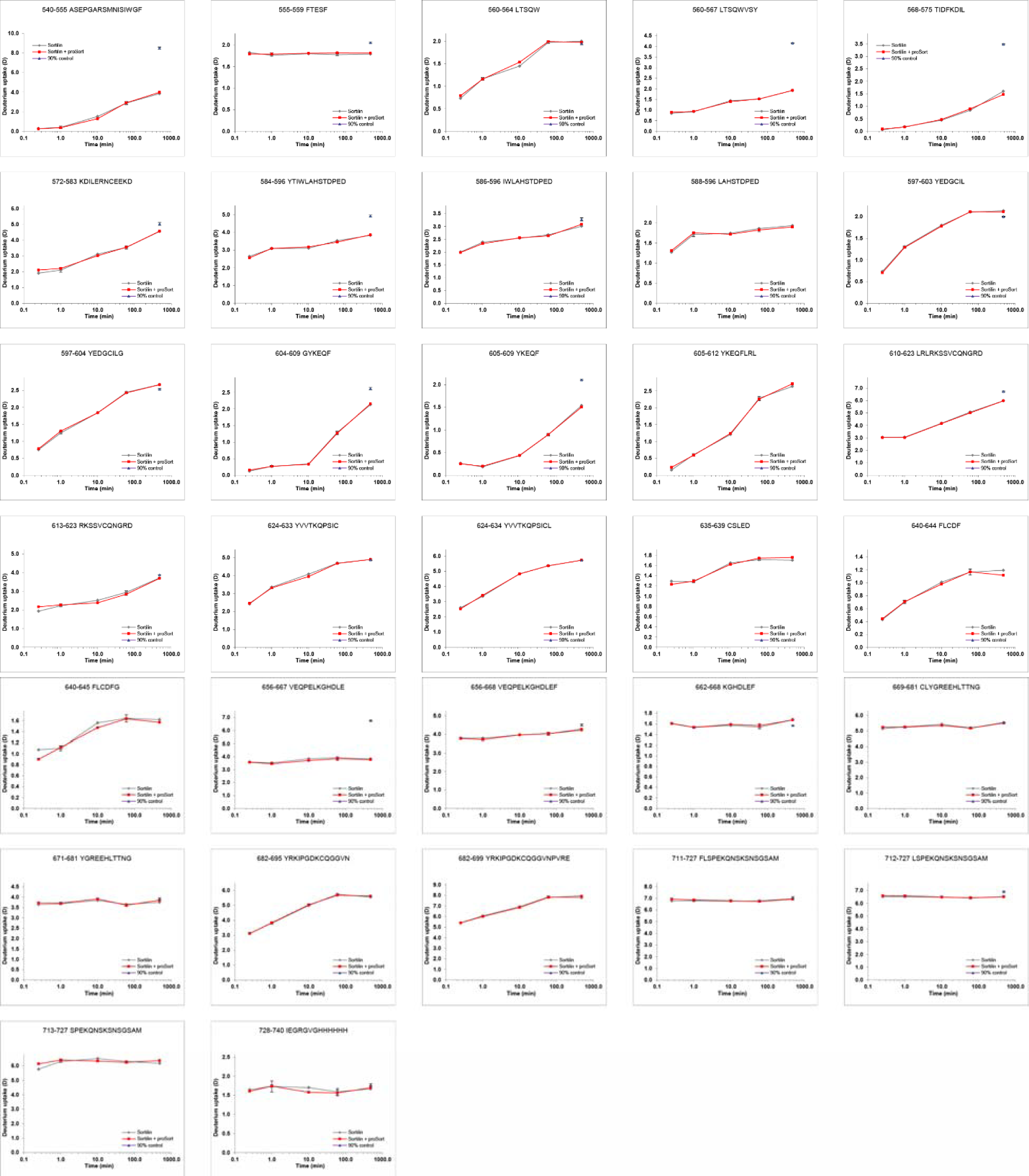

HDX plots of proSort in presence of Sortilin. Absolute deuterium incorporation is plotted as a function of time for proSort (gray lines) and proSort in presence of Sortilin (red lines). Equilibrium labelled (90%) proSort control samples are plotted as filled purple triangles circles at the lh time point. SD is plotted as error bars (are only slightly visible). (n=l for all time points, except the 10s time point, where n=4)

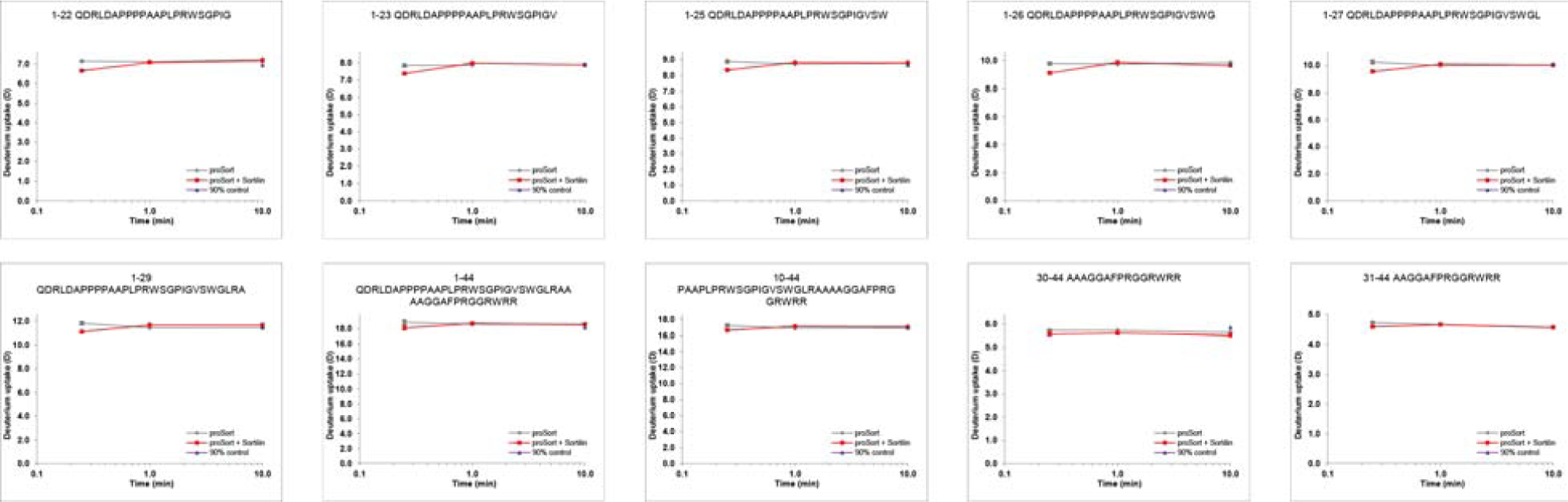

HDX plots of Sortilin in presence of proNGF. Absolute deuterium incorporation is plotted as a function of time for Sortilin (gray lines) and Sortilin in presence of proNGF (red lines). Equilibrium labelled (90%) Sortilin control samples are plotted as filled purple triangles at the 8h time point. SD is plotted as error bars (are only slightly visible). (n=3 for the 15s, lmin and lOmin time points and the equilibrium labelled sample. n=l for the lh and 8h time points).

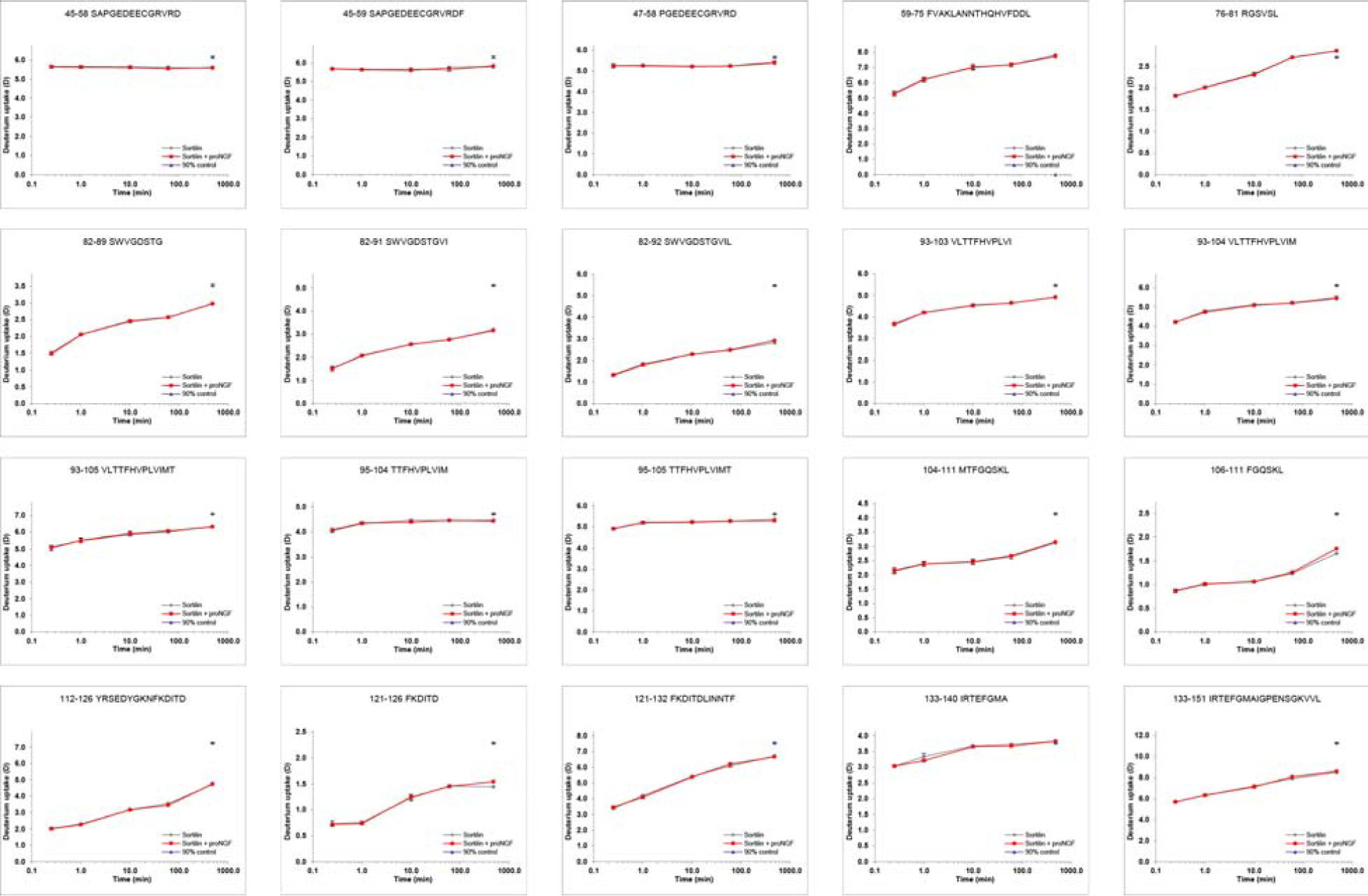

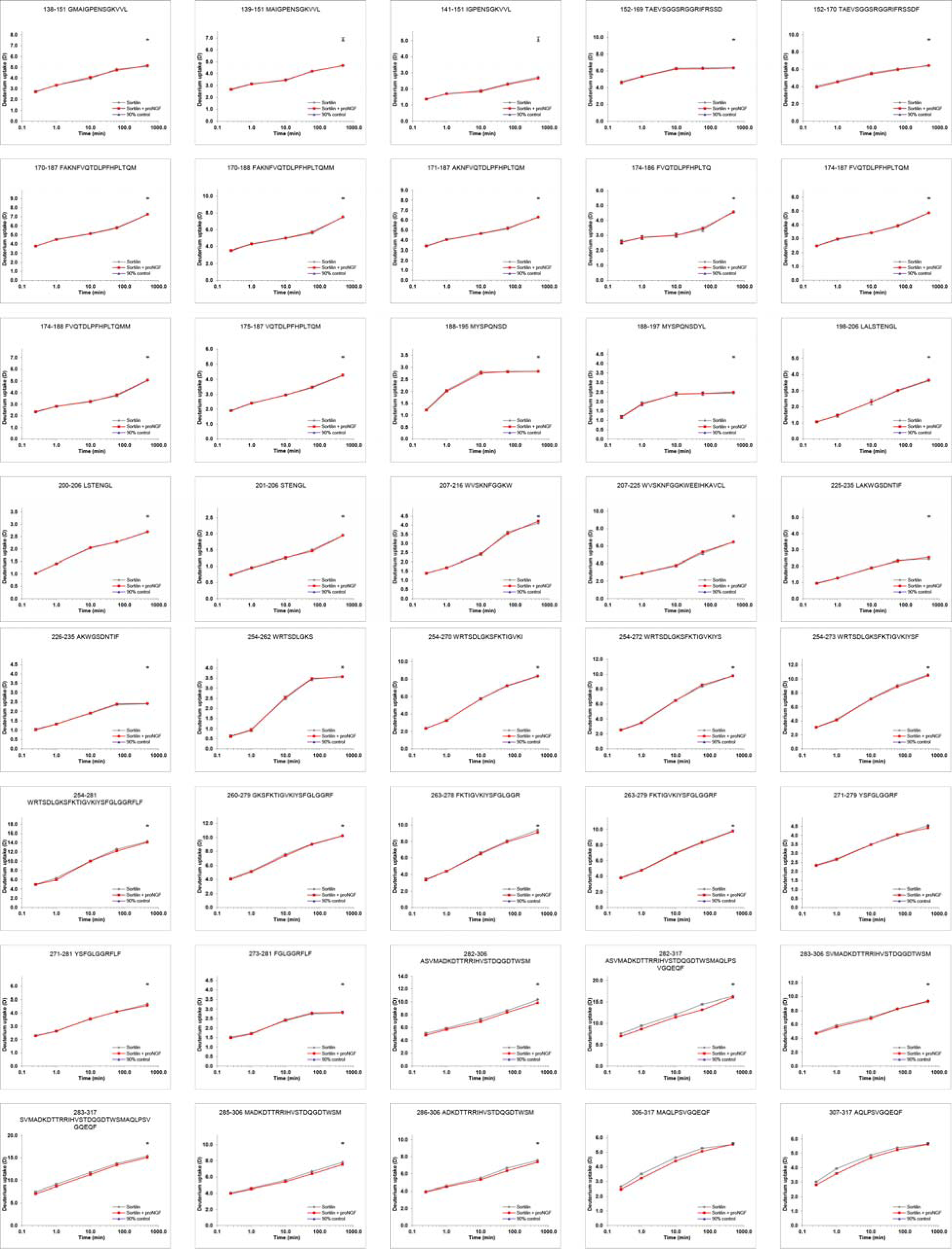

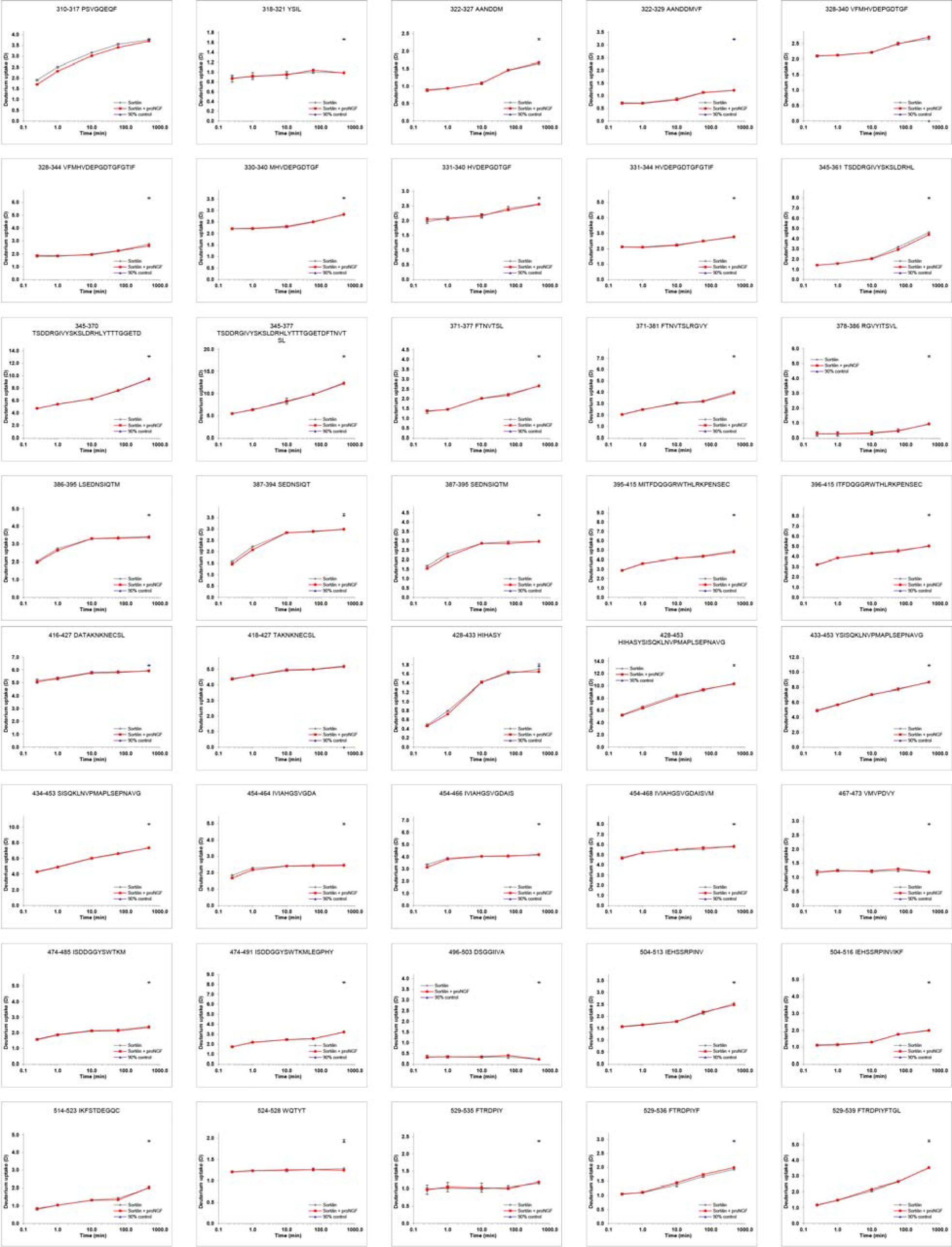

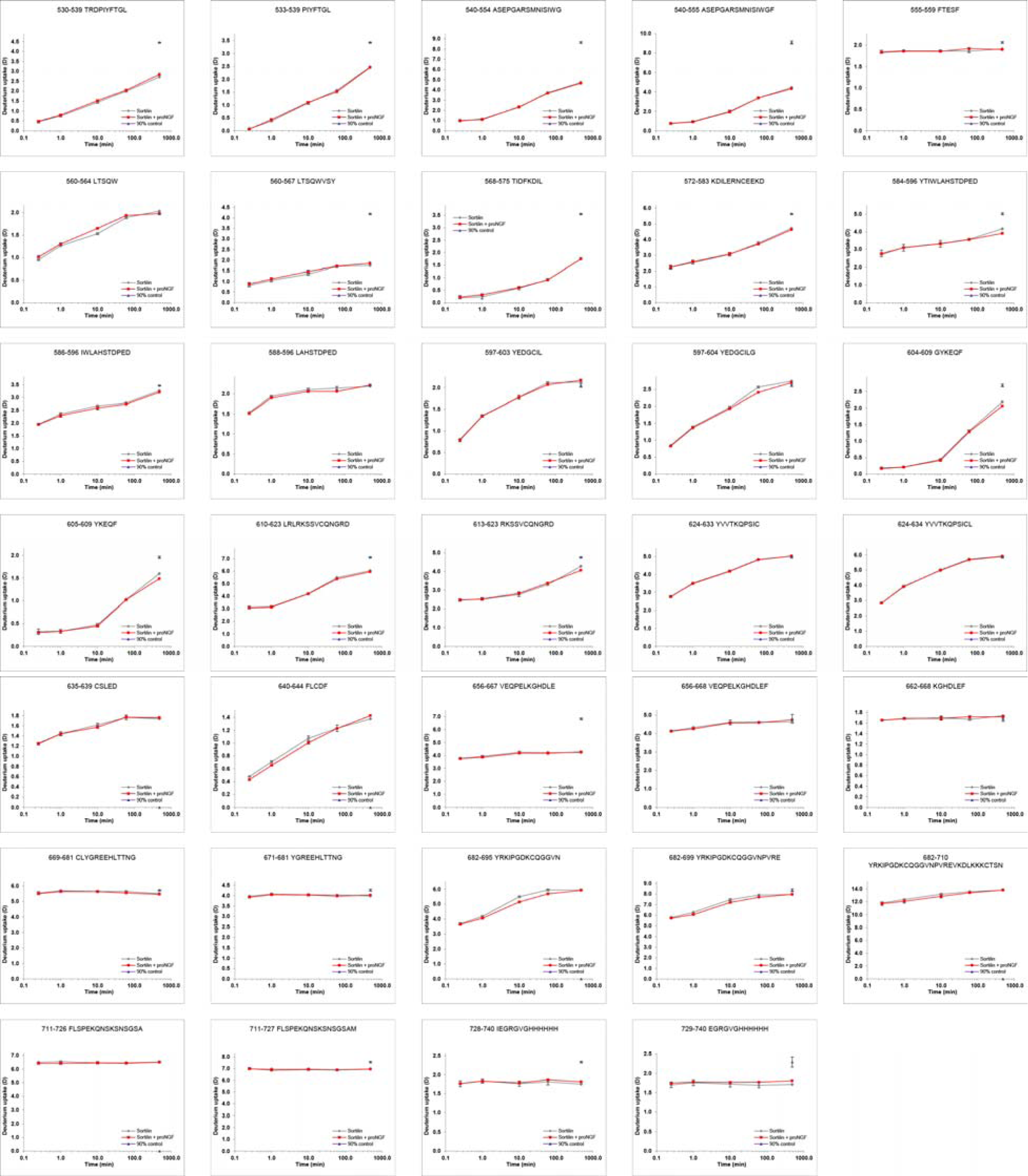

HDX plots of Sortilin in presence of GSTpro. Absolute deuterium incorporation is plotted as a function of time for Sortilin (gray lines) and Sortilin in presence of GSTpro (red lines). Equilibrium labelled (90%) Sortilin control samples are plotted as filled purple triangles at the 8h time point. SD is plotted as error bars (are only slightly visible). (n=3 for the 15s, lmin and 8h time points and the equilibrium labelled sample. n=l for the lOmin and lh time points).

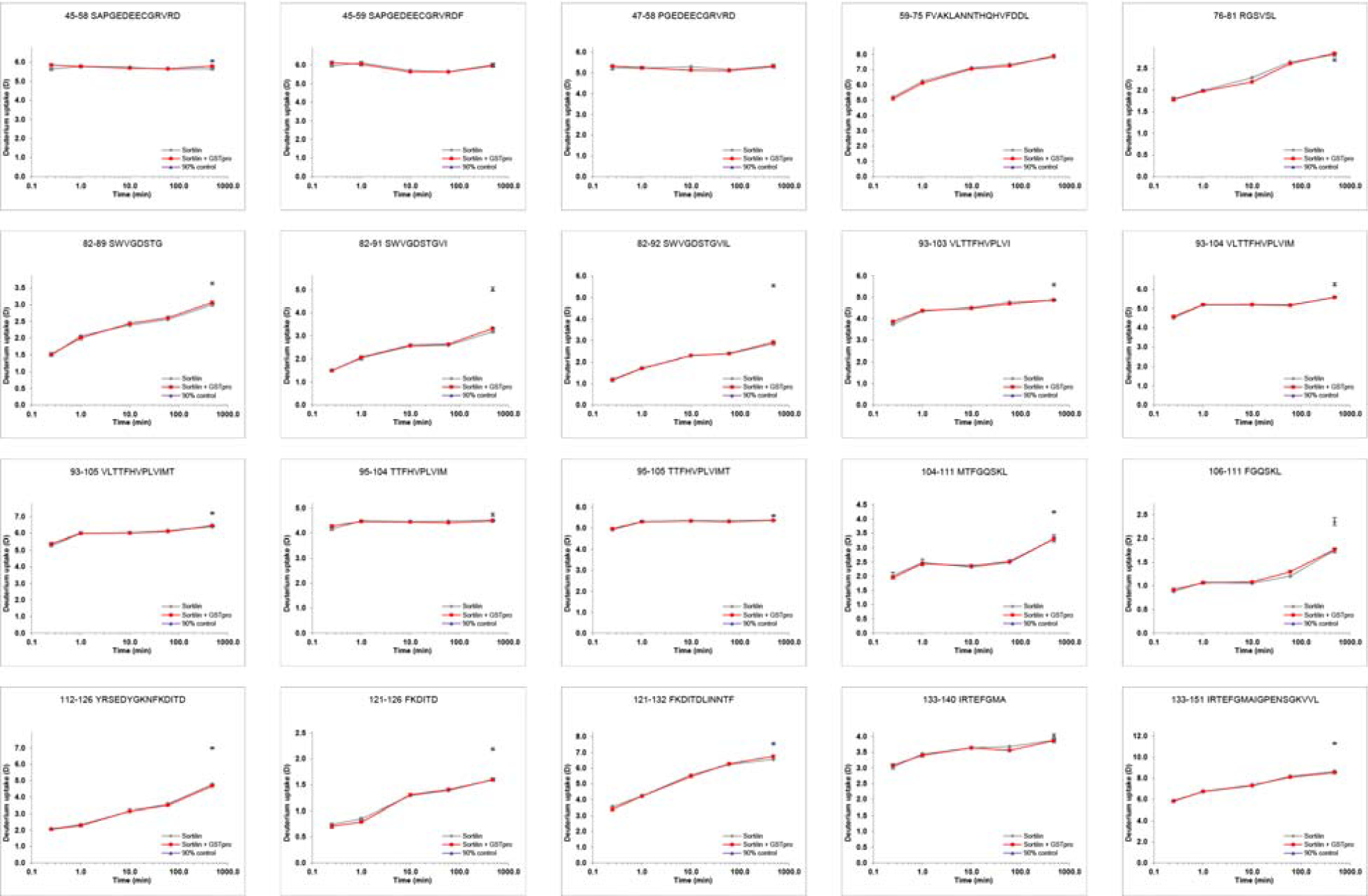

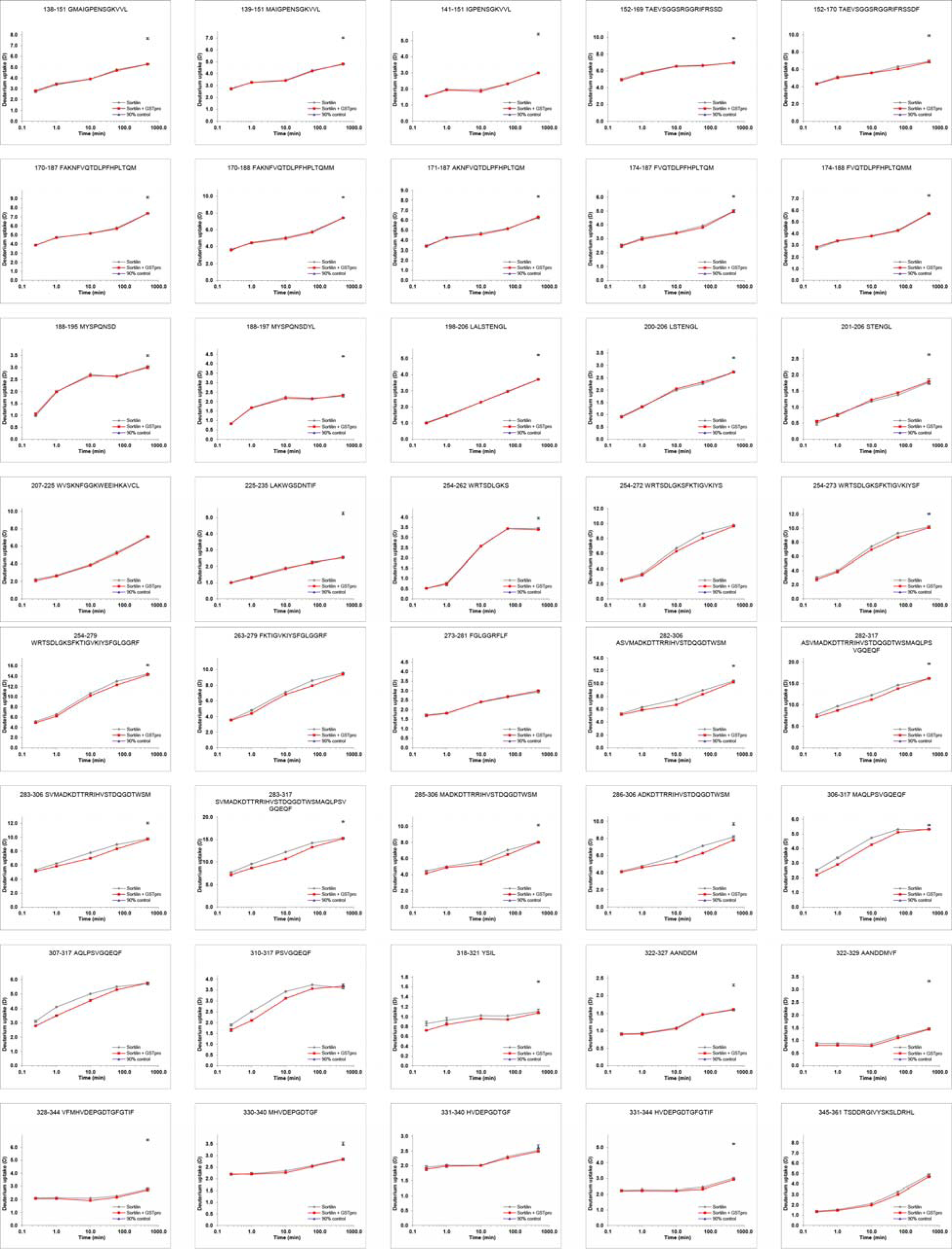

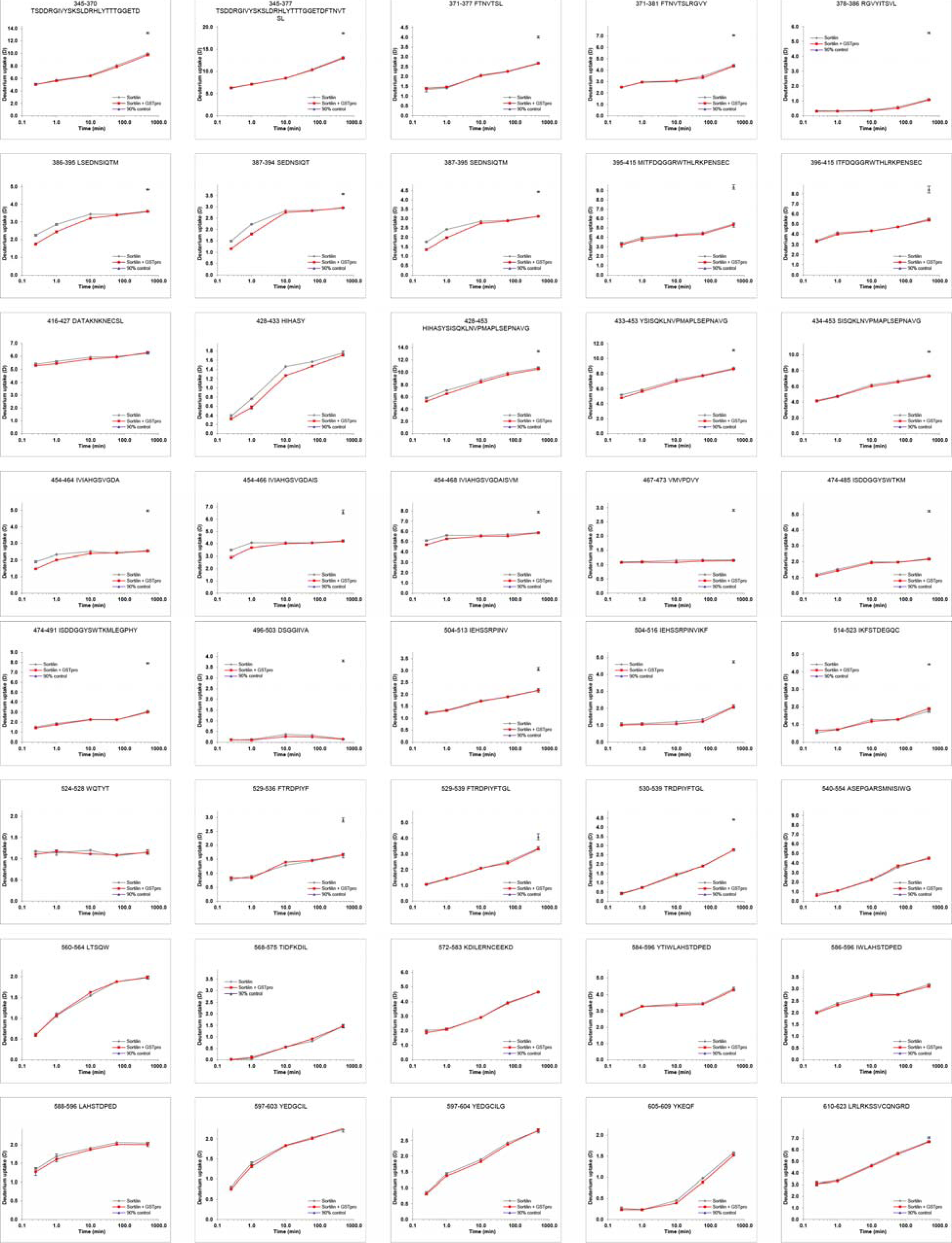

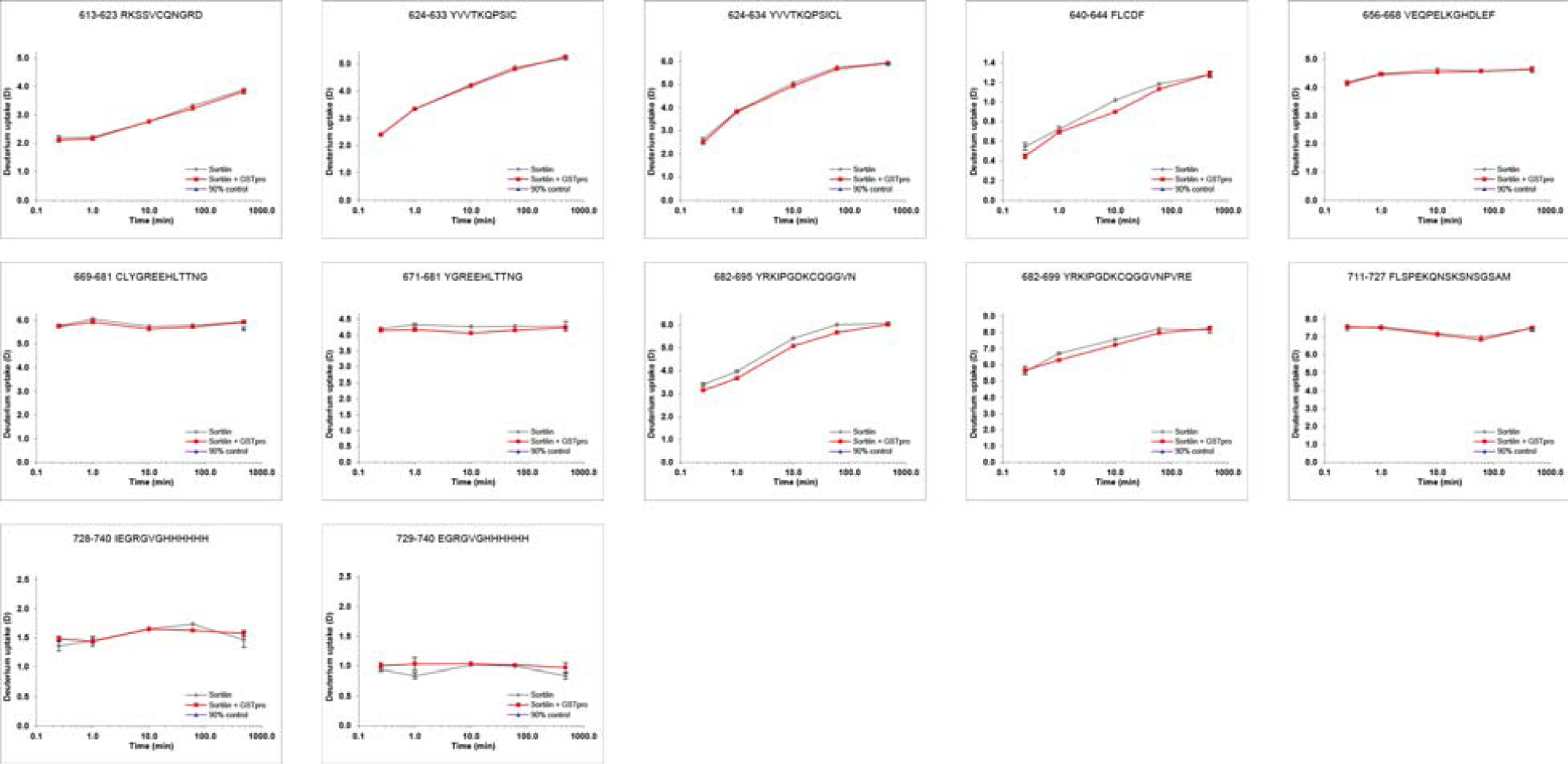

HDX plots of Sortilin in presence of GST. Absolute deuterium incorporation is plotted as a function of time for Sortilin (gray lines) and Sortilin in presence of GST (red lines). Equilibrium labelled (90%) Sortilin control samples are plotted as filled purple triangles at the 8h time point. SD is plotted as error bars (are only slightly visible). (n=3 for the lmin and lh time points and the equilibrium labelled sample. n=l for the 15s, lOmin and 8h time points).

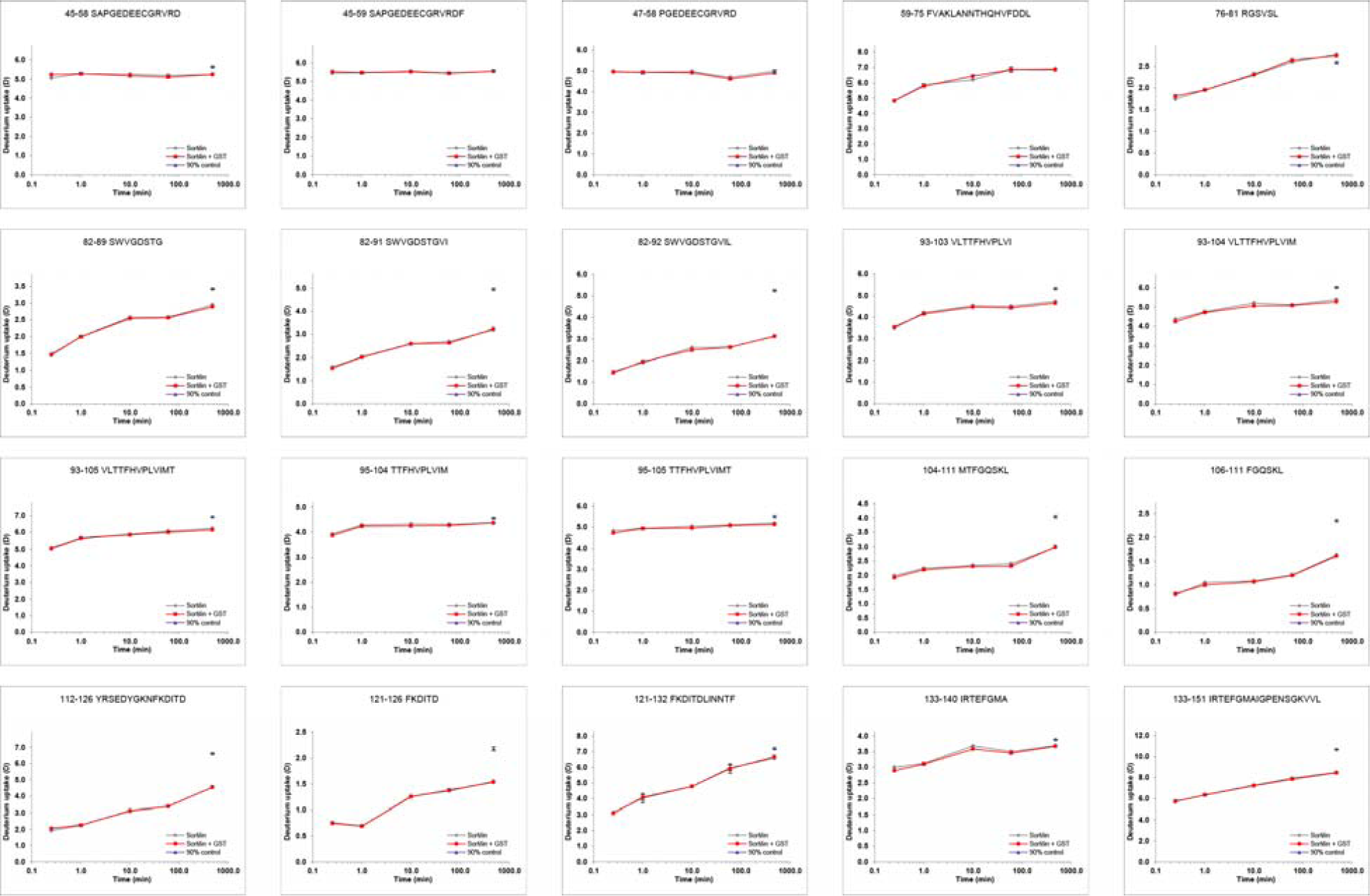

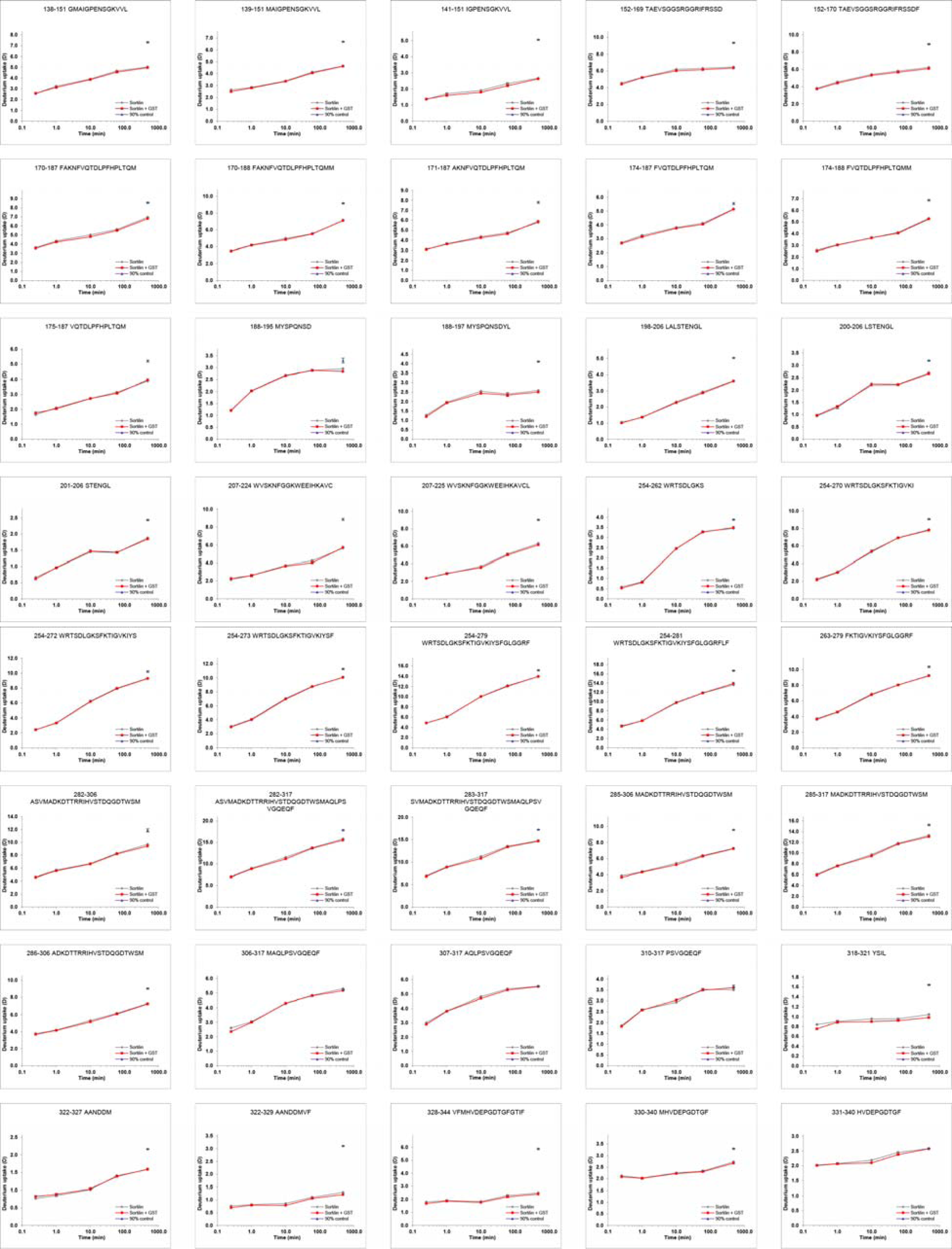

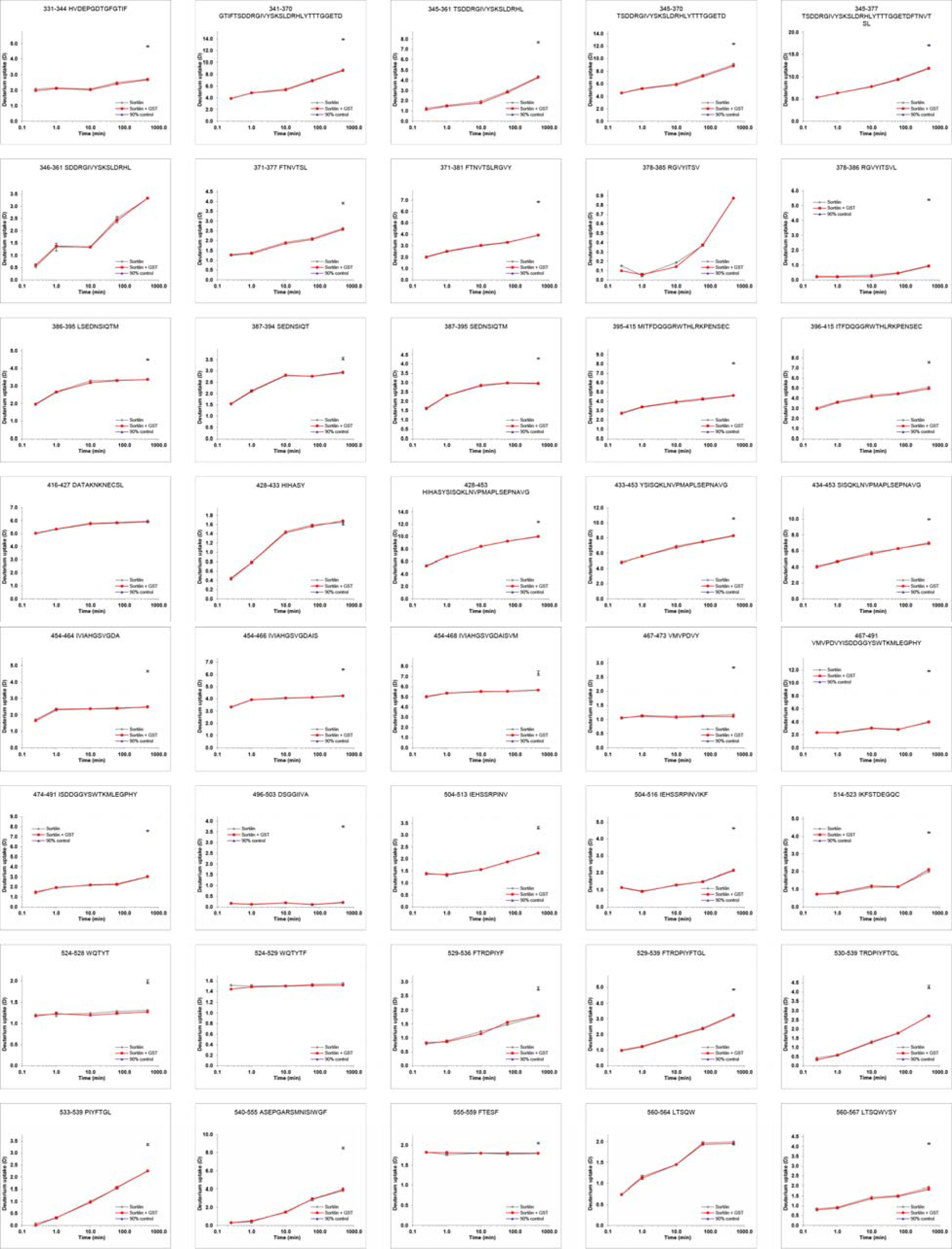

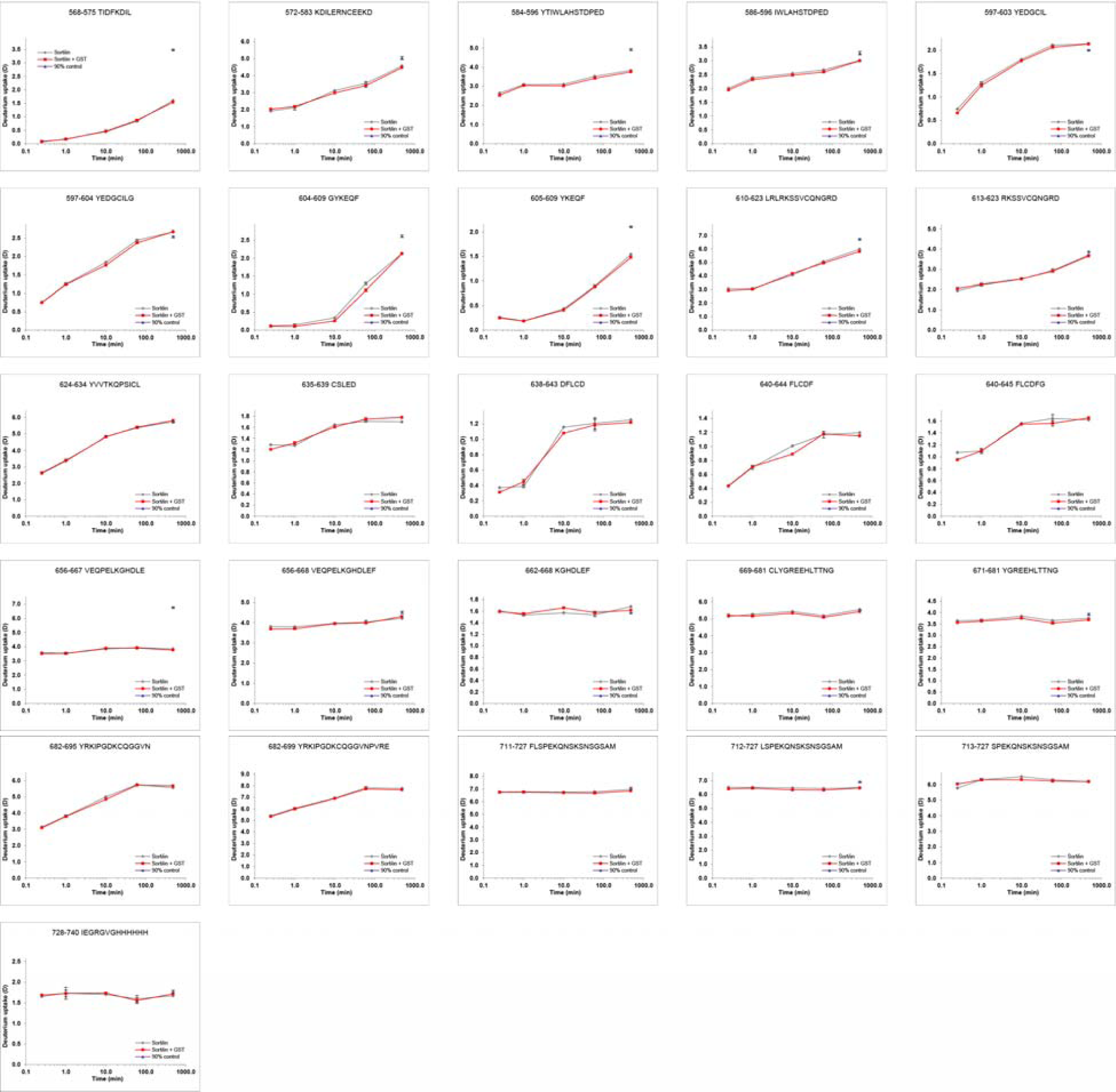

HDX plots of Sortilin in presence of NGF. Absolute deuterium incorporation is plotted as a function of time for Sortilin (gray lines) and Sortilin in presence of NGF (red lines). Equilibrium labelled (90%) Sortilin control samples are plotted as filled purple triangles at the 8h time point. SD is plotted as error bars (are only slightly visible). (n=3 for the lmin, lh and 8h time points and the equilibrium labelled sample. n=1 for the 15s, 10min and 8h time points).

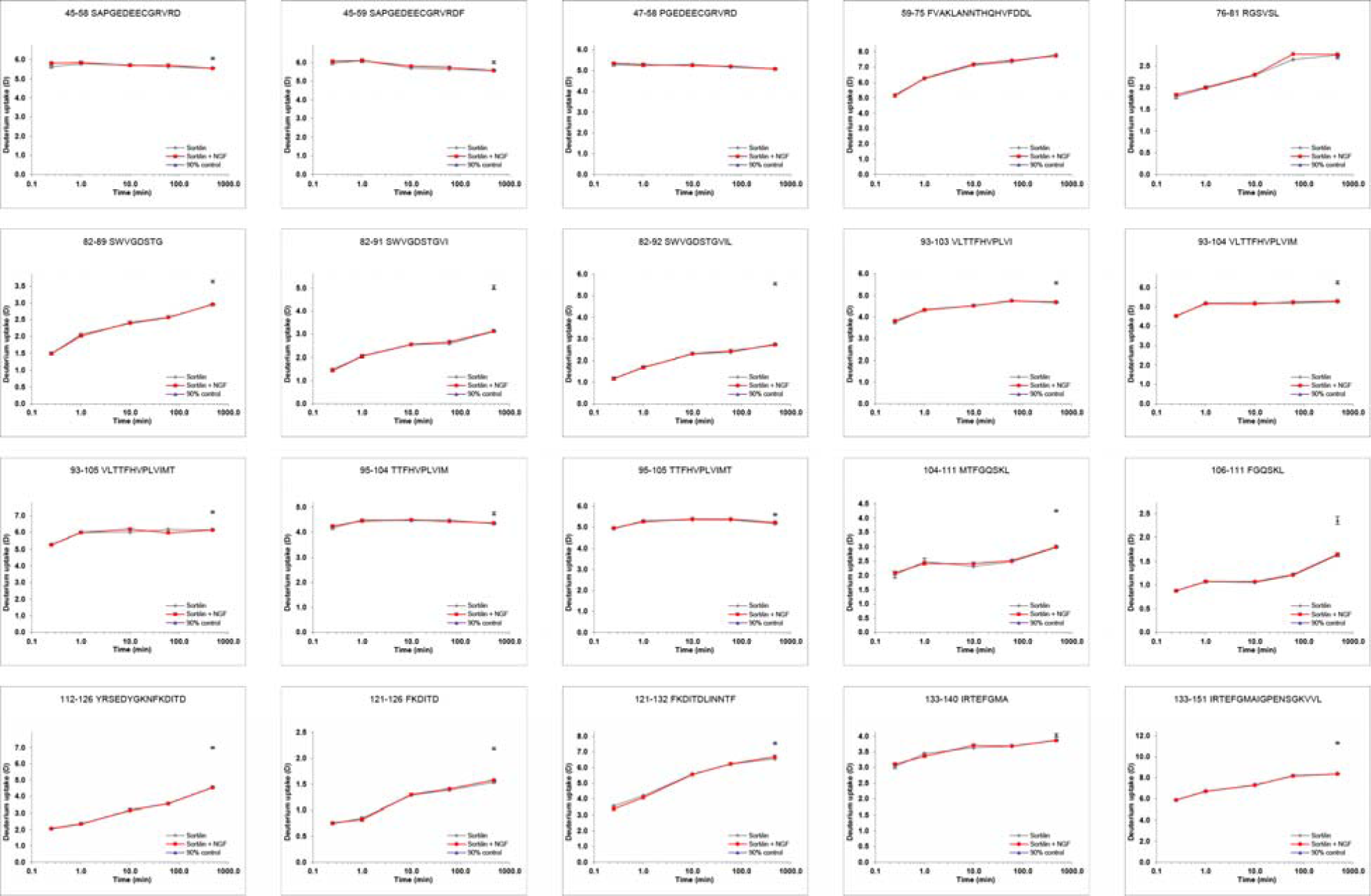

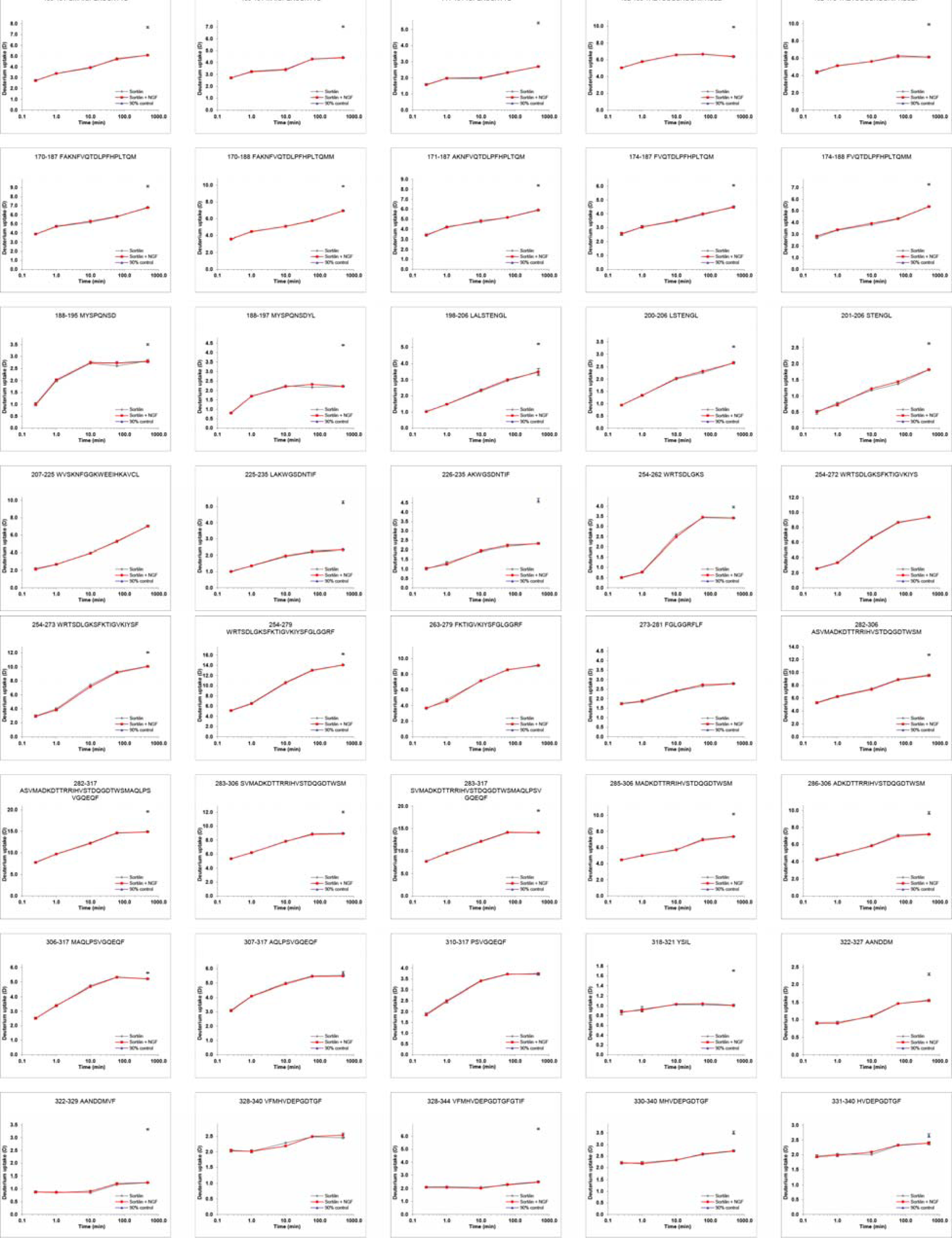

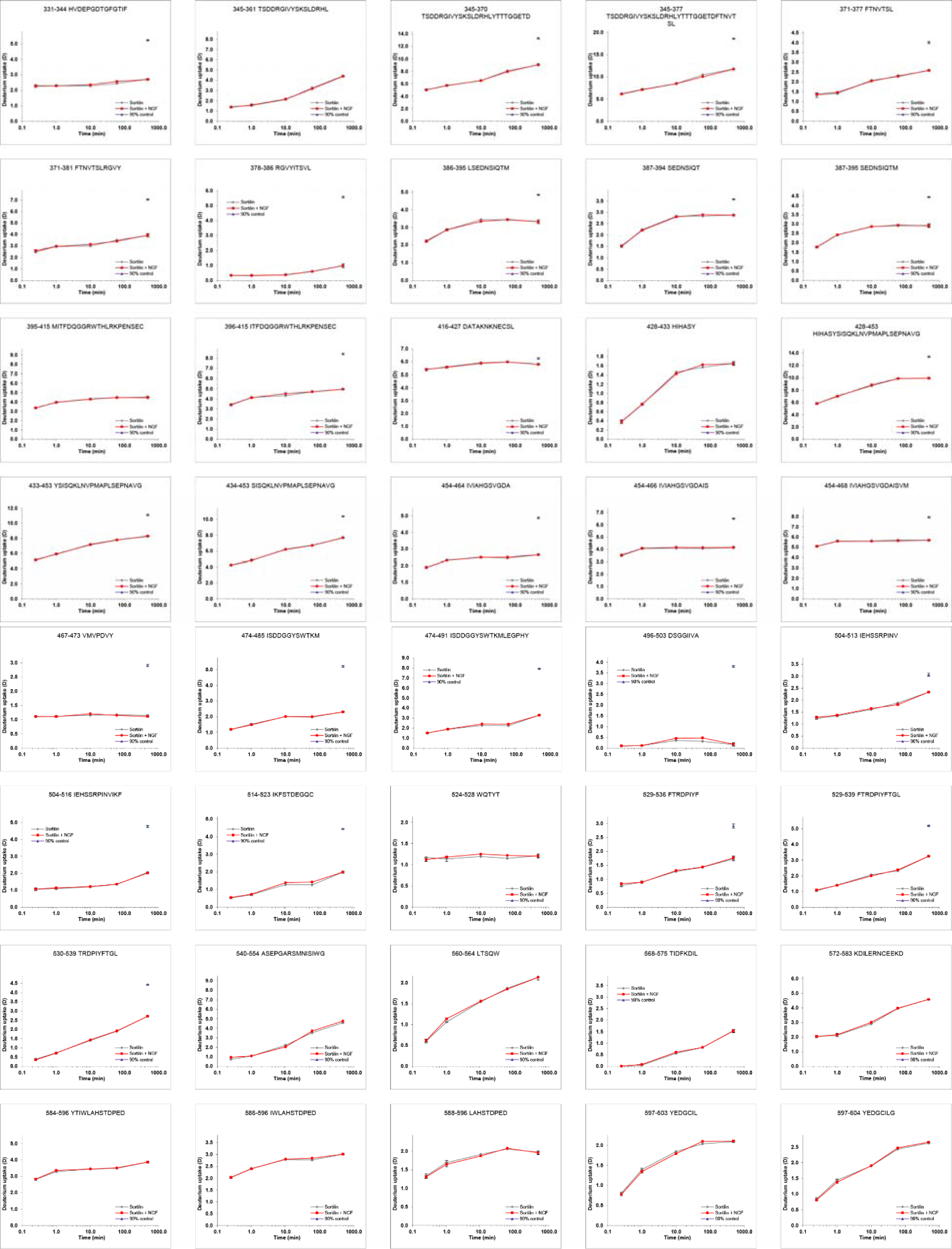

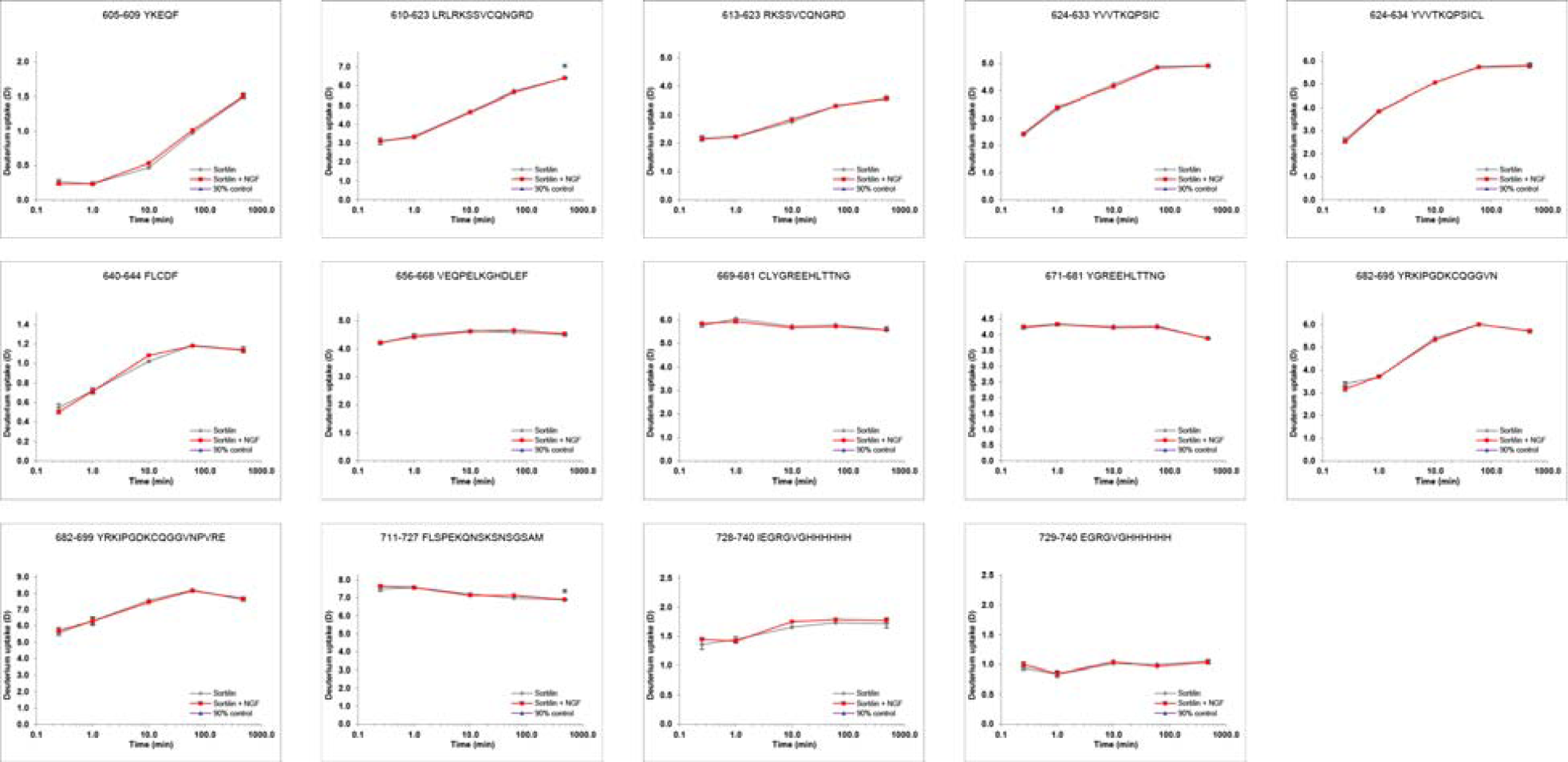

HDX plots of Sortilin in presence of PGRN. Absolute deuterium incorporation is plotted as a function of time for Sortilin (gray lines) and Sortilin in presence of NGF (red lines). Equilibrium labelled (90%) Sortilin control samples are plotted as filled purple triangles at the 8h time point. SD is plotted as error bars (are only slightly visible). (n=3 for the 15s, lmin and lOmin time points and the equilibrium labelled sample. n=l for the lh and 8h time points).

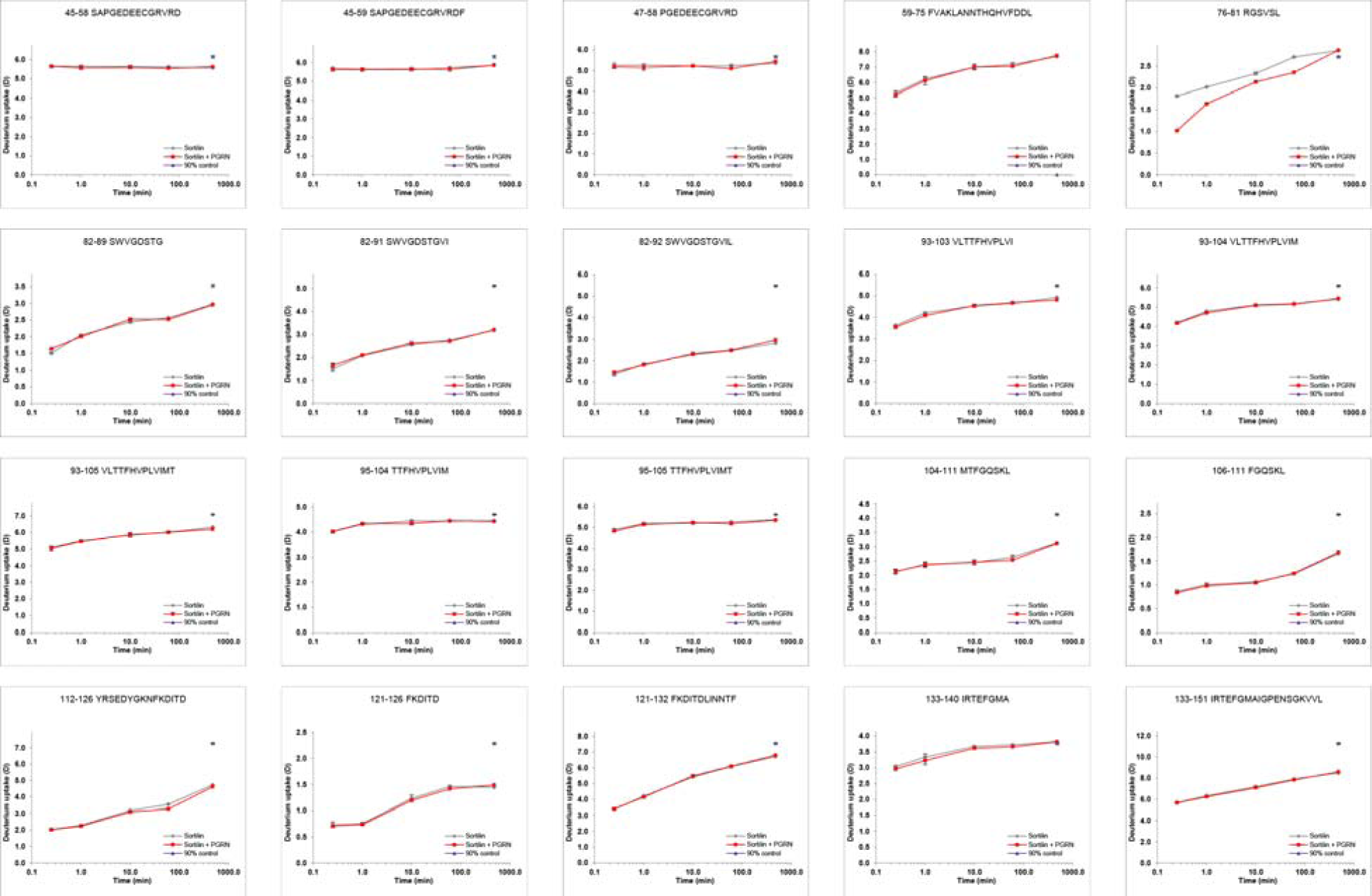

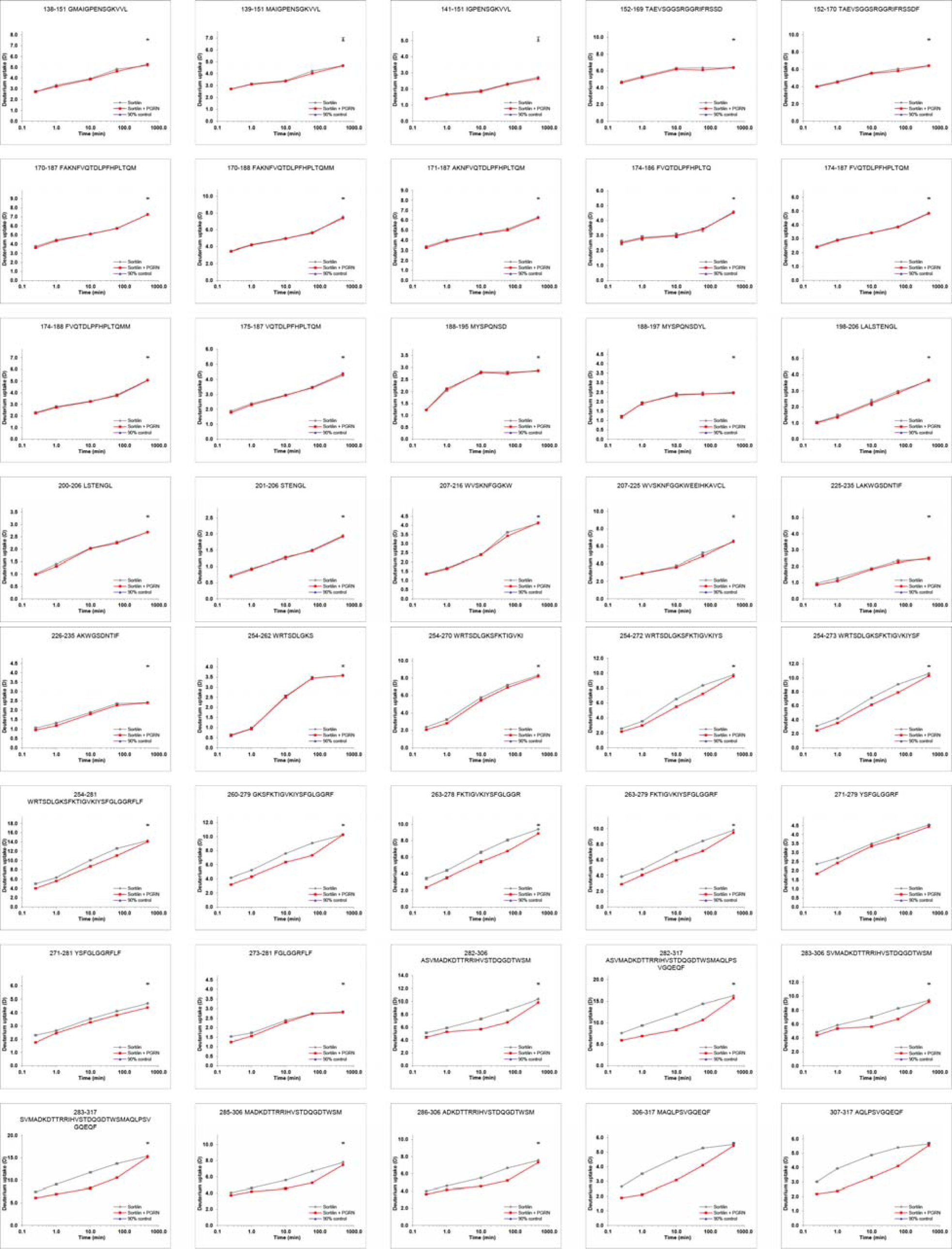

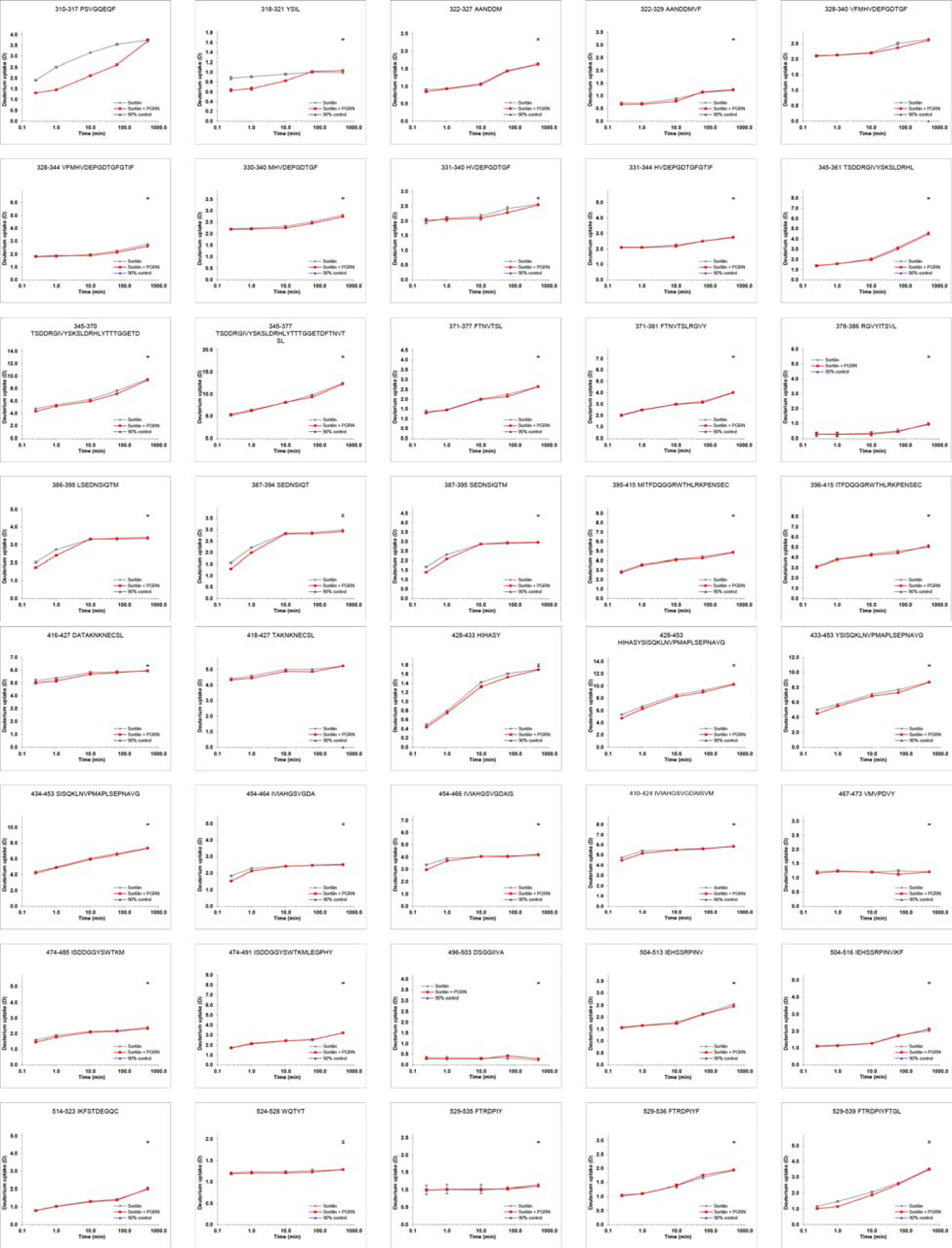

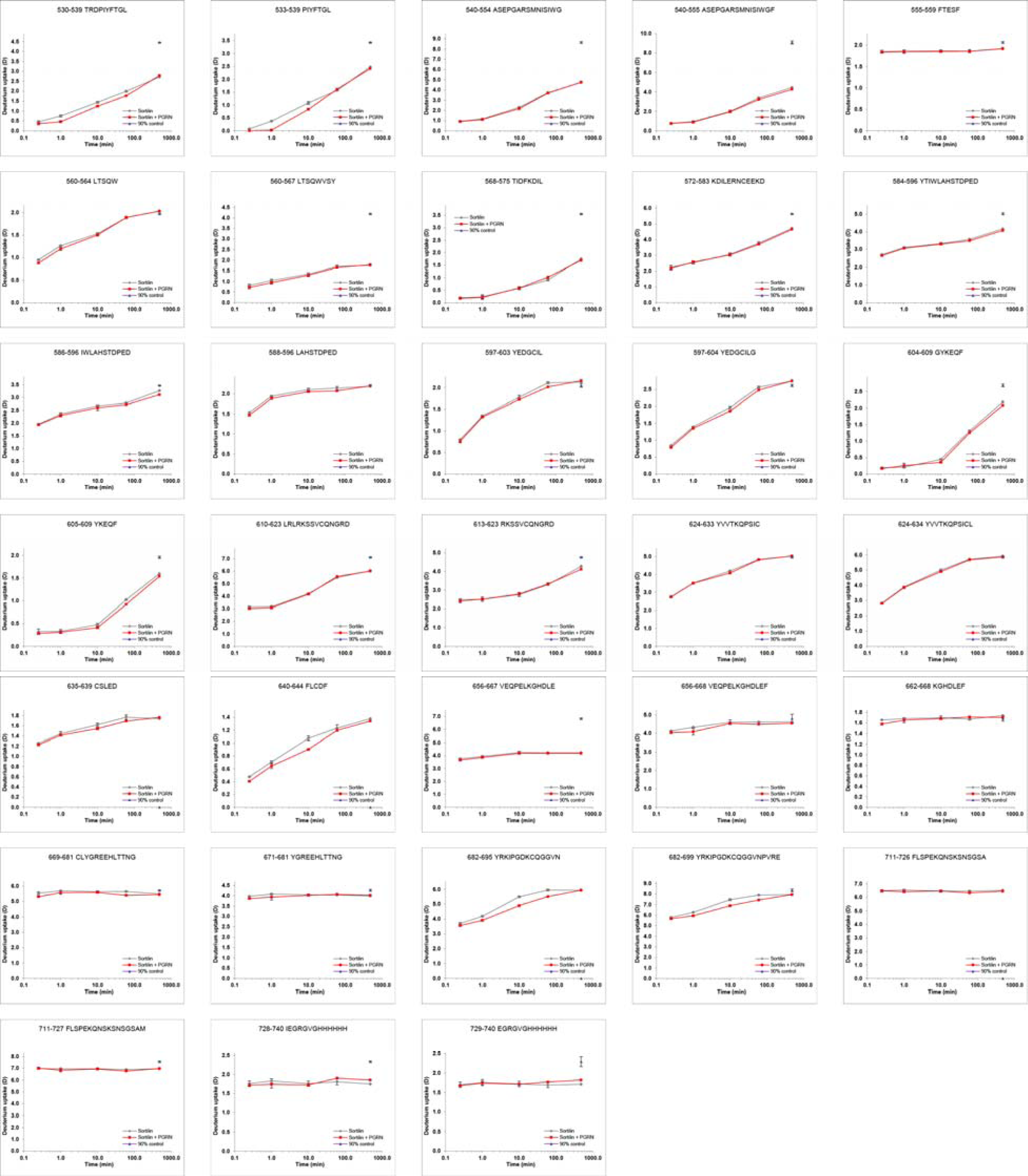

HDX plots of PGRN in presence of Sortilin. Absolute deuterium incorporation is plotted as a function of time for PGRN (gray lines) and PGRN in presence of Sortilin (red lines). Equilibrium labelled (90%) proSort control samples are plotted as filled purple triangles circles at the 8h time point. SD is plotted as error bars (are only slightly visible). (n=3 for the 15smin and lmin time points. n=l for the lOmin, lh and 3h time points and the equilibrium labelled sample).

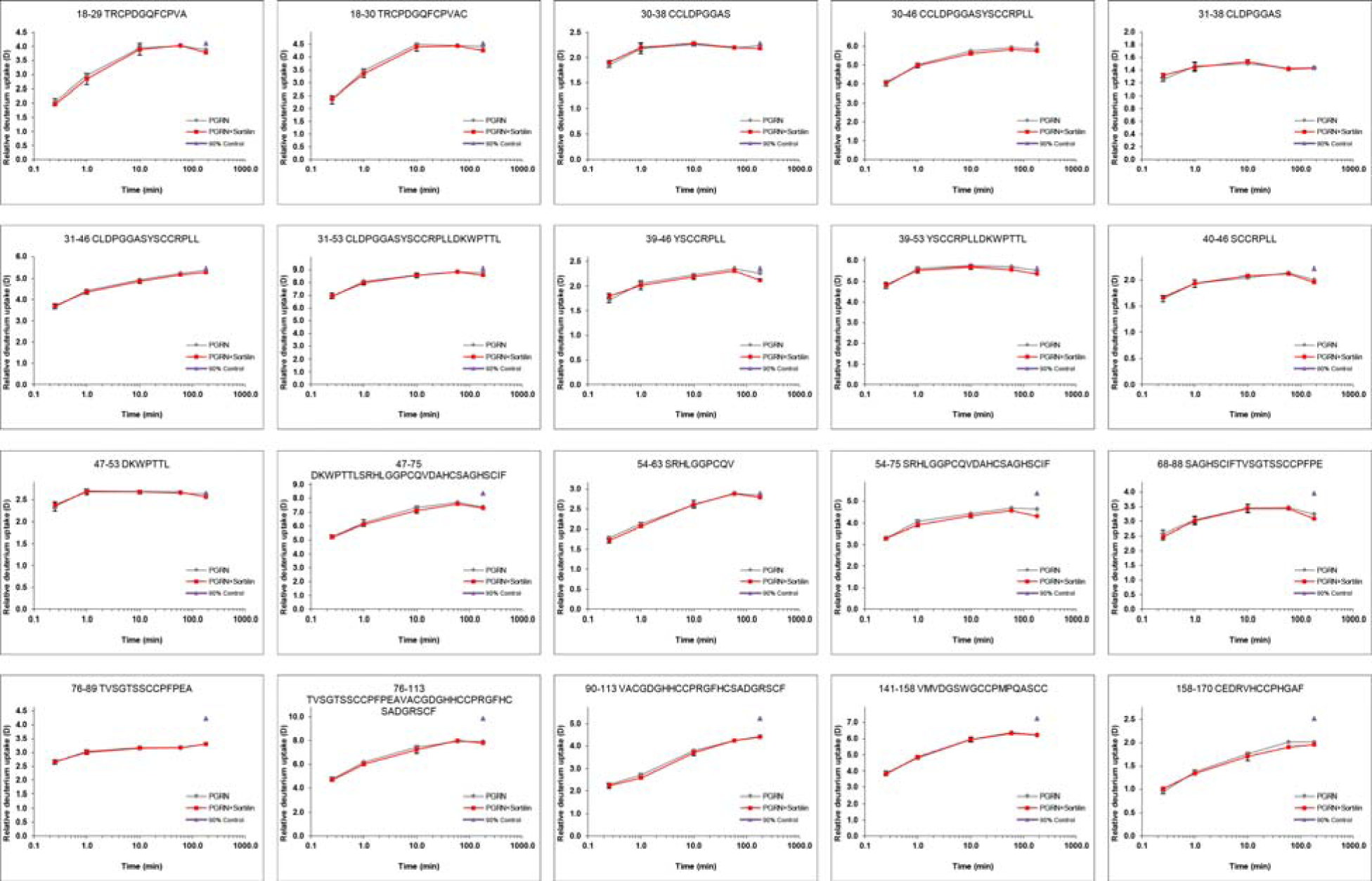

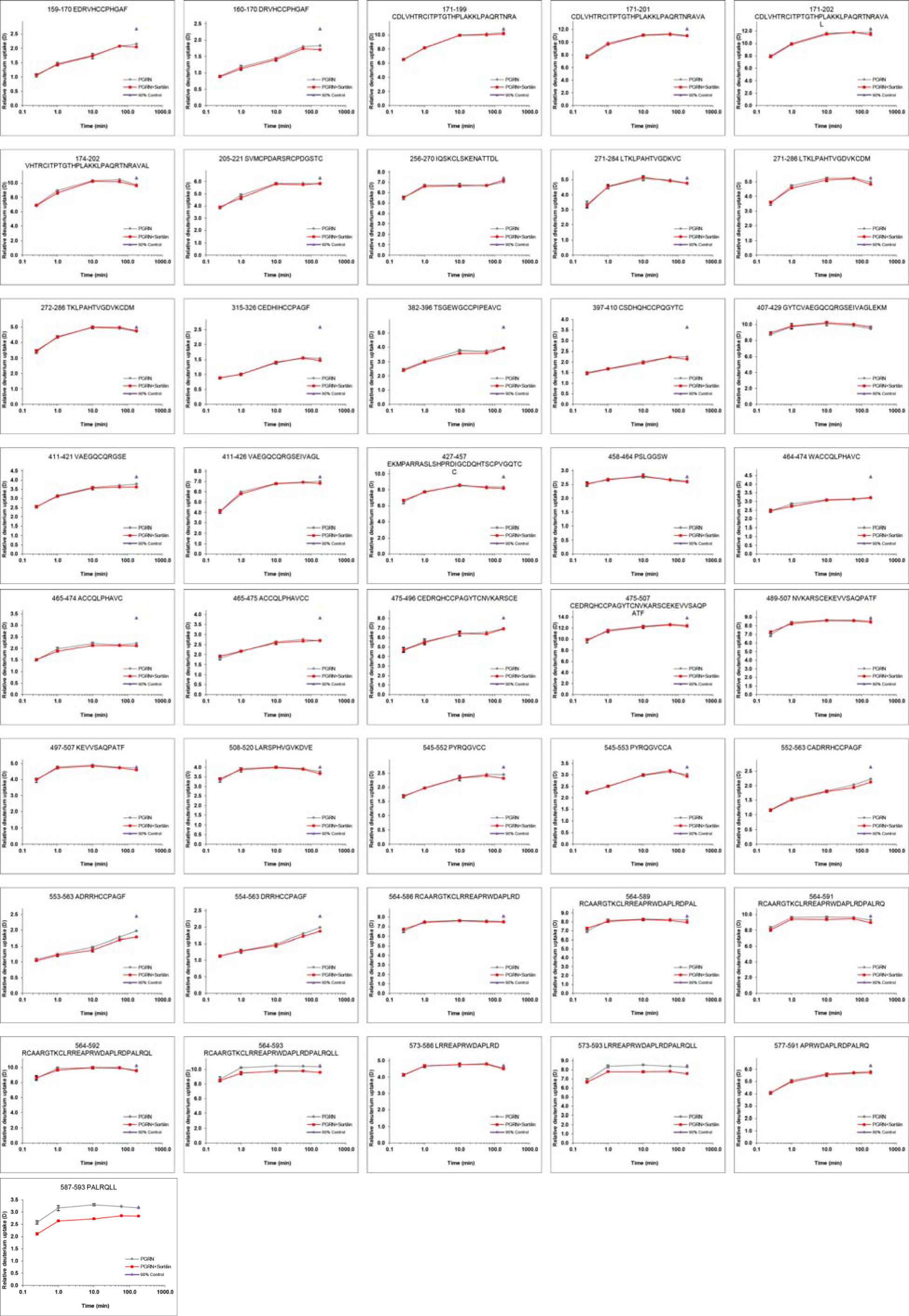

